# High-resolution, genotype-free mapping of genetic variation with CRI-SPA-Map

**DOI:** 10.1101/2025.09.16.676640

**Authors:** Sheila Lutz, Megan Lawler, Samuel Amidon, Frank W. Albert

## Abstract

Genetic variation within species shapes phenotypes, but identifying the specific genes and variants that cause phenotypic differences is costly and challenging. Here, we introduce CRI-SPA-Map, a genetic mapping strategy combining CRISPR-Cas9 genome engineering, selective ploidy ablation (SPA), and high-throughput phenotyping for precise genetic mapping with or without genotyping in the yeast *Saccharomyces cerevisiae*. In CRI-SPA-Map, a donor strain carrying SPA machinery is mated to a genetically different recipient strain harboring a genome-integrated selectable cassette. In the resulting diploid, CRISPR-Cas9 cuts the cassette for replacement with DNA from the homologous donor chromosome. Donor chromosomes are then removed using SPA to yield haploid recombinant strains. To establish CRI-SPA-Map, we mated a W303 SPA strain to 92 strains from the BY4742 yeast knockout collection that carry gene deletion cassettes on the left arm of chromosome XIV and created 1,451 recombinant isolates. Whole-genome sequencing verified that deletion cassette replacement introduced short donor DNA tracts of variable length, resulting in a finely recombined mapping population. Using only the known location of the gene deletions, which marks where donor DNA is introduced, we identified a 6.5 kb-region shaping yeast growth. Further dissection of this region pinpointed two causal variants in two genes, *MKT1* and *SAL1*. Engineering these variants alone and in combination revealed gene-by-environment interactions at both genes, as well as epistatic interactions between them that were in turn dependent on the environment. CRI-SPA-Map is a cost-effective strategy for creating high-resolution recombinant panels of yeast strains for identifying the genetic basis of phenotypic variation.

## Introduction

Most phenotypic traits, ranging from morphological, physiological, and molecular quantities to the risk for common disease vary continuously among individuals in a population. Studies in numerous species have shown that quantitative traits have a complex genetic basis, in which they are shaped by DNA variants at dozens to thousands of genes (Mackay and Anholt 2024). Each causal variant typically has a small effect on the trait, but due to the large number of causal variants, they contribute substantially to trait variation.

Two main experimental designs are used to dissect genetic variation in complex traits. Genetic mapping by linkage analysis involves crossing genetically different individuals to generate recombinant progeny, which are subsequently phenotyped for the trait of interest (Lynch and Walsh 1998). Genomic regions harboring causal alleles are identified through linkage with genetic markers determined by genome sequencing or other methods. Linkage analysis in crosses of two inbred parents can detect the effects of all variants that differ between the parents with equal statistical power, irrespective of their frequency in the wider population.

However, the size of the regions identified by linkage mapping depends on the frequency of meiotic recombination and the number of recombinant progeny that can be both phenotyped and genotyped. Genome-wide association studies (GWAS) rely on past recombination events that have reshuffled genetic variation in natural populations of unrelated individuals (Visscher et al. 2012). GWAS can assay more variation than biparental crosses but has low power for variants that are rare in the population. GWAS is also prone to confounding by population structure (Peter et al. 2022) and by non-genetic, environmental factors (Young et al. 2019). The regions identified by linkage mapping and GWAS typically contain multiple variants and genes, making the identification of causal variants and mechanisms challenging (Lappalainen et al. 2024; Mackay and Anholt 2024). Crucially, both linkage mapping and GWAS require that all individuals in the study are genotyped. Together, these reasons render the identification of individual genes and variants whose variation shapes complex phenotypes challenging.

The yeast *Saccharomyces cerevisiae* is a key model for the dissection of genetically complex traits (Liti and Louis 2012). The uniformly high meiotic recombination rate along the genome coupled with the ease with which large panels of strains can be generated has led to powerful applications of linkage analysis. Prominent efforts have included panels of thousands of progeny from a biparental cross (Nguyen Ba et al. 2022; Bloom et al. 2015), crosses that spanned multiple generations for additional recombination (She and Jarosz 2018; Cubillos et al. 2013), the use of large pools of single, recombined cells (Ehrenreich et al. 2010; Albert et al. 2014), and crosses among multiple parents that cover much of the natural genetic variation in this species (Bloom et al. 2019). Successful GWAS studies have also been conducted (Peter et al. 2018). However, applications of these approaches in yeast still suffer from many of the same shortcomings as in other species. In particular, the identified regions remain wide due to the limited ability of meiotic recombination to separate neighboring variants in even the largest panels.

New experimental designs for the dissection of complex traits have been developed in yeast. Reciprocal hemizygosity scanning in diploid hybrids can identify causal genes (Weiss et al. 2018; Steinmetz et al. 2002) but can be prone to false positives (Wilkening et al. 2014). In addition, when performed in a pool, unequal representation of cells that are hemizygous for each gene can lead to missed causal genes due to differences in statistical power among genes. CRISPR-Cas9 genome engineering enables targeted introduction of genetic variants into genomes (DiCarlo et al. 2013). The high efficiency of homology-directed double-strand break repair in yeast has enabled massively parallel, pooled CRISPR-Cas9 strategies that can edit thousands of single variants in single cells. These methods make use of barcodes, guide RNA (gRNA) sequences, or repair templates as indicators of the intended edits in each cell (Chen et al. 2023; Sadhu et al. 2018; Roy et al. 2018; Sharon et al. 2018; Bao et al. 2018). However, not all genetic variants can be targeted by CRISPR, and not all regions in the genome have equal editing efficiencies. Further, because there is no sequencing of the targeted sites, there is no guarantee that any given cell in the pool is correctly edited. Lastly, genome-wide CRISPR engineering requires synthesis of large libraries of gRNAs and repair templates.

Beyond direct variant editing, CRISPR-Cas9 has been used to reshuffle yeast genomes by inducing mitotic recombination in diploid hybrids (Sadhu et al. 2016). However, this approach requires allele-specific gRNAs for the creation of DNA breaks in only one of the two genomes in the hybrid, which may not be possible in all genome regions. In spite of these technical problems, yeast studies have revealed considerable complexity in the genetic basis of quantitative traits even within individual mapped regions, several of which have been found to harbor multiple causal genes or variants (Steinmetz et al. 2002; Lutz et al. 2022; Schell et al. 2022; Gerke et al. 2009).

A thorough understanding of the genetic basis of complex traits in *S. cerevisiae* and other species remains a fundamental challenge that requires new approaches. Here, we introduce CRI-SPA-Map, a new strategy for the systematic dissection of complex traits. CRI-SPA-Map combines CRISPR-Swap genome engineering (Lutz et al. 2019) in diploid hybrid strains of *S. cerevisiae* with selective ploidy ablation (Reid et al. 2011). This engineering approach generates collections of haploid isolates with introduced variants that finely tile across the genome or a specific region of interest. The known location of the introduced variants makes genotyping of these isolates unnecessary for genetic mapping. Additionally, CRI-SPA-Map provides improvements in spatial resolution over traditional mapping approaches, enabling identification of causal genes that shape complex traits.

## Results

### The CRI-SPA-Map strategy for transferring alleles between yeast strains

We developed CRI-SPA-Map, a genome engineering strategy that efficiently replaces alleles of a recipient strain with those from a donor strain at defined positions and allows genetic mapping with or without sequencing (Figure 1). CRI-SPA-Map combines three key steps: 1) Mating of a haploid recipient strain carrying a selectable marker cassette (e.g., KanMX) at a known genome position and a haploid donor strain to form a diploid hybrid, followed by replacement of the cassette with donor alleles via CRISPR-Swap (Lutz et al. 2019) (“CRI”), 2) removing the donor strain chromosomes through selective ploidy ablation (”SPA”, (Reid et al. 2008, 2011)) to create haploid isolates, and 3) mapping of causal alleles by associating phenotype with the introduced donor alleles in a collection of isolates. A related method (“CRI-SPA”) also involves mating of a donor SPA strain to a collection of recipient strains, but is conceptually different in that it drives a reporter construct from the donor into a single genome location (Cachera et al. 2023). Instead, the final product of CRI-SPA-Map engineering is isolates in which selectable marker cassettes have been replaced with donor alleles.

**Figure 1.**
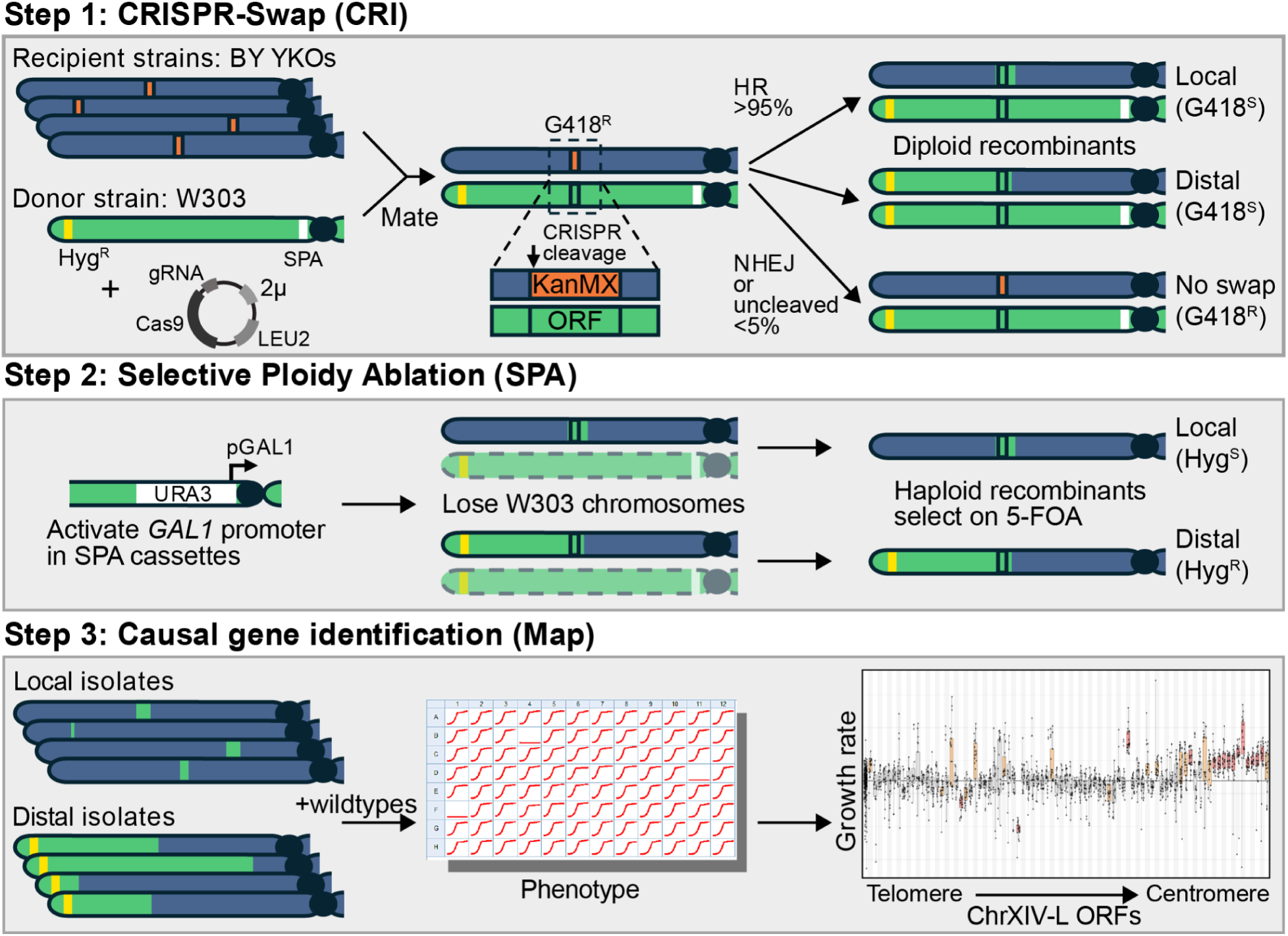
The CRI-SPA-Map procedure. See text for details. To implement and test CRI-SPA-Map, we used the BY4742 (BY) *S. cerevisiae* gene knockout (YKO) collection as our recipient strains and a W303 strain as our donor strain. Each of the YKO strains has the same BY MATα genetic background but a different open reading frame (ORF) replaced with a KanMX cassette that confers resistance to the chemical G418 (Winzeler et al. 1999; Giaever et al. 2002). The genomes of W303 and the reference strain S288C, which is nearly identical to BY, differ at approximately 9,476 single nucleotide variants (SNVs), including non-synonymous SNVs at ∼700 genes (Matheson et al. 2017). The modest genetic divergence of these strains narrows the search space for dissecting quantitative traits and provides an excellent test bed for CRI-SPA-Map.

We opted to focus on the left arm of chromosome XIV (chrXIV-L), which carries a known causal variant encoding a D30G amino acid substitution in *MKT1* that influences several complex traits including growth at high temperature, mitochondrial stability, and gene expression (Steinmetz et al. 2002; Dimitrov et al. 2009; Zhu et al. 2008; Deutschbauer and Davis 2005; Demogines et al. 2008; Sinha et al. 2006). We aligned BY and W303 sequences of chrXIV-L and identified 445 variants (including SNVs and small indels), in addition to Ty element polymorphisms at two positions. We selected 92 BY YKO strains where the KanMX cassette either replaced or neighbored these variants (STable1). As the W303 donor strain, we used a W303-SPA strain that is engineered to contain a cassette at each centromere that allows conditional loss of the W303 chromosomes (Reid et al. 2011). We further added a modified hygromycin resistance cassette (HphNT1Δlinker) near the chrXIV-L telomere of the W303-SPA strain and transformed this strain with the CRISPR-Swap plasmid (Lutz et al. 2019). This plasmid encodes a constitutively expressed Cas9 protein and the CRISPR-Swap gRNA that directs Cas9 to create a double-strand break within a linker sequence at the 5′-end of the KanMX cassette (SFigure 1).

We mated the BY MATα YKO strains to the W303-SPA MATa strain and selected for diploids harboring the CRISPR-Swap plasmid (Figure 1, Step 1). In the diploid, the CRISPR-Swap Cas9/gRNA can target the KanMX cassette located on the BY chromosome and create a double-strand break. Repair of this break using the W303 donor chromosome as a template removes the KanMX cassette and renders the cell sensitive to G418 (G418^S^). Rarely, cells are repaired by non-homologous end joining (NHEJ) or have no repair. Such cells are easily identified as they remain resistant to G418 (G418^R^). Importantly, it can be assumed that G418^S^ cells have repairs that include reinstatement of the deleted ORF with the W303 allele.

Replacement of the KanMX cassette with W303 alleles can occur by different mechanisms to create ‘local’ or ‘distal’ incorporation of W303 alleles (Anand et al. 2013). Local repairs are limited to the site of the ORF and its surroundings. Distal repairs extend from the centromeric side of the ORF to the telomere (Figure 1, Step 1).

We then induced SPA of the W303 chromosomes by activating the conditional *GAL1* promoter adjacent to each centromere by switching the cells to media containing galactose (Figure 1, Step 2). Transcription from the *GAL1* promoter leads to centromere instability and loss of the W303 chromosomes during mitotic division (Reid et al. 2008). We selected cells that lost all W303 chromosomes via transfer to media containing galactose and 5-FOA, a compound that is toxic to cells expressing the *URA3* gene that is present upstream of the *GAL1* promoter at each W303-SPA centromere.

In haploid cells after SPA, the local and distal repair types can be distinguished by selection on hygromycin (Figure 1, Step 2). Cells with distal repairs are resistant to hygromycin (Hyg^R^) because the HphNT1Δlinker cassette near the chrXIV-L telomere of the W303 strain is introduced as part of this repair process. In contrast, cells with local repairs do not introduce the cassette and are hygromycin sensitive (Hyg^S^).

At the final stage of selective ploidy ablation, we obtained an average of 171 colonies per targeted ORF (range of 42 – 305) growing on plates containing galactose and 5-FOA after three days (STable 2). The colonies for one YKO strain, YNL084C, took an additional two days to form on this medium (discussed below). On average, 67% of colonies were Hyg^S^ based on replica plating to YPD + Hyg, consistent with having cells that contain a local repair. The remaining 33% of colonies contained Hyg^R^ cells, consistent with distal repairs. Seven of the targeted ORFs had a notably small fraction (< 7%) of Hyg^R^ colonies, which we found was due to engineering an unexpected YKO strain (discussed below).

We chose at least eight Hyg^S^ and eight Hyg^R^ colonies from each YKO strain with sufficient colonies and used single-colony streaking to obtain CRI-SPA-Map isolates with a unique repair event from each colony. Each of the 1,501 isolates were assayed to verify the expected genotypic markers, including auxotrophic markers that distinguish BY from W303 (Lys^-^ and Met^+^), loss of the KanMX cassette (G418^S^), loss of the CRISPR-Swap plasmid (Leu^-^), and the absence or presence of the HphNT1Δlinker cassette (Hyg^S^ or Hyg^R^) for local or distal repairs, respectively (STable 2). Only 67 isolates did not have the desired phenotypes. These included all 16 isolates from the YNL268W strain, which we found to be Lys^+^, as stated in the YKO collection notes. All but 24 isolates were Leu^-^, indicating that the CRISPR-Swap plasmid is easily lost after selection for the plasmid is stopped. We cured 17 of these Leu^+^ isolates of the CRISPR-Swap plasmid to complete a set of 1,451 isolates for further analyses. In sum,

CRI-SPA-Map can be used to create large panels of recombinant haploids using standard, straightforward yeast laboratory techniques.

### CRI-SPA-Map transfers narrow tracts of DNA between yeast strains

To explore the genomic outcomes of CRI-SPA-Map engineering, we whole-genome sequenced 555 CRI-SPA-Map isolates (typically four local and two distal isolates, STable 2) created from the 92 YKO strains. We analyzed genotype calls at 8,257 high-quality variants across the genome (7,954 SNVs and 303 indels, STable 3). These analyses confirmed that CRI-SPA-Map successfully incorporated donor DNA from W303 into the recipient BY YKO strains on chrXIV-L (Figure 2A, SFigure 2).

**Figure 2.**
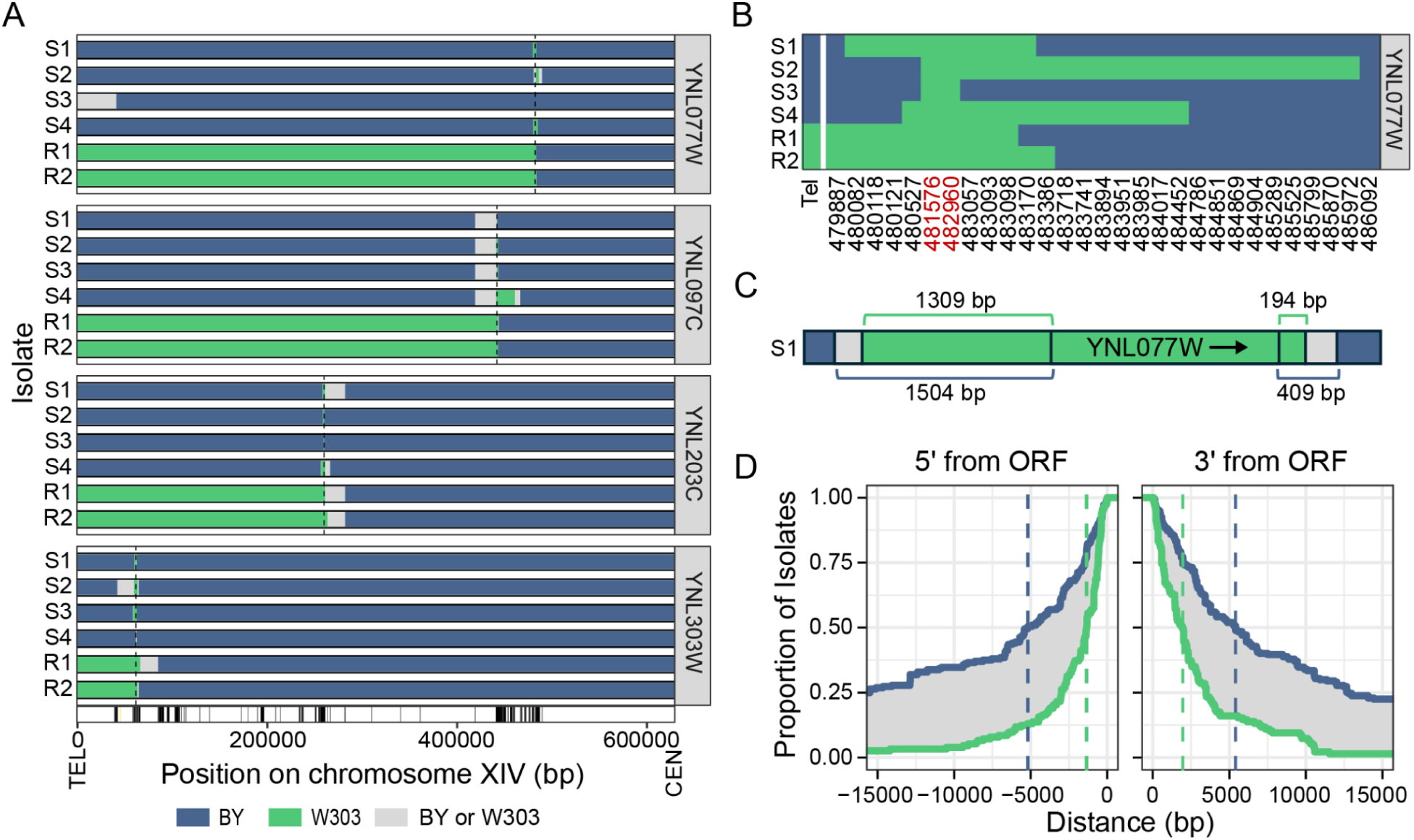
The CRI-SPA-Map isolates contain tracks of W303 variants on the left arm of chromosome XIV. (A) Position of the W303 tracts in four local (S for Hyg^S^) and two distal (R for Hyg^R^) isolates from four YKO strains. BY alleles are in blue and W303 alleles in green. The sequence between the last W303 variant in the tract and the first BY variant could originate from either the BY or W303 strain and is therefore designated as gray. The position of the targeted ORF is shown with a dotted line. The base pair (bp) position of the variants are marked by vertical lines with the position of the telomere (TEL) and centromere (CEN) at the bottom of the plot. (B) Allele designation at each variant site in isolates of YNL077W. Variant positions within the ORF are in red. (C) The position of the YNL077W ORF in the W303 variant tract in isolate S1. The distances to the first W303 and BY variants at each end of the ORF are shown. (D) Cumulative distribution showing the proportion of 286 isolates from 69 YKO strains with the first BY and last W303 variant at the indicated distance from the 5’-end and 3’-end of the ORF. The vertical dashed lines designate the maximum (blue) and minimum (green) of the median tract lengths. For distal isolates, only the distances on the centromeric side were analyzed. Isolates with mosaic tracts or W303 tracts that do not extend beyond the ORF were excluded.

In some isolates, we identified genome changes in addition to or other than the expected W303 variants on chrXIV-L (STable 2). In 3.4% (19/555) of the sequenced isolates, we observed W303 variants on the right arm of chromosome XII (chrXII-R) that extended from the rDNA locus to the telomere (SFigure 3). To screen for this recombination in the unsequenced isolates, we utilized a polymorphism at the *HAP1* gene located between the rDNA locus and the chrXII-R telomere that confers resistance to fluconazole (Saha et al. 2024). In total, we found that 2.8% (40/1451) of the isolates had a recombination event on chrXII-R (STable 4).

We found evidence of preexisting genome alterations and well-to-well contamination in our copy of the YKO collection in 12.3% (68/555) of the sequenced isolates. For example, all six sequenced isolates from the YKO strain YNL079C/*TPM1* had a duplication of chrIX, suggesting that this aneuploidy existed prior to CRI-SPA-Map engineering. This chromosome contains *TPM2*, the paralog of *TPM1*, and this aneuploidy may have been selected during the construction of the *TPM1* YKO strain (SFigure 4). All YKO strains noted previously to have a low fraction of Hyg^R^ colonies had genotypes that suggested inadvertent engineering of a YKO strain on another chromosome arm. For example, all local isolates of YNL173W had W303 variant tracts in and surrounding YMR316W on chrXIII-R, suggesting the YKO strain for this ORF had been targeted. If the majority of sequenced isolates from a YKO strain had genotypes suggesting an unintended YKO strain had been targeted or shared unexpected genome alterations, we flagged all isolates from this strain as having issues arising from the YKO collection.

Apart from the recombination at the rDNA locus, only 2.9% (16/555) of the sequenced isolates from 13 YKO strains had unexpected genotypes that might be due to CRI-SPA engineering. Specifically, heterozygous genotypes surrounding the targeted ORF were observed in six isolates, suggesting that these isolates are diploid. A recombination event on another chromosome was seen in six isolates. Aneuploidies were observed in five isolates, one of which also had heterozygous genotypes. Thus, CRI-SPA-Map-based engineering rarely caused unintended genome alterations. Because isolates with unexpected genotypes are rarely obtained and arise from unique YKO strains, their effects are easily buffered by obtaining multiple isolates from each YKO strain.

We used the W303/BY variants to gauge the length of W303 DNA introduced into each isolate (Figure 2A and SFile 1). There was variation in the length of introduced donor DNA even among isolates derived from one YKO strain (Figure 2B). In 18 isolates, mosaic tracts of W303 and BY variants flanked the targeted ORF (SFigure 5), similar to double-strand break repair outcomes previously observed in diploids (Gorter de Vries et al. 2019; Yin et al. 2017). To determine the length of the incorporated W303 tracts on each end of the ORF, we calculated the distance from the ORF to the last introduced W303 allele and to the first BY allele. The likely end of the repair tract is located between the corresponding variants (Figure 2C and STable 5). The median length of W303 allele tracts flanking each end of the ORF was between 1,532 bp for the last observed W303 allele outside of the ORF and 5,397 bp for the first observed BY allele. The spread between these two values is due to the sparsity of W303/BY variants (Figure 2D). In sum, our CRI-SPA-Map procedure efficiently replaces the KanMX cassette at each ORF position with tracts of donor DNA. Importantly, the introduced allele tracks are narrow, suggesting potential for high-resolution genetic mapping.

### Mitochondrial content varies among CRI-SPA-Map isolates

After mating two yeast strains, one mitochondrial type is thought to be maintained at random in the hybrid (Berger and Yaffe 2000). Therefore, we anticipated that the mitochondrial genotypes of the CRI-SPA-Map isolates would differ due to variants between the BY and W303 mitochondrial genomes. To control for potential effects of mitochondrial genotype in our genetic mapping, we created 16 wildtype control isolates by subjecting a BY4742 strain without a KanMX cassette to the CRI-SPA-Map procedure. The sequence of the mitochondria in the wildtype and CRI-SPA-Map isolates, even from the same YKO strain, showed a variety of patterns of BY and W303 variants (SFigure 6), indicative of both parental as well as recombinant mitochondria types (Fritsch et al. 2014). Thus, the wildtype isolates provide a well-matched control for CRI-SPA-Map genetic mapping.

### CRI-SPA-Map identifies a narrow causal genomic region

To test the utility of CRI-SPA-Map isolates for mapping complex traits, we used a plate reader to measure maximum growth rates in liquid YPD medium of 1,259 isolates (644 local (Hyg^S^) and 615 distal (Hyg^R^)) together with the 16 CRI-SPA-Map wildtype isolates (STable 6). Typically, seven local and seven distal isolates for each of the 92 YKO strains were phenotyped. To correct for the issues with our YKO collection, we removed isolates from YKO strains with preexisting aneuploidies, that were obtained from an unexpected ORF, or that appeared to arise from cross-contamination. We also removed isolates with recombination on chrXII-R identified in our *HAP1* screen. In total, we analyzed the growth rate of 1,057 isolates from 78 ORFs.

To identify ORFs with or near causal variants, we grouped the CRI-SPA-Map isolates by their targeted ORF and analyzed their growth rates compared to the wildtype isolates. This strategy, which we term “ORF-based mapping”, leverages the fact that any donor variants within the targeted ORF are guaranteed to be introduced in the CRI-SPA-Map engineering, and any additional introduced donor variants will extend from this ORF. As such, ORF-based mapping utilizes the known locations of the targeted ORFs and does not require knowledge of the precise genotype of each isolate.

The local isolates from 14 ORFs had growth rates significantly different from the wildtype isolates based on mixed linear models and a Bonferroni-corrected p-value threshold of p < 0.00064. A cluster of significant genes occurred at *MKT1* and its three neighboring genes, *SNN1*, *END3*, and *SAL1*. These four genes all had effects in the same direction, such that the introduced W303 allele caused an increase in growth rate (Figure 3, upper panel and STable 6).

**Figure 3.**
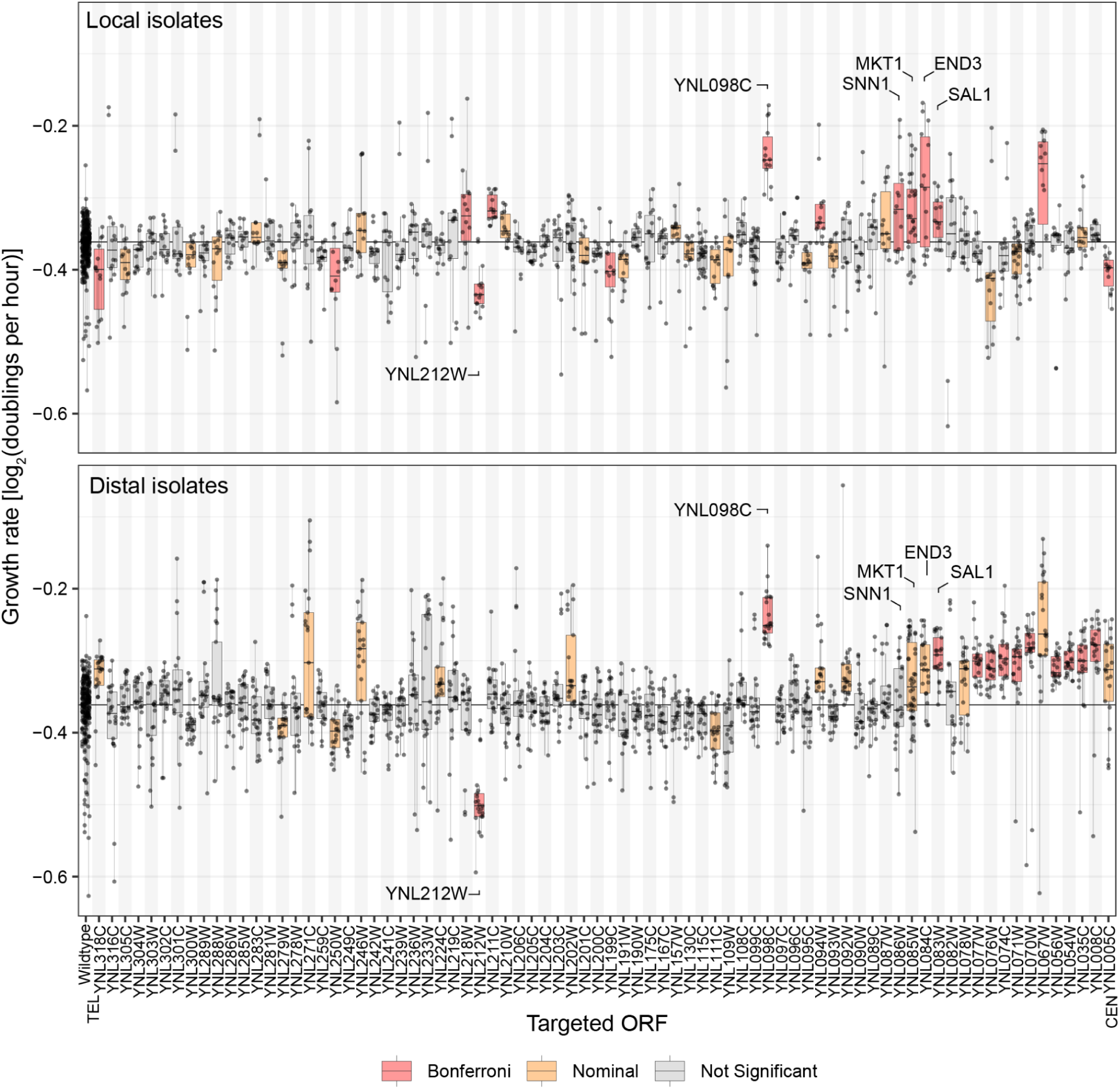
Growth rate in liquid YPD of local and distal isolates grouped by targeted ORF. Local isolates are in the top panel and distal isolates in the bottom panel. The targeted ORFs are ordered by position on the chromosome from telomere (TEL, left) to centromere (CEN, right). Growth rate is compared to the wildtype isolates shown on the far left. Red ORFs are significant at the Bonferroni threshold of p < 0.00064, and orange at p < 0.05. Phenotypes of 532 local and 525 distal isolates are shown.

Due to the compact nature of the yeast genome, where most ORFs are separated by a few hundred bases of intergenic DNA, the introduced W303 DNA tracts typically comprise at least some of the sequence of genes neighboring a given targeted ORF. Therefore, when several adjacent ORFs are targeted for CRI-SPA-Map engineering, we expect the effects of any causal variants to spill over into isolates of the neighboring ORFs due to linkage, creating a wave of significance. This pattern is seen at *SNN1*, *MKT1, END3*, and *SAL1*, but nowhere else along chrXIV-L. The ORFs YNL098C and YNL212W had significant effects, whereas their neighboring ORFs had non-significant effects or did not show the same direction of effect (Figure 3, upper panel).

To further investigate these observations, we analyzed distal isolates, which carry tracts of W303 alleles extending from the targeted ORF to the telomere. Distal isolates carry the W303 allele of a causal variant if the targeted ORF either contains the causal variant or lies between the causal variant and the centromere. In contrast, they should not carry the W303 allele if the targeted ORF lies on the far telomeric side of the causal variant. Some distal isolates from ORFs located in close proximity to the causal variant on the telomeric side can contain the causal allele, similar to local isolates.

Based on these expectations, growth rates of distal isolates plotted against the position of their targeted ORF, ordered from telomere to centromere, should rise or fall (depending on the direction of the causal allele effect) as the positions of the targeted ORFs become closer to the causal variant. This change in growth rate will then be followed by a plateau of similar growth rates for the ORFs that are on the centromeric side of the causal variant, since isolates from these ORFs should all contain the causal donor allele.

To test these expectations, we compared the growth rates of the distal isolates from each YKO strain to the growth of the wildtype isolates. To rule out the possibility that the introduction of the HphNT1 cassette into the distal isolates affected their growth, we compared the growth of the CRI-SPA-Map wildtype isolates to BY4742 strains with the HphNT1Δlinker cassette at the same location as the distal isolates and found that growth rates of these different wildtypes were not significantly different (p-value = 0.47; SFigure 7).

Based on the results of the local isolates, we anticipated that the distal isolates targeting ORFs in the vicinity of *MKT1* should show a rise in growth rates followed by a centromeric plateau. Indeed, the growth rate began to climb for ORFs just telomeric of *MKT1* (first without reaching statistical significance, e.g. at *SNN1*), became different from wildtype at nominal significance for *MKT1* and *END3*, reached Bonferroni-significance for *SAL1*, and continued to be nominally significant for 12 (9 of which with Bonferroni-significance) of the remaining 13 targeted ORFs between *SAL1* and the centromere. All of these significant ORFs had estimated allele effects in the same direction (Figure 3, lower panel).

In contrast, no rise or fall followed by a plateau was seen at YNL098C and YNL212W, which again stood out as prominent outliers as they did with the local isolates (Figure 3). Given the consistency in phenotypes among the local and distal isolates from these two genes and the unexpected specificity of their association signals compared to neighboring ORFs, a parsimonious explanation for the phenotypes of the YNL098C and YNL212W isolates is that they are not due to introduced W303 variants at these genes. Instead, these isolates likely contain variants that influence growth that were pre-existing in the YKO strains from which they originated (see below).

To examine the spatial resolution and statistical power of CRI-SPA-Map, we focused on the region of adjacent ORFs (*SNN1*, *MKT1, END3* and *SAL1*; YNL086W to YNL083W) that had reached statistical significance in the local CRI-SPA-Map isolates. Considering only the coding sequences of these four genes, this region spans 6,527 bp, which is about six times smaller than typical yeast growth QTLs (e.g. a median of ∼40 kb in (Nguyen Ba et al. 2022)). The average estimated fold increase in growth rate caused by the region demarcated by these four genes was 3% (STable 6). Thus, CRI-SPA-Map has the ability to map genomic regions shaping quantitative traits with high spatial resolution and statistical power sufficient to detect subtle effect sizes typical of natural genetic variation.

### Incorporation of sequence data improves mapping resolution

We incorporated our WGS data to determine if knowledge of the specific W303 alleles introduced into each isolate verifies and augments the ORF-based CRI-SPA-Map results. We tested for causality at each BY/W303 variant by comparing the phenotypes of CRI-SPA-Map isolates that carried the given W303 allele to those that carried the BY allele. These analyses grouped isolates based on their observed genotypes irrespective of which targeted ORF the isolates had been generated from. Because not all isolates had been sequenced, these analyses represent a trade-off between a smaller sample size and the additional precision gained from incorporating genome sequence information.

In the local isolates, these analyses showed a narrow peak, at which four variants located within 1,333 bp exceeded a permutation-based significance threshold of p < 0.000128 (Figure 4A and B; STable 7). The four variants were located inside or 135 bp upstream of the *SAL1* ORF. The most significant p-value was observed at the *sal1-1* allele, in which a 1-bp insertion in BY4742 creates a frameshift at codon 403 out of 545 (Chen 2004). W303 does not carry this frameshift. The distal isolates showed a wider peak extending from *MKT1* to *SAL1*, reflecting the more extensive linkage in distal than in local isolates (SFig 8). Thus, incorporation of genotyping data for CRI-SPA-Map isolates can further increase mapping resolution to a narrow region. This is especially true for local isolates, where variation in repair tract length on both sides of a targeted ORF provides a highly-resolved mapping population.

**Figure 4.**
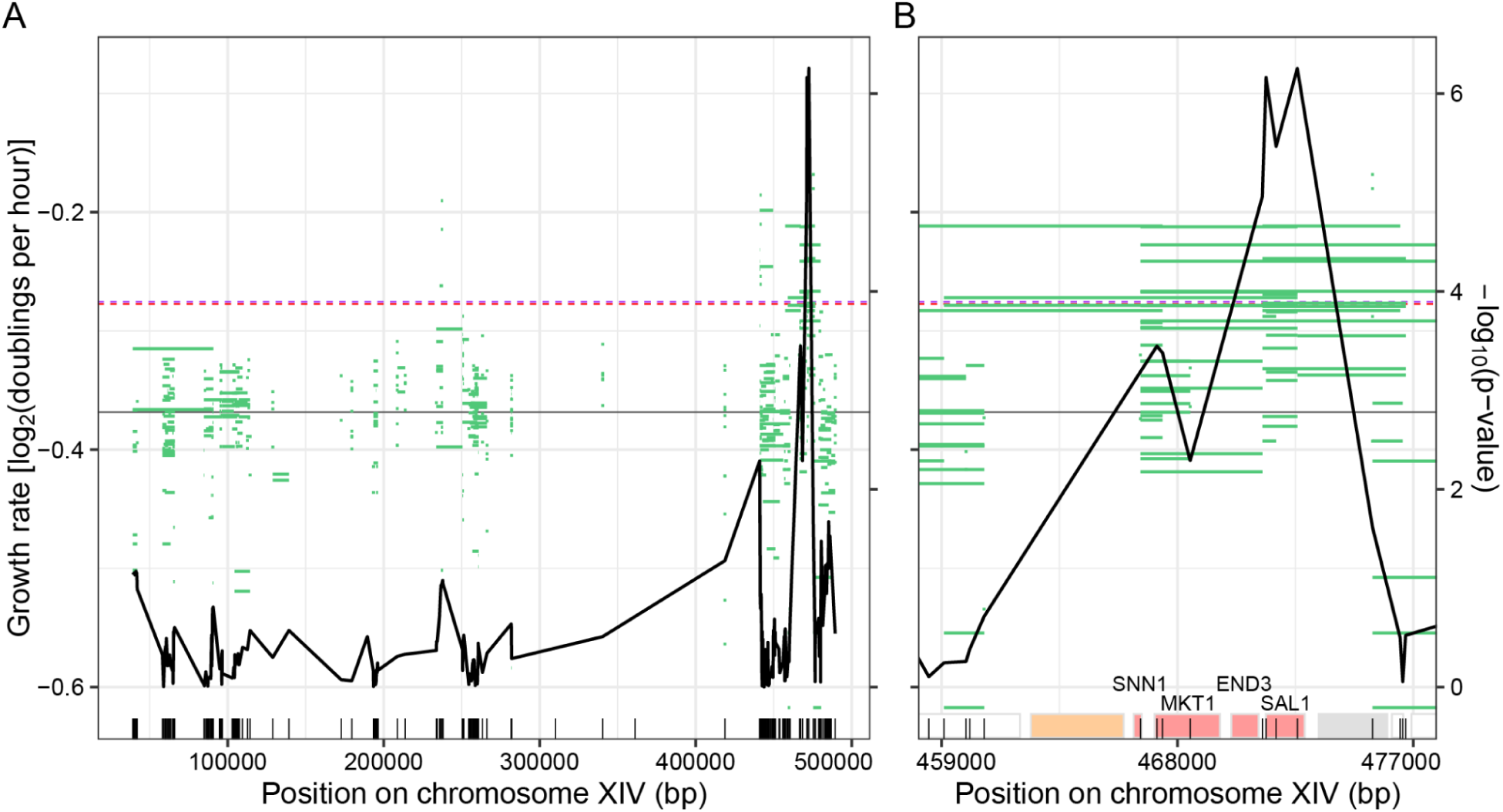
Incorporation of genotype data in CRI-SPA-Map for local isolates. (A) The variant positions on the left arm of chromosome XIV are depicted with vertical lines on the bottom of the plot. Green lines depict the W303 allele tracts in each of 332 local isolates as well as the growth rate of each isolate in liquid YPD. The median growth rate of all local isolates is denoted by a solid horizontal gray line. Dashed lines denote significance thresholds as determined by Bonferroni correction (red) and 1,000 permutations (purple). Note that these thresholds were nearly identical, resulting in overlapping lines. (B) Zoom-in on the region containing the four significant variants. The ORFs in this region are depicted as rectangles, with those used in the ORF-based mapping colored based on the significance of their effect as in the upper panel of Figure 2.

### Variants in *MKT1* and *SAL1* shape growth in liquid YPD

To independently validate and further dissect the region identified by CRI-SPA-Map, we used dual-cassette CRISPR-Swap (Lutz et al. 2019) to engineer the region spanning *MKT1*, *END3*, and *SAL1* in a BY strain (Figure 5A, Methods). We chose to focus on this region due to the significance of *SAL1* in both the local and distal ORF-based mapping and due to the fact that *MKT1* is known to affect numerous complex traits. The engineered region harbors a total of seven DNA differences between BY and W303 (Figure 5A). BY strains engineered to carry all seven W303 alleles had significantly higher growth rate compared to strains in which all seven BY alleles were reintroduced in this region (Figure 5B), validating the discovery of this causal locus by CRI-SPA-Map.

**Figure 5.**
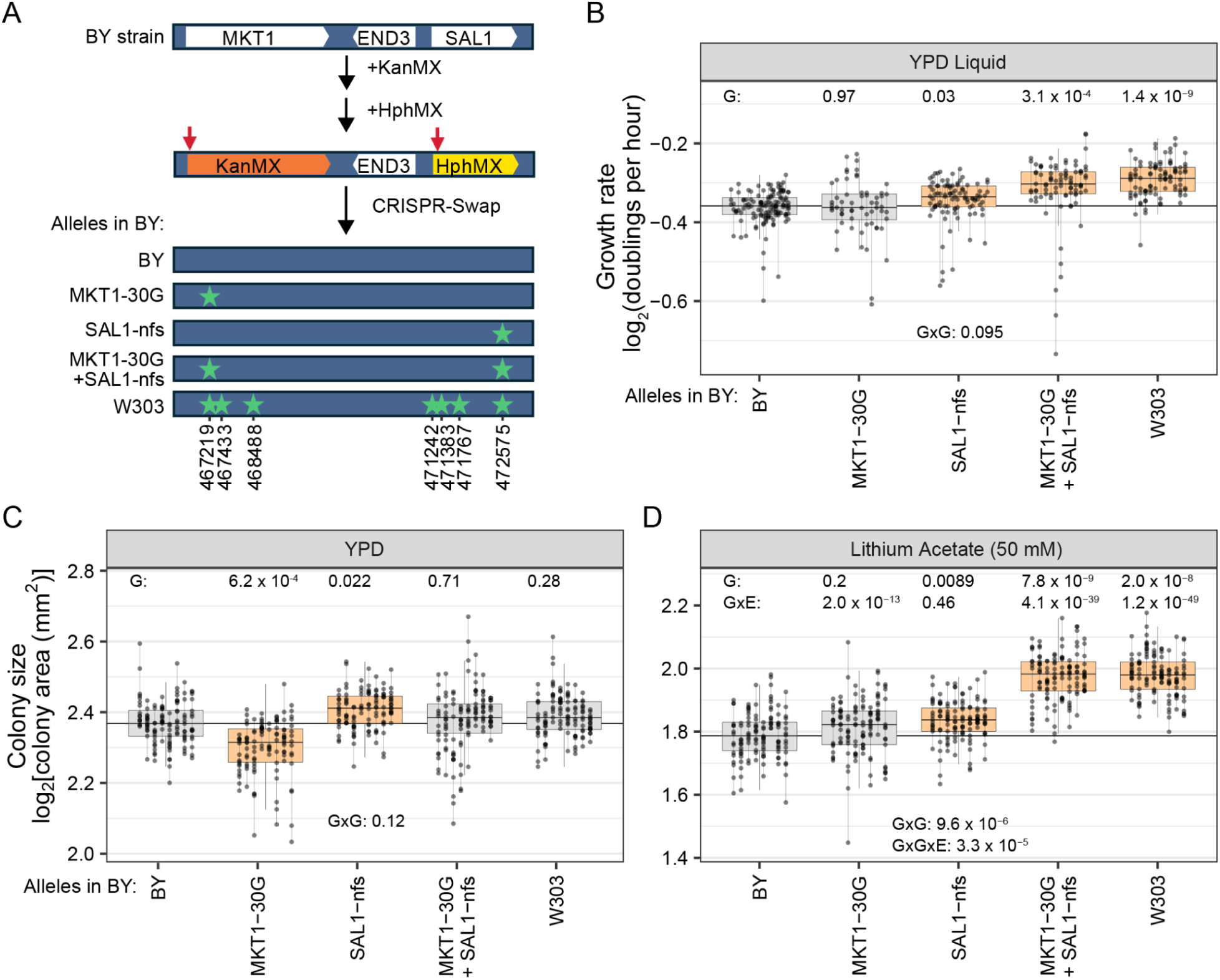
Effects of the *MKT1-30G* and *SAL1-nfs* alleles on growth. (A) Engineering of the *MKT1*-*END3*-*SAL1* locus by double-cut CRISPR-Swap. A BY strain was engineered to replace the *MKT1* ORF with KanMX followed by replacing the *SAL1* ORF with HphMX. These two cassettes were then cut by CRISPR and the locus was replaced by the depicted alleles (variant positions are starred) provided on a PCR amplified fragment. (B) Growth rate of the engineered strains in liquid YPD. P-values are shown for comparing each genotype to the engineered BY control (G) and for the test of the epistatic interaction between the *MKT1-30G* and *SAL1-nfs* (GxG). Box plots are shaded orange if p<0.05. Growth measurements from the same strain are connected by a vertical line. Box plots designate the central 50% of the data, with the median marked as a horizontal line within each box. The median of the BY wildtype strain is depicted as a horizontal line through the plot. (C) Colony size of engineered strains after 40 hours of growth on solid YPD. Boxplots and p-values for G and GxG are as in B. (D) Colony sizes on solid YPD with lithium acetate (50 mM). Boxplots and p-values are as in B. P-values for GxE interactions comparing the given condition to solid YPD are indicated for each genotype, along with that for the GxGxE term.

We hypothesized that the effect of this region is due to one or both of two variants implicated by prior work. In *MKT1*, the D30G variant is known to affect multiple traits, including sporulation efficiency, the expression of many genes, growth and formation of *petite* cells, and ethanol tolerance (Fay 2013). The *sal1-1* allele, which causes a frameshift in *SAL1* and had the highest significance in our variant-based mapping, is known to increase *petite* frequency (Dimitrov et al. 2009) and has been linked to slowed growth on ethanol (Schell et al. 2022). To test causality of these two variants, we introduced their W303 alleles (i.e., the glutamine-coding “G” allele at *MKT1-D30G* and the frameshift-free “*SAL1-nfs*” allele) into BY, individually and in combination. BY strains harboring *SAL1-nfs* grew significantly faster than the control (T-test p-value = 0.03), while the *MKT1-30G* allele had no effect on growth rate in liquid YPD (p = 0.97) (Figure 5B).

Combining the W303 alleles *MKT1-30G* and *SAL1-nfs* resulted in significantly faster growth than the BY control (Figure 5B), at a rate not different from that of BY strains carrying the entire W303 region (p = 0.15). Therefore, these two variants are responsible for most, and perhaps all, of the effect of this region. Strains harboring both alleles together grew significantly faster than those with the *MKT1-30G* allele alone (p = 0.018), as expected given the effect *SAL1-nfs* had in isolation. Interestingly, they also grew faster than strains carrying just the *SAL1-nfs* allele (p = 0.030), suggesting a contribution from *MKT1-30G* in the context of the *SAL1-nfs* allele even though it had no effect in isolation. While a formal test for a genetic interaction between *MKT1-30G* and *SAL1-nfs* failed to reach statistical significance (ANOVA, interaction p-value = 0.095), these results suggest epistasis between these two alleles such that their joint effect exceeds that of their individual effects.

### Gene-by-gene, gene-by-environment, and gene-by-gene-by environment interactions of the *MKT1* and *SAL1* causal alleles

To explore the extent of epistasis between *MKT1-30G* and *SAL1-nfs* (i.e., GxG) and the environmental dependence of the effects of these alleles (i.e., GxE), we grew our strains carrying all combinations of these two alleles on 11 diverse solid media and measured colony size after 40 hours of growth (SFigure 9, STable 8). Colony size is often used as a proxy for the number of cell divisions (a quantity closely related to the growth rate); however, smaller cell size can also lead to a smaller colony without a change in growth rate. The *SAL1-nfs* allele increased colony size on solid YPD (T-test p-value = 0.022), mirroring its effect on growth rate in liquid YPD. In contrast, the *MKT1-30G* allele significantly reduced colony size on solid YPD (p = 6.2 x 10^-4^), even though it had no effect on growth rates in liquid YPD (Figure 5B and 5C). The combined effect of the *MKT1-30G* and *SAL1-nfs* alleles on solid YPD was not significantly different from the BY control (p = 0.71) and was not different from the sum of the two individual allele effects (GxG test: p-value = 0.12). This absence of a joint effect of the two combined alleles differed from liquid YPD, where they had significantly increased growth rate. These analyses show that colony size on solid YPD and growth rates in liquid YPD are not interchangeable traits.

An epistatic relationship between the *MKT1-30G* and *SAL1-nfs* alleles was observed on lithium acetate (GxG test: p = 9.6 x 10^-6^) (Figure 5D). This interaction involved no individual effect for *MKT1-30G*, a slight increase in colony size for *SAL1-nfs*, and a joint effect of these alleles that led to significantly larger colony sizes than the sum of their individual effects (Figure 5D).

Compared to solid YPD, *MKT1-30G* and *SAL1-nfs* exhibited GxE in nine and five of the ten other conditions, respectively (GxE p-value < 0.05, SFigure 9, STable 8). For both individual alleles, this GxE involved some conditions in which the allele effect was stronger than on solid YPD, some in which it was weaker, and some in which it was not significant. There were no cases in which the allele effect was significant in the given condition and YPD but in opposite directions. Specifically, *MKT1-30G* reduced colony size compared to the wildtype control in seven conditions (p < 0.05), with no significant effect in the remaining conditions. Conversely, the *SAL1-nfs* allele increased colony size in six conditions and had no significant effect in the remaining conditions.

The epistatic effect of *MKT1-30G* and *SAL1-nfs* seen on lithium acetate was not present on solid YPD, resulting in an environment-specific genetic interaction (GxGxE p-value = 3.3 x 10^-5^; Figure 5D). Four additional conditions showed GxGxE (SFigure 9). These conditions differed from the solid YPD baseline in that the joint effects of *MKT1-30G* and *SAL1-nfs* were more similar to that of *MKT1-30G* than to that of *SAL1-nfs*, even though none of these conditions showed significant epistasis (SFigure 9). Collectively, these results exemplify the sensitivity of the effects of causal alleles on the environment and demonstrate that even narrow regions can harbor multiple causal DNA variants.

### Strain-specific variants in the YKO collection

Although the YKO strains were created in the same BY4742 genetic background, strain-specific variants in the YKO collections have been reported (Teng et al. 2013; Lehner et al. 2007; Van Leeuwen et al. 2016). We therefore asked whether background variants in the YKO strains could explain the phenotypic outliers in the CRI-SPA-Map isolates, particularly those of YNL098C and YNL212W. We searched our sequencing data for variants present in at least three of the sequenced isolates from a given YKO strain. We then removed variants that were present in the strains with a low percentage of Hyg^R^ colonies or the Lys^+^ strain. In total, we found 86 background variants derived from 53 of the 84 (63.1%) YKO strains. These background variants are predicted to cause 4 frameshifts, 6 premature stop codons, and 41 amino acid changes in the coding sequences of 45 different genes (STable 9). Given their prevalence and the difficulty of predicting their effects, we did not exclude isolates carrying background variants from our analyses.

One of the background variants we identified is a 1-bp insertion that causes a frameshift in the *GAL2* ORF in all isolates of YNL084C. *GAL2* encodes a galactose permease that is required for the utilization of galactose (Tschopp et al. 1986). We noticed during the CRI-SPA-Map procedure that the isolates of the YNL084C YKO strain took two days longer to develop colonies on the SGC 5-FOA plates, in which galactose is the carbon source (see above). It is likely that this frameshift in *GAL2* is the cause of the growth delay we observed.

We also identified high impact background variants in the YNL098C and YNL212W YKO strains. For YNL098C/*RAS2*, which encodes a GTP-binding protein (Kataoka et al. 1984), all six sequenced isolates had a variant causing a premature stop codon in *VPS13*, which encodes a lipid transport protein (Bankaitis et al. 1986). Five of the six sequenced isolates of the YKO strain YNL212W/*VID27,* which encodes a protein with a putative function in vacuolar import and degradation (Engel et al. 2025), had three background variants (one isolate had only 2 of these variants). Of these, one is a synonymous variant in *SLX4*, another causes a missense variant in *YLH47* (resulting in Q426R), and the final variant creates a frameshift (Y361X) in *VIP1*, which encodes an inositol hexakisphosphate kinase (Mulugu et al. 2007). It is unclear whether these background variants cause the large growth differences observed in these YKO strain isolates relative to the wildtype or to most other YKO strain isolates. Nonetheless, their presence suggests that such variants have the potential to influence the phenotypes of CRI-SPA-Map isolates created from the YKO collection.

## Discussion

We have introduced CRI-SPA-Map, a new strategy for genetic mapping of complex traits. CRI-SPA-Map generates collections of otherwise genetically identical individuals with variants introduced from a donor strain at defined genomic regions. We applied CRI-SPA-Map to dissect genetic variation on a chromosome arm between a W303-SPA strain and strains from the YKO deletion collection and uncovered a complex pattern of environmentally dependent epistasis at two causal variants.

CRI-SPA-Map has several key advantages for mapping genetic variation. First, using CRI-SPA-Map to target an ORF that had been deleted with a selectable marker cassette guarantees that this ORF is replaced with donor DNA in every isolate that lost the marker. This property makes it possible to conduct genetic mapping without any genotyping, as the known position of the deleted ORF specifies the genomic location at which donor DNA was introduced. While whole-genome sequencing of individual yeast strains is now routine and affordable, the materials and labor cost of constructing sequencing libraries for hundreds or thousands of strains remains a significant hurdle. Additionally, CRI-SPA-Map enables detection of causal variants in regions that might be hard to genotype with current sequencing methods, such as repetitive regions of the genome. Second, even genotype-free, ORF-based mapping has high resolution because the donor DNA tracts introduced by CRI-SPA-Map are short in local isolates and end in the vicinity of the targeted ORF in distal isolates. Resolution can be further increased if CRI-SPA-Map is applied to stretches of neighboring ORFs, where partially overlapping repair tracts of variable length create a panel of isolates with densely interlocking recombination points. Third, statistical power is equally high for all variants between the CRI-SPA-Map donor and recipient. Lastly, the cost of creating panels of CRI-SPA-Map isolates is minuscule, as only yeast media, plates, and equipment for handling and phenotyping isolates are needed.

We demonstrated the effectiveness of CRI-SPA-Map by engineering a collection of BY YKO strains with introduced W303 variants. To verify the fidelity of this engineering, we whole-genome sequenced more than 500 CRI-SPA-Map isolates. These data showed that tracts of W303 variants were integrated at the targeted KanMX cassette at each ORF in the YKO collection, and that CRI-SPA-Map introduced few off-target genome alterations.

In addition to validating CRI-SPA-Map, sequencing uncovered preexisting issues with our YKO collection. For example, one YKO strain (YNL079C) carried a likely duplication of chromosome IX, while others harboured unique background variants that likely affect phenotype, consistent with the known variability in the YKO collections (Teng et al. 2013). We also detected cross-contamination, with some wells containing an unexpected YKO strain or a mixture of YKO strains, which likely results from repeated use of this collection.

A series of safeguards can mitigate these issues with the YKO or similar collections when used in CRI-SPA-Map. First, engineering multiple ORFs surrounding specific variants or engineering densely and uniformly across the genome enables the flagging of ORFs with outlier phenotypes compared to their neighboring ORFs. In CRI-SPA-Map, the introduction of donor variants into the recipient strain is usually not limited to the targeted ORF, especially not in the distal isolates that carry all variants from the targeted ORF to the telomere. Therefore, the effects of a true causal variant are expected to be seen in the local isolates from neighboring ORFs and in all distal isolates from ORFs between the causal variant and the centromere. Second, a disproportionately small fraction of distal (Hyg^R^) compared to local (Hyg^S^) colonies in the CRI-SPA-Map procedure is a strong indicator that an ORF on a different chromosome arm was inadvertently engineered. We implemented these safeguards in our mapping, where we removed all isolates from YKO strains that produced a low percentage of Hyg^R^ colonies and prioritized for fine-mapping the region where multiple neighboring ORFs displayed significant phenotypes in the same direction. These safeguards are easily implemented without genotyping the CRI-SPA-Map isolates.

Using the CRI-SPA-Map local isolates for ORF-based mapping, we identified a 6.5-kb genomic region including four genes, *SNN1*, *MKT1*, *END3* and *SAL1*, as causal for the growth difference in liquid YPD. Mapping using the distal isolates suggested the effect of this region was due to variation in *SAL1*. Our sequencing data corroborated these results and further refined the resolution of this region. A frameshift in *SAL1* (*sal1-fs*, also called *sal1-1*) was a likely candidate for the growth difference, which we confirmed by engineering the *SAL1-nfs* allele into the BY background. Although the *MKT1-30G* variant affects numerous complex traits (a reason we opted to study chrXIV-L in this work), this variant did not have a significant effect on growth rates in liquid YPD. Identification of this region, and particularly the single gene *SAL1*, demonstrates the effectiveness of CRI-SPA-Map at high-resolution genetic mapping of genetically complex traits.

We explored the relationship between the *MKT1-30G* and *SAL1-nfs* alleles by growing BY strains carrying all combinations of these alleles in eleven different environments. These experiments revealed a gene-by-gene (GxG) interaction between the two alleles, gene-by-environment interactions (GxE) for each of the alleles, and gene-by-gene interactions between the alleles that are dependent on the environment (GxGxE). The interactions between the *MKT1-30G* and *SAL1-nfs* causal variants, and the dependence of their effects on the environment, highlight the complexity of variant effects even within a small genomic region.

The mechanism behind these condition-specific genetic effects on colony size and growth rate remains unclear but may involve the formation of *petites*, yeast cells lacking functional mitochondrial DNA. *Petite* cells are viable on glucose and other fermentable carbon sources but are smaller and grow more slowly than cells with intact mitochondria. Mkt1 and Sal1 have been shown to affect mitochondrial stability (Dimitrov et al. 2009), likely through different mechanisms: Mkt1, an RNA-binding protein, possibly targets mRNAs to the mitochondrion (Lee et al. 2009; Caballero et al. 2025), while Sal1, an ADP/ATP transporter, helps maintain mitochondrial ATP levels and inner membrane potential (Traba et al. 2008). Both the ancestral *MKT1-30G* allele in W303 and the derived *sal1-fs* allele in BY increase the rate of petite formation (Dimitrov et al. 2009). Introducing the *MKT1-30G* allele into BY led to decreased colony size in multiple conditions, possibly due to a higher proportion of *petite* cells within the colonies. Conversely, introduction of the W303 *SAL1-nfs* allele into BY increased colony size in multiple conditions, possibly by decreasing petite formation.

While we have shown that CRI-SPA-Map is effective and efficient at identifying genomic regions that harbor variation shaping complex traits, limitations do remain. Most prominently, CRI-SPA-Map relies on compatible pairs of donor and recipient strains. In our application here, we opted for the W303/BY4742 pair because a SPA strain and the YKO collection exist in these backgrounds. In addition, these strains are largely syntenous. In more distantly related strains, structural rearrangements such as translocations to other chromosomes may create incompatibilities following double-strand break repair.

A shortcoming of using the W303/BY pair for genetic mapping arises from the close genetic relatedness of these two strains, especially given that W303 was created in part by crossing BY to other strains (Matheson et al. 2017). This history has resulted in a distribution of DNA variants that is characterized by an unusual pattern, in which runs with many variants are separated by long stretches without any sequence differences. This limited our ability to precisely pinpoint DNA repair breakpoints. The low overall amount of variation between these two strains limits the number of causal variants that exist and may not be representative of more typical strain pairs.

Our current implementation of CRI-SPA-Map relies on SPA to remove the donor genome. To our knowledge, W303 is the only strain with SPA-cassettes integrated at all centromeres, enabling complete donor genome removal. Additional SPA strains could be generated using the established procedure (Reid et al. 2008, 2011), or alternatively, CRISPR-based strategies could be developed to destabilize donor centromeres (Zuo et al. 2017). Notably, donor genome removal is not strictly required for genetic mapping, as panels of diploid recombinants with loss of heterozygosity can also serve this purpose (Sadhu et al. 2016).

The CRI-SPA-Map procedure has the potential to be modified and expanded beyond its use here. CRI-SPA-Map is not limited to using the YKO collection as the recipient strain. Any arrayed collection in which selectable marker cassettes, with or without being coupled to the deletion of ORF sequences, could be used. Many of these arrayed collections exist, including in other species, which could allow the use of CRI-SPA-Map beyond *Saccharomyces cerevisiae*. CRI-SPA-Map creates recombinants by introducing variants from a donor strain into a recipient strain without meiotic recombination and sporulation. As a result, CRI-SPA-Map has the potential to create recombinants between infertile hybrid strains, perhaps allowing investigation of the differences in genetic architecture within as well as between species.

In conclusion, we have presented CRI-SPA-Map, a method for the efficient and effective introduction of donor DNA into a recipient genome. We have shown that panels of CRI-SPA-Map isolates can be used for high-resolution mapping of the effects of genetic variation on complex, quantitative phenotypes, even without genotyping. CRI-SPA-Map is a promising new tool for the dissection of the genetic basis of phenotypic variation.

## Material & Methods

### Computational analyses and code availability

Unless otherwise stated, all analyses were performed in R (R Core Team 2025). Analysis code is available at https://github.com/lawls18/CRI-SPA-Map.

### Yeast strains, oligos and media

Yeast strains and genotypes are listed in STable 10 and the YKO strains used are listed in STable 1. DNA oligo sequences used for plasmid and strain construction, genotyping and sequencing are listed in STable 11. Media formulations are in STable 12.

### CRI-SPA-Map strain engineering

As CRI-SPA-Map recipient strains, we used BY4742 YKO strains with ORF deletions on the left arm of chromosome XIV (*MAT*α *his3Δ1 leu2Δ0 lys2Δ0 ura3Δ0 ynlxxxxΔ::kanMX*) from a copy of the YKO deletion collection received as a kind gift from David Kirkpatrick. As the CRI-SPA-Map donor, we used a W303-SPA strain (YFA1743) with a fusion of mCherry::hphNT1Δlinker on the C-terminus of *EMW1* / YNL313C (45,308 bp from the telomere) and harboring the CRISPR-Swap plasmid (pFA0055, Addgene #131774). The full YFA1743 genotype is: *MAT***a** *can1-100 his3-11,15 leu2-3,112 trp1-1 ura3-1 CEN1–16::pGal1-URA3-K.lactis EMW1/YNL313C-mCherry::hphNT1Δlinker* + pFA0055-CEN-LEU2.

We removed the linker sequence within the mCherry-HphNT1 cassette from pFM689 (BFA0135, Addgene #74635 (Malcova et al. 2016)) by restriction digestion with BamHI and BglII to release the intervening sequence and then ligated the compatible DNA overhangs to form the plasmid mCherry-HphNT1Δlinker (BFA0262). We PCR-amplified the mCherry-HphNT1Δlinker cassette with primers OFA1369 and OFA1370 and then used this DNA fragment to transform the W303 SPA strain using a standard lithium acetate/PEG procedure (Gietz and Schiestl 2007). The location of the mCherry-Δlinker-HphNT1 insertion at the 3’-end of *EMW1* and the absence of an *EMW1* allele without the insertion was verified by PCR.

### Production of the CRI-SPA-Map isolates

The W303-SPA strain carrying the CRISPR-Swap plasmid (YFA1743) was mated to 92 BY4742 YKO strains (STable 1). After growing the haploids for one to two days on YPD plates, haploid strain pairs were mated by combining cells of each on a YPD plate. After eight hours of incubation at 30 °C, a volume of approximately 30 µl of the mated cells was resuspended in 100 µl of sterile water and spread onto an SRC -Lys -Trp -Leu plate and incubated at 30 °C for two days to select for cells carrying the CRISPR-Swap (pFA0055) plasmid and against the haploid strains. The raffinose instead of glucose in the media stops the strong repression of the *GAL1* promoter.

To start the selective ploidy ablation of the W303 chromosomes, diploid colonies (Lys^+^ and Trp^+^) carrying the pFA0055 plasmid (Leu^+^) were combined for a total of ∼30 µl volume of cells and resuspended in 1 ml of sterile water in a deep-96-well plate. The cells were then diluted 10,000-fold in sterile water and 100 µl were plated onto galactose-containing YPG plates to induce transcription from the *GAL1* promoter adjacent to each of the W303 centromeres. After incubation at 30 °C for three days, the YPG plates were replica-plated to SGC +5-FOA and grown at 30 °C for two days to ensure loss of *URA3* gene expression at each centromere.

To distinguish between local and distal repairs, each SGC +5-FOA plate was replica-plated to a YPD +Hyg plate and grown at 30 °C for one day. The percent of distal repairs was determined by dividing the number of colonies growing on YPD +Hyg by the number of colonies growing on SGC +5-FOA. The calculated fraction of colonies that are Hyg^R^ is likely an overestimate of the number of distal repairs because a colony growing on SGC +5-FOA may contain multiple CRI-SPA events leading to both Hyg^S^ and Hyg^R^ cells in the colony. Colonies growing on SGC +5-FOA but not YPD +Hyg were identified and saved as local repairs (Hyg^S^). Colonies growing on YPD +Hyg were saved from the YPD +Hyg plates as distal repairs (Hyg^R^).

To isolate single repair events, eight Hyg^S^ and eight Hyg^R^ colonies were streaked for single colonies on YPD plates. Three Hyg^S^ colonies only produced very small colonies in the single colony streaks on YPD, possibly due to genome instability These isolates, YNL079C_2, YNL082W_1, and YNL291C_1, were therefore not analyzed further. Each of the remaining isolates was tested for: 1) the auxotrophic markers of the BY4742 background by testing for growth on SDC -Lys, 2) loss of the KanMX cassette (YPD + G418), 3) the expected Hyg resistance (YPD +Hyg), 4) loss of the W303 chromosomes (SDC -Ura) and 5) loss of the CRISPR-Swap plasmid (SDC -Leu). After this screening, to ensure that we had at least 7 isolates from each strain, we obtained eight additional Hyg^S^ isolates from four YKO strains (YNL301C, YNL079C, YNL073C, YNL056W). In addition, we cured the CRISPR-Swap plasmid from some isolates by single colony streaking onto YPD and then replica-plating to SDC -Leu to identify colonies that had lost the plasmid. Thus, all isolates used in our further analyses were verified to be G418^S^, Lys^-^, Ura^-^, Leu^-^ and all local isolates were verified to be Hyg^S^ and all distal isolates to be Hyg^R^.

To create wildtype control isolates, BY4742 (Horizon Discovery, Cat #YSC1049) was mated to the W303 SPA strain carrying the CRISPR-Swap plasmid (YFA1743) and brought through the CRI-SPA procedure. Because no KanMX cassette was present in this BY4742 strain, these wildtype isolates should not be cut by the CRISPR-Swap machinery. Therefore, their nuclear genomes should be BY, but their mitochondrial genomes underwent the same shuffling as the CRI-SPA-Map isolates.

As additional control strains for distal CRI-SPA-Map isolates, we engineered three BY4742 strains to have the mCherry::hphNT1Δlinker cassette fused in-frame to the 3’-end of *EMW1* on ChrXIV-L as described above. These strains did not undergo the CRI-SPA-Map procedure.

### Whole-genome sequencing

We sequenced 555 isolates (STable 2) as well as several additional strain types: 1) six wildtype control isolates that carry the BY nuclear genome, 2) the three HphMX-tagged BY4742 strains used as controls for distal CRI-SPA-Map isolates, and 3) four W303 SPA strains containing HphMX tags at various genome locations. These additional strains were used for calling high-quality variants as described below.

The isolates and strains were arrayed in six 96-deep-well plates (STable 2) in 500 µl YPD and grown overnight at 30°C on an Eppendorf MixMate shaker set at 1100 rpm. After growth, each plate was centrifuged at 3700 rpm for 5 min to collect the cells. To remove the supernatant, plates were inverted over a collection container and while still inverted placed onto paper towels to allow excess liquid to be removed. The cells were resuspended in 800 µl of a solution containing 1 M sorbitol, 0.1 M EDTA, 14.3 mM β-mercaptoethanol, and 4 units of Zymolyase lytic enzyme (United States Biological) and sealed with a foil seal and incubated for 2 hrs at 37°C with occasional mixing to digest the cell walls. The cells were then centrifuged and the supernatant discarded as described above.

We isolated genomic DNA using the Omega, Quick DNA 96 Plus Kit following the Biological fluids and Cells protocol. After completing the DNA extraction protocol, we added 30 μL of DNA elution buffer (10 mM Tris-HCl, pH 8.5, 0.1 mM EDTA) prewarmed to 55°C to each well and incubated the plates at room temperature for 3 min prior to centrifugation. The concentration of the eluted DNA in each well was determined using the Qubit dsDNA BR assay kit (Thermo Fisher Scientific) in a 96 well format using a Synergy H1 plate reader (BioTek Instruments) using a self-made standard curve.

We prepared the genomic DNA for short-read sequencing on the Illumina Next-Seq platform using a modified Nextera (Illumina) based approach as previously described (Brion et al. 2020). Each 96-well plate of genomic DNA was processed alone or with one other plate. We diluted the genomic DNA to 5 ng in 11 µl in the wells of a new PCR plate and then added 4 µl of Tagment DNA buffer and 5 µl of Tagment DNA enzyme (diluted 1:20 in tagment DNA buffer). The tagmentation reaction plates were covered with PCR film and incubated at 55 °C for 10 minutes in a SimpliAmp thermal cycler (Thermo Fisher Scientific). Each sample of tagmented genomic DNA was then amplified with unique Illumina i5 and i7 index primer pairs to allow pooling of all samples prior to sequencing (STable 2). The amplification reactions contained 10 μL of the tagmentation reaction, 2 μL of a 5-µM index primer pair mixture, 5 μL of ExTaq buffer, 4 μL of dNTPs, 0.375 μL of ExTaq polymerase (Takara), and 29.625 μL of sterile molecular biology grade water. We then sealed the plates and ran the amplification reactions in a SimpliAmp thermal cycler (Thermo Fisher Scientific) using the following parameters: initial denaturation at 95 °C for 30 seconds, then 17 cycles of 95 °C for 10 seconds, 62 °C for 30 seconds, and 72 °C for 3 minutes.

We pooled 10 µl of each amplified sample from one plate and ran 200 µl of this mixture on a 2% agarose gel, and excised the DNA in the 450–550 bp size range. We then purified the DNA from the gel slice using the Monarch Gel Extraction Kit (NEB). We determined the concentration of DNA in the pooled libraries from each of the six plates using Qubit and then diluted each pool to 10 ng/µl. We then combined 5 µl from each of the six pools to create the final library pool.

The average fragment size of the final pooled library was 585 bp, as determined using an Agilent TapeStation. High-output sequencing of 150-bp paired-end reads was performed on an Illumina NovaSeq 6000 instrument SP run at the University of Minnesota Genomics Core. The average quality scores were above 30 and an average of 625,816 read-pairs were obtained for each sample. The median read depth across the nuclear genomes was 7.8, with a range of 1.4 to 27 across all samples (STable 2).

### Identifying variants between the BY4742 and W303 SPA strains

Analyses in this section used GNU parallel (Tange 2022). We used available genome sequences of BY4742 (http://sgd-archive.yeastgenome.org / sequence / strains / BY4742 / BY4742_Toronto_2012) and W303 (Matheson et al. 2017) to determine the nucleotide variants on chrXIV-L. We aligned the chrXIV-L sequences from BY4742 and W303 to the S288C reference genome (sacCer3 R64-5-1) and used show-snps to create two variant lists using MUMmer (v 3.23) (Kurtz et al. 2004). These lists were then transformed into VCFs using a custom python script. The VCFs were then combined to identify the BY4742/W303 chrXIV-L variants. We identified 445 variants and two breaks in synteny due to Ty polymorphisms (STable 1). We used these variant positions to select the YKO strains with the nearest KanMX cassette for CRI-SPA-Map.

To determine the genotype of the CRI-SPA-Map isolates and the other sequenced strains we used BWA (v 0.7.17) (Li 2013) to align the sequencing reads to the S288C reference genome. We then used samtools (v 1.16.1) (Danecek et al. 2021) to remove reads tagged as duplicates, unaligned, mapping to more than one location, or with a mapping quality of less than 30. Using these alignments, haploid genotypes of all samples were called across the genome using bcftools (v 1.16) (Danecek et al. 2021) to create a genome-wide VCF across all sequenced CRI-SPA-Map isolates and the other sequenced strains. We retained variants with a quality score of at least 20.

To identify any variants present between BY4742 (the background of the strains in the deletion collection) and the S288C reference genome, we used a custom R (v 4.4.3) script to extract the columns of the VCF of the six wildtype isolates and the three HphMX-tagged BY4742 strains. After filtering for variant sites at which 75% or more of these nine BY4742 isolates and strains carried the same alternative allele, we identified 86 sequence differences between BY4742 and S288C. We then used the same criteria on the four sequenced W303 SPA strains and found 8,305 sequence differences between them and the S288C reference. Comparison of these two variant lists identified 67 variants that were shared between BY4742 and W303 SPA but different from the S288C reference. This left 8,257 variants across the nuclear genome to distinguish BY4742 from W303 SPA with 377 variants on chrXIV-L (STable 3).

To determine mitochondrial variants between BY4742 and S288C, we analyzed the mitochondrial sequences of the three HphMX-tagged BY4742 strains, which all carry the same BY4742 mitochondrial genome. We did not analyze the six wildtype isolates due to the mitochondrial shuffling that occurs during the mating step of CRI-SPA-Map. After filtering for variant sites at which all three BY4742 strains had the same alternative allele, there were five mitochondrial variants between BY4742 and S288C. We then compared the four sequenced W303 SPA strains to S288C and found 53 mitochondrial variants called in at least 75% of the W303 SPA strains. All five mitochondrial variants identified between BY4742 and S288C were also present between W303 SPA and S288C, leaving us with 48 variants to distinguish the BY4742 and W303 SPA mitochondrial genomes (STable 3).

### Identifying variants in the YKO strains

Using the genome-wide VCF described above, we retained variants that were called and had a genotype quality score of at least 30 in a minimum of 90% of the 555 sequenced CRI-SPA-Map isolates. To investigate whether any of these variants likely arose before CRI-SPA-Map and were therefore present in the original YKO strain, we looked for any remaining variants that had at least three isolates from the same targeted ORF with alternate allele calls. After identifying these variants, we manually removed the variants that arose from YKO strains with a low percentage of Hyg^R^ colonies or from the Lys^+^ YNL268W YKO strain.

This criteria identified 69 variants that are different between S288C and BY, 9 fingerprint variants (including 1 located on ChrXIV, which we identified during our other analyses), and 86 strain-specific background variants (S Table 9). We used Ensembl VEP (McLaren et al. 2016) release 115 to predict the consequences of the variants.

### Genotype screening of CRI-SPA-Map isolates

We designed criteria (described in detail below) to detect unexpected outcomes of the CRI-SPA-Map procedure as well as potential strain contaminations or mislabelling. These criteria included sequencing data and other information as detailed below. These criteria were not mutually exclusive, and some isolates were flagged by multiple criteria. The results for each criterion and isolate are provided in STable 2. Isolates were removed from analyses depending on the nature of the specific criterion and analyses. For example, an isolate with W303 variants on ChrXIV-L that do not include the targeted ORF (suggesting that it was engineered at a gene other than the one intended but on the same chromosome arm) would be excluded from ORF-based mapping but would be used when incorporating individual isolate genotypes. A list of the isolates used in each analysis is provided in STable 2.

#### Strains with low percentage of HygR isolates

After replica-plating colonies to YPD + Hyg, an average of 33% of colonies were Hyg^R^ (STable 2). Seven YKO strains had notably lower percentages (0 – 7%). A likely reason for this low recovery of Hyg^R^ isolates is that these YKO strains carried deletions of genes different from those we targeted. Specifically, if the KanMX cassette was on a different chromosome arm than ChrXIV-L, the HphMX cassette on ChrXIV-L would not be introduced into the given isolate during repair of the double-strand break. In this scenario, Hyg^R^ colonies would be rare, as they can only be obtained from infrequent spontaneous breaks and repairs on ChrXIV-L. All sequenced isolates from the seven YKO strains with few Hyg^R^ colonies were also flagged by sequencing-based analyses (see below), further supporting that these isolates arose from YKO strains with gene deletions on other chromosomes. All 112 isolates arising from the seven YKO strains with low Hyg^R^ recovery were flagged according to this criterion.

#### Aneuploidy

To determine if the CRI-SPA-Map isolates exhibited large-scale copy number variants (CNVs), we used CNVnator (v 0.4.1) (Abyzov et al. 2011) to divide the chromosomes of the S288C reference genome into non-overlapping 5,000 bp bins and to identify bins with read depth differing from the average read depth across the nuclear genome (STable 2). Isolates were considered aneuploid when they contained regions of higher or lower read depth (“eval-1” value reported by CNVnator < 0.00000539, corresponding to 0.05 / the total number of CNVs reported by the software) that extended for at least half of the given chromosome. We identified 11 sequenced isolates with aneuploidy (SFigure 4).

Of the six sequenced CRI-SPA-Map isolates from the YNL079C (*TPM1*) YKO strain, five exhibited a duplication on chromosome IX. Upon closer examination, the 6th isolate also showed a duplicated segment just below half of the length of chromosome IX. Thus, all sequenced isolates from the YNL079C YKO strain showed evidence for duplication of this chromosome, which contains *TPM2*, the paralog of the deleted *TPM1* gene. Therefore, we flagged all 24 isolates derived from YNL079C. An additional five isolates from three additional YKO strains showed evidence for aneuploidy. In total, we removed 29 isolates from four YKO strains.

#### Mosaic tracts and tracts with heterozygous variant calls

We flagged 26 isolates from 22 ORFs because they showed more than one W303 variant track on ChrXIV-L. Analysis of these variant tracts revealed two categories: mosaic tracts and heterozygous variant call tracts.

Mosaic tracts were observed in 18 isolates (SFigure 5). We defined these mosaic tracts as tracts of homozygous W303 variants interrupted by tracts of at least three homozygous BY variants. The isolate YNL099C_S2 has a tract of W303 variants adjacent to but not spanning the YNL099C ORF and would normally be flagged for not having W303 variants spanning the ORF. However, there is one BY variant between this W303 tract and the ORF and there are no additional variants for 22,302 bp. All other isolates of this YKO strain have tracts of W303 variants spanning the ORF and all isolates had matching DNA fingerprints (see below). We conclude that this isolate has a mosaic tract arising from a correctly targeted ORF.

Heterozygous variant calls were observed within the repair tracts of 8 isolates. These isolates were homozygous for W303 alleles in the targeted ORF and had runs of at least six variants with heterozygous calls showing both BY and W303 alleles. The rest of their genomes were homozygous for BY alleles. We mated 4 of these isolates to both MATa and MAT⍺ strains and they mated to only MATa. All of the isolates flagged by this analysis had distal repairs, likely due to the difficulty of confidently calling a short heterozygous tract that may have arisen from local repair. We noticed that the isolates with heterozygous tracts grew faster than isolates from the same YKO strain without apparent heterozygous tracts. We hypothesize that these isolates are MATa diploids that underwent separate DNA repairs on two BY chromosomes present in a tetraploid or during the G2 phase of the cell cycle, followed by an error in chromosome segregation. After SPA, the cell then became diploid.

#### BY variants in the ORF

Because the YKO strains carry KanMX cassettes instead of the sequence of the deleted gene, CRI-SPA-Map isolates derived from YKO strains should have W303 alleles at the targeted gene. Therefore, isolates with a BY allele call at a variant within the targeted ORF were flagged in this analysis, as such calls suggest that the targeted ORF had not been deleted in the BY YKO strain used in the CRI-SPA-Map cross. Instead, the KanMX cassette had likely been present (and was then replaced by CRI-SPA-Map), at a different gene than the targeted one. For YKO strains with ≥ 75% of sequenced CRI-SPA-Map isolates with BY variants in the targeted ORF, we flagged all Hyg^S^ and Hyg^R^ isolates, sequenced or unsequenced, as we assumed that the majority of these isolates were not from the intended deletion strain. All 72 isolates from four YKO strains were flagged based on this criterion, of a total 80 isolates from 10 YKO strains.

#### W303 variant tracts not spanning the targeted ORF

For the targeted ORF YNL133C, none of the observed tracts of introduced W303 variants in local isolates spanned this ORF. These isolates are likely from the YNL092W YKO strain, as their DNA fingerprints (see below) and repair tract positions matched isolates from YNL092W. The YNL133C and YNL092W strains are in adjacent wells in the YKO collection, suggesting possible cross-contamination. We flagged all 16 isolates (Hyg^S^, Hyg^R^, sequenced, and unsequenced) from YNL133C.

#### Deletion strain fingerprints

We identified nine sequence variants that segregated among the CRI-SPA-Map isolates (STable 9). For eight of these variants, the alternative allele was observed in half (range of 46 – 53%) of our isolates, and the remaining variant was observed in 16% of our isolates.

One method by which strains were created in the YKO collection was to replace the ORF with the KanMX cassette in a BY4741 / BY4742 diploid followed by sporulation. This method was used to create 70% of the haploid YKO strains (Giaever et al. 2002). Thus, any DNA variants between BY4741 and BY4742 would be expected to segregate in the YKO collection. To test this, we performed blastn (BLAST+ tools (Camacho et al. 2009)) searches using each of the nine variants and flanking sequences to query databases created from partial genome sequences of BY4741 and BY4742 (Song et al. 2015). We found that the BY4741 and BY4742 differ at the eight variant positions with the highest alternative allele frequency (positions in S288C are chrI:41475 G/T, chrIV:769328 G/A, chrVIII:499958 C/T, chrVIII:511255 C/T, chrXIV:189585 C/T, chrXV:773264 G/C, chrXVI:667520 T/G, and chrXVI:677102 A/G). The origin of the lower frequency variant (chrXI:589175 T/A) is unknown.

We used these nine variants as a DNA fingerprint to verify that isolates obtained from the same deletion strain have the same variant call at each fingerprint position. A consensus DNA fingerprint was determined for each deletion strain when at least five of the six sequenced isolates had the same variant calls. If a consensus DNA fingerprint could not be determined, all isolates from this YKO strain were flagged. If only one isolate did not match the consensus fingerprint at more than one of the nine positions then this isolate was flagged. This analysis flagged 131 isolates from 11 ORFs.

#### Recombination at locations other than ChrXIV-L

We searched for mitotic recombination events on chromosome arms other than ChrXIV-L by locating tracts of W303 variants elsewhere in the genome. This analysis suggested mitotic recombination at locations other than ChrXIV-L in 42 CRI-SPA-Map isolates from 28 YKO strains. Three isolates had W303 variant tracts on two chromosome arms other than ChrXIV-L (SFigure 3).

Nearly half of these 42 isolates had W303 variant tracts on the right arm of chromosome XII (chrXIIR; 9 Hyg^S^ and 10 Hyg^R^ isolates). In these 19 isolates, the tracts of W303 variants extended from near the rDNA locus to the telomere of chromosome XIIR, suggesting increased mitotic recombination at this highly repetitive locus.

Due to the length and frequency of these recombination events on chrXII, we wanted to exclude all isolates that carry these events, including those that were not sequenced. This was possible due to the existence of a polymorphism in the *HAP1* gene located between the rDNA array and the chrXIIR telomere. BY strains have a Ty1 insertion at the 3’-end of the *HAP1* ORF (*hap1-Ty1*), while the W303 *HAP1* gene is wildtype (Gaisne et al. 1999). BY strains with the *hap1-Ty1* allele are more susceptible to fluconazole than strains with the *HAP1* allele (Saha et al. 2024).

To screen for mitotic recombination of chrXII-R, all isolates (sequenced and non-sequenced) were pinned from glycerol stocks to YPD PlusPlates using a ROTOR+ instrument (Singer Instruments, Roadwater, United Kingdom). After three days of growth, the colonies were pinned with the ROTOR to Plus plates (Singer Instruments) containing SDC + 16 µg/ml fluconazole (Cayman Chemical) and incubated at 30°C for two days. Colony size was then determined using a PIXL® instrument (Singer Instruments). Local isolates were considered likely to carry the wildtype, W303 *HAP1*, allele if their colony size was at least 30% larger than the median colony size of all local isolates from the corresponding YKO strain. For distals, the colony needed to be at least 20% larger to be considered. The genotypes of such isolates were then verified by PCR using primers specific to each *HAP1* allele (primers OFA1691 and OFA1692 amplified wildtype *HAP1* and OFA1691 and OFA1693 amplified *hap1-Ty1*. This screen identified 24 additional *HAP1* strains (5 Hyg^S^ and 19 Hyg^R^).

Across all 1,451 screened isolates, we identified 40 (2.8%) that had mitotic recombination on chrXII that likely extends from the rDNA locus to the telomere (STable 4). We did not detect mitotic recombination in 16 wildtype isolates that went through CRI-SPA-Map without a Cas9/gRNA cleavage. It is possible that CRI-SPA-Map increases the frequency of mitotic recombination.

#### Other filtering

Two isolates (YNL291C_S2 and YNL301C_R2) were flagged for having unexpected tracts of W303 variants based on their hygromycin susceptibility. YNL291C_S2 had a distal tract and YNL301C_R2 had a local tract. This suggests a possible strain mix-up after testing growth on YPD + Hyg and before sequencing.

#### Filtering summary

Application of these analyses to the 1,451 isolates without auxotrophy issues resulted in flagging of 273 isolates across 65 YKO strains. Of these, 139 isolates (51%) from 37 YKO strains were flagged by analyses that did not require sequencing. A large majority of the flagged isolates (209 from 18 YKO strains, 77%) were flagged due to likely problems with our copy of the YKO collection. There were 1,178 (81%) isolates spanning 78 YKO ORFs without any flags. The isolates used in each analysis are listed in STable 2.

### Locating tracts of W303 variants in the CRI-SPA-Map isolates

DNA variant calls were simplified based on the called genotype and genotype quality score as follows: a site with a quality score of zero was designated as “no call”, a call with genotype quality score higher than zero but below 30 was designated as “low quality,” and a call with a score of at least 30 was designated as either “BY” or “W303” depending on the called allele. An additional variant was added marking the centromere of ChrXIV and each isolate was assigned “BY” at this site. Using the resulting allele designations at the 377 variants on ChrXIV-L plus the ChrXIV centromere call, we identified runs of at least three consecutive variant calls with the same parental (BY or W303) allele. We discarded “no call” and “low quality” calls and subsequently merged any consecutive runs of the same parental allele. For each isolate, we determined the genomic coordinates of the start and end of each run using the variant locations marking the start and end of the identified tracts. If a run began at either of the two most telomeric variants on ChrXIV-L, we considered the run to start at the end of the chromosome.

To estimate the length of the W303 DNA segment introduced into the BY background, we used variant runs on ChrXIV-L to determine the distance between the targeted ORF and the 5’ and 3’ ends of the repair tracts. For example, moving 5’ from the start codon of the ORF we identified the farthest W303 variant call in a run of W303 variants. The distance between the start of the ORF and this farthest W303 variant position is the minimal repair length on the 5’ end of the ORF. The maximal repair tract length is the distance from the start of the ORF to the first BY variant in a run of BY variants. We calculated at least one repair tract length in 286 isolates. If the run of W303 variants did not extend outside of the ORF, the isolate was not considered. This results in a conservative overestimate of the median minimal amount of introduced W303 DNA, because very short W303 DNA tracts that replace the ORF but no flanking variants are not considered. Additionally, we removed all isolates derived from the 11 YKO strains in which the deleted ORF did not fully lie between our most telomeric and centromeric BY/W303 variants, as the lack of variants near these 11 ORFs would likely cause an overestimate of repair tract lengths. Finally, we excluded any isolates flagged for unexpected auxotrophies or issues suggestive of deletion collection problems (low Hyg^R^, BY ORF variants, W303 variants not spanning ORF, mosaic tract, heterozygous reads, and inconsistent deletion strain fingerprints).

### Measuring growth rates in liquid YPD

Growth rates in liquid YPD were assayed using a Synergy H1 (BioTek Instruments) plate reader. Each of the Hyg^S^ isolates were arrayed once on one of nine 96-well plates with the exception of isolates of YNL055C, YNL070W, YNL071W, YNL099C, YNL115C, which were present twice on a single plate (plate 9), and isolates of YNL085C, which were present on two plates (6 and 9).

Each of the nine plates contained the same 16 CRI-SPA-Map wildtype strains and 3 blank wells arrayed in different positions. The Hyg^R^ isolates were arrayed once on one of eight 96-well plates. On each Hyg^R^ plate were the same 6 CRI-SPA-Map wildtype strains and the same 4 BY4742 *EMW1*-*mCherry::hphNT1Δlinker* wildtype strains and 2–3 blank wells arrayed in different positions. The nine Hyg^S^ plates were phenotyped twice and the eight Hyg^R^ plates were phenotyped three times.

Prior to phenotyping, all isolates were pinned from glycerol stocks to YPD or YPD + Hyg agar for the local and distal isolates, respectively. After 3 days of growth at 30 °C the plates were stored at 4 °C for up to two weeks.

We inoculated each culture for the plate reader runs with actively growing starter cultures. For the first run of a plate, starter cultures of 400 μl of YPD in a 2-ml deep-96-well-plate were inoculated with cells from the agar plates stored at 4 °C. These starter cultures were then sealed with foil and stored at 4 °C until they were used to inoculate new starter cultures for the second and third plate reader runs. To grow the starter culture plate, we sealed it with a Breathe Easy membrane (Diversified Biotech) and placed it on an Eppendorf MixMate set at 1100 rpm. After overnight growth at 30 °C, we diluted the starter cultures in 100 µl of YPD to an approximate OD_600_ of 0.05 in a shallow 96-well plate. We then sealed the plate with a Breathe Easy membrane and placed it in the plate reader.

The strains were grown for ∼18 h in the plate reader set at 29 °C with OD 600 absorbance readings taken every 6 min for 181 cycles with orbital shaking (1 mm) between reads.

Growth rates were determined using the Growthcurver R package (Sprouffske and Wagner 2016) to fit growth curves to each well and identify the inflection point at which the yeast culture begins to exit exponential growth. The growth rates were then defined as the number of doublings per hour at the inflection point. Statistical analyses used the log_2_ of the growth rate (“growth rate” hereafter).

To avoid possible confounding by systematic differences between plates, we corrected growth rate for the specific plate they were grown in using the following mixed linear model:

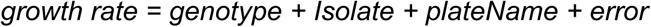

Growth rate was used as the dependent variable, modeled as the sum of a series of random effects including the plate the measurement stemmed from. To fully represent the data during pre-processing, the model also included random effect terms for the genotype (i.e. the targeted ORF) and the isolate from which a given measurement was taken. We obtained corrected phenotypes by subtracting the effects of *plateName* from the raw measurements and used these residuals in further analyses.

### ORF-based genetic mapping

Isolates from each targeted ORF were compared, one ORF at a time, to the set of 16 wildtype isolates. Multiple measurements were available from each isolate (see above). We analyzed growth rates using the following mixed linear model:

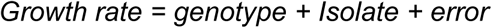

Here, *genotype* is a fixed effect denoting whether a given measurement is from a wildtype or from an isolate from the targeted ORF. The random *Isolate* term denotes the isolate a given measurement was taken from, and controls for any systematic differences among isolates. Statistical significance of *genotype* was determined using ANOVA. Multiple testing correction was performed using the Bonferroni procedure, correcting for the number of ORFs in the given analysis.

### Incorporation of genotype data in genetic mapping

For each analyzed DNA variant, we compared isolates (irrespective of the ORF from which they were obtained) carrying the W303 allele to isolates carrying the BY allele as determined by the sequencing data. Bonferroni correction was performed for the number of analyzed DNA variants. To additionally gauge significance, we performed 1,000 random permutations of phenotype data, extracted the best p-value from each permutation, and used the 95% percentile of these best p-values as a permutation-based significance threshold.

In our initial analyses, we noticed that the *sal1-1* variant (variant position 472575) showed substantially fewer isolates with allele calls compared to neighboring variants, likely due to the difficulty of calling isolate genotypes from low-coverage sequencing data at this indel variant. These missing allele calls reduced the sample size available at this specific variant, reducing power and worsening the initially observed p-values. To address this, we manually inspected sequencing read alignments to call the *sal1-1* variant as BY or W303 in 13 local and 12 distal isolates in which the repair event was near the *sal1-1* variant. For distal isolates with repair occurring clearly on the centromeric side of the *sal1-1* variant, this variant was called as W303. For the remaining isolates, we manually added BY allele calls at this position. We acknowledge that this manual curation at *sal1-1* but not other variants could have led to a slightly higher statistical power at this site.

### Engineering of *MKT1* & *SAL1* alleles

We created different combinations of *MKT1* and *SAL1* alleles in the BY4742 background by engineering the region on ChrXIV-L that includes *MKT1*, *END3* and *SAL1* with double-cut CRISPR-Swap (Lutz et al. 2019). First, we replaced *MKT1* with a KanMX cassette to create YFA1822 by amplifying the KanMX fragment from YFA0007 with OFA1283 and OFA1284 and transforming it into BY4742 (YFA1608). The presence of KanMX replacing the *MKT1* ORF was verified by colony PCR with OFA1365 and OFA0036 and the absence of the *MKT1* allele with OFA1365 and OFA1366. Next, we replaced *SAL1* with a HphMX cassette by amplifying the HphMX cassette from YLK1950/YFA0006 with OFA1471 and OFA1472 and transforming it into BY4742 mkt1Δ::kanMX (YFA1822). The presence of HphMX replacing *SAL1* was verified by colony PCR with OFA1473 and OFA0036 and the absence of *SAL1* with OFA1473 and OFA1474. The BY4742 mkt1Δ::kanMX sal1Δ::hphMX (YFA1831) strain was then transformed with ∼100 ng of the CRISPR-Swap plasmid (pFA0055) and ∼1 µg of PCR generated alleles of the 6,275-bp *MKT1*, *END3* and *SAL1* locus.

The allele fragments were created using splicing overlap extension PCR as follows (Horton et al. 1989). The BY and W303 alleles were amplified from YFA1608 and YFA1718 genomic DNA, respectively using primers OFA1365 and OFA1475. MKT1-30G was amplified from W303 DNA with OFA1497 and OFA1366 and BY gDNA with OFA1498 and OFA1501. SAL1-nfs was amplified from BY DNA with OFA1497 and OFA1499 and from W303 DNA with OFA1500 and OFA1501. MKT1-30G+SAL1-nfs was amplified from W303 DNA with OFA1497 and OFA1366; and OFA1500 and OFA1501 and from BY DNA with OFA1498 and OFA1499. An additional BY allele was created as described for the MKT1-30G+SAL1-nfs allele, except the template was BY for each of the three reactions.

Amplicons were run on a 1% agarose gel and the fragments were purified by gel extraction. To create the final alleles, the fragments were combined by amplifying with OFA1365 and OFA1475. All reactions were run on a 1% agarose gel and purified prior to transformation.

### Phenotyping on solid agar plates

Isolates from the *MKT1-END3-SAL1* locus engineering were pinned, randomized and colony sizes measured using a ROTOR+ pinning robot and a PIXL® instrument (Singer Instruments). Details regarding instrument settings and consistent PlusPlate (Singer Instruments) preparation are described below.

Glycerol stocks of the engineered *MKT1-END3-SAL1* isolates were thawed and pinned from glycerol stock plates to YPD agar PlusPlates using the ROTOR+ to create the source plates. The strains on each plate were then randomized using the PIXL Re-Array workflow onto new YPD PlusPlates to reduce the impact of edge and plate effects. Two unique random arrays were created from each source plate. The randomization uses Well-Plate Maker (WPM) (Borges et al. 2021) and additional corrections to avoid the following: colonies of the same strain being neighbors, wild types being unevenly distributed across different plates, a given strain being on an edge position in both random arrays, and any one strain being on edge positions more frequently than any other strain. These random array plates were then used to inoculate cultures for glycerol stocks.

The glycerol stock plates containing the randomized arrays were thawed and pinned using the ROTOR+ to 96 density YPD agar PlusPlates to create “pre-activation plates”, which were incubated at 30 °C for 18 – 36 hours. Pre-activation plates were stored at 4 °C for up to 3 days before use. Pre-activation plates were pinned using the ROTOR+ to 384 density PlusPlates containing YPD medium to create “activation plates”, which were incubated at 30 °C for 20 – 24 hours. Immediately after incubation, activation plates were pinned using the ROTOR+ to 384 density PlusPlates containing the conditional medium to create “phenotyping plates”, which were incubated at 30 °C for exactly 40 hours. Immediately after incubation, the phenotyping plates were analyzed using the PIXL.

#### PlusPlate preparation

We used the following procedure for the preparation of PlusPlates for use in the ROTOR+ and PIXL to ensure the plates were level and uniform. To prepare the media, yeast extract (1%) and peptone (2%) were dissolved in deionized water. This YP medium was divided into bottles with stir bars and autoclaved for 30 minutes. Immediately after autoclaving, 20% glucose solution (2%) was added to each bottle along with non-heat sensitive additives and the bottles were shaken to mix. Heat sensitive additives were added after YPD cooled to 60 °C and mixed by stirring. Media was allowed to cool in a 53 °C incubator for approximately 2 hours before plates were filled.

To fill the PlusPlates, a 1 L bottle with 500 mL of 53 °C molten agar media was put on a stir/hot plate set to low heat and was stirred without producing bubbles. PlusPlates were laid out on a level surface. A 50-mL serological pipette was used to draw 55 mL of media, and exactly 50 mL were dispensed into each plate. The remaining 5 mL were dispensed back into the bottle. These steps were repeated until all media was used. The filled plates were allowed to cool and dry for 18 to 24 hours before either being used or they were inverted and bagged for longer term storage. Plates were not used after being stored for more than three weeks.

#### Yeast pinning

Plates were pinned using the ROTOR+. Plates were pinned from 96 density liquid arrays to 96 density solid arrays with 96 density long RePads (Singer Instruments) using default settings except the following: 10% target pin pressure, 2.25 mm target overshoot, 1 wet mix rotation cycle. Plates were pinned from 96 density solid arrays to 384 density solid arrays with 96 density short RePads using default settings except the following: no recycle, automatic offset, 15% target pin pressure, 2.25 mm target overshoot. Plates were pinned from 384 density solid arrays to 384 density solid arrays with 384 density short RePads using default settings except the following: automatic offset, 2.25 mm target overshoot.

#### Measuring colony size

The PIXL’s Colony Detection protocol was used to photograph the plates and perform image analysis to identify yeast colonies and measure colony sizes. A Colony Detection template was used with the following settings: Background Subtraction algorithm, *S. cerevisiae* species, and 0.5-3.0 mm radius filter. The PIXL’s blackout screen was inserted prior to image acquisition. If colonies were too small to be detected using the Background Subtraction algorithm but larger than 0.5 mm, the process was repeated using the Colony Separation algorithm. Raw image analysis data from the PIXL was processed so that each colony was matched to its strain information. The output from these procedures was the area of each colony, measured in mm^2^.

#### Correction for edge and plate effects

Colonies at the edge of solid agar plates grew larger than those in the center of the plate, a well-known phenomenon in solid plate-based phenotypic assays (Wagih et al. 2013). Our randomization strategy (see above) was designed to avoid confounding of genotype with plate position. In addition, we corrected colony sizes for colony position on the plate using the following mixed linear model:

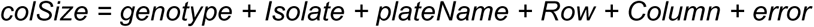

In this model, we used colony size as the dependent variable, modeled as the sum of a series of random effects including the row and column in which a colony was located, the plate a colony was on (to control for systematic differences between plates). Rows and columns were modeled as categorical factors. To fully represent the data during pre-processing, the model also included random effect terms for the genotype (i.e. the introduced DNA fragment in the validation experiments) and isolate of a given colony. We obtained corrected colony sizes by subtracting the effects of *plateName*, *Row*, and *Column* from the raw measurements and used these residuals in further analyses.

### Statistical analysis of engineered *MKT1* & *SAL1* strains

We performed comparisons of a standard environmental condition (colony size on solid YPD after 40 hours of growth) and one other environment at a time. We analyzed the effects of the causal variants in *MKT1* and *SAL1* using the following linear model:

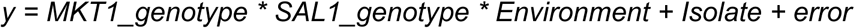

Here, *y* denotes the phenotype (liquid growth rates or colony sizes), *MKT1_genotype* and *SAL1_genotype* are fixed effects denoting whether a given isolate carried the BY or W303 allele at the *MKT1*-D30G or *sal1-1* variant, *Environment* denotes a fixed effect for the environment the measurements were taken in (either the standard condition or the given other condition), and *Isolate* is a random effect denoting isolate identity. The model included all interaction terms between the fixed effects, allowing simultaneous determination of the marginal effects of each individual allele, GxG between alleles, GxE for each allele, and the global GxGxE term that asks if the pattern of epistasis (GxG) depends on the environment. Statistical significance of the GxGxE term and of the GxE terms for each allele was gauged using ANOVA of the above model.

Statistical significance of the GxG term was assessed using models of one condition at a time:

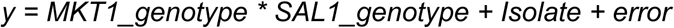

Marginal effects of each allele and of multi-allele constructs compared to the wildtype BY strain were tested as described in the section “ORF-based genetic mapping” above. No multiple-test correction was performed for these analyses. Instead, raw p-values were reported and interpreted keeping in mind the exploratory nature of these GxE experiments.

## Acknowledgements

We thank Robert Reid for detailed technical advice and for providing the W303 SPA strain, David Kirkpatrick for sharing a copy of the YKO collection, Gloria Baxter for assistance with strain construction, and Kevin Zhan and Samantha Graham for critical reading of the manuscript. We thank the University of Minnesota Genomics Center for Illumina sequencing and technical support.

## References

Abyzov A, Urban AE, Snyder M, Gerstein M. 2011. CNVnator: An approach to discover, genotype, and characterize typical and atypical CNVs from family and population genome sequencing. Genome Res 21: 974–984.

Albert FW, Treusch S, Shockley AH, Bloom JS, Kruglyak L. 2014. Genetics of single-cell protein abundance variation in large yeast populations. Nature 506: 494–497.

Anand RP, Lovett ST, Haber JE. 2013. Break-Induced DNA Replication. Cold Spring Harbor Perspectives in Biology 5: a010397–a010397.

Bankaitis VA, Johnson LM, Emr SD. 1986. Isolation of yeast mutants defective in protein targeting to the vacuole. Proc Natl Acad Sci USA 83: 9075–9079.

Bao Z, HamediRad M, Xue P, Xiao H, Tasan I, Chao R, Liang J, Zhao H. 2018. Genome-scale engineering of Saccharomyces cerevisiae with single-nucleotide precision. Nature Biotechnology 36: 505–508.

Berger KH, Yaffe MP. 2000. Mitochondrial DNA inheritance in Saccharomyces cerevisiae. Trends Microbiol 8: 508–513.

Bloom JS, Boocock J, Treusch S, Sadhu MJ, Day L, Oates-Barker H, Kruglyak L. 2019. Rare variants contribute disproportionately to quantitative trait variation in yeast. eLife 8: e49212.

Bloom JS, Kotenko I, Sadhu MJ, Treusch S, Albert FW, Kruglyak L. 2015. Genetic interactions contribute less than additive effects to quantitative trait variation in yeast. Nature Communications 6: 8712.

Borges H, Hesse A-M, Kraut A, Couté Y, Brun V, Burger T. 2021. Well Plate Maker: a user-friendly randomized block design application to limit batch effects in large-scale biomedical studies ed. J. Kelso. Bioinformatics 37: 2770–2771.

Brion C, Lutz SM, Albert FW. 2020. Simultaneous quantification of mRNA and protein in single cells reveals post-transcriptional effects of genetic variation. eLife 9: e60645.

Caballero D, Sutter BM, Xing Z, Wang C, Choo E, Wang Y, Yang Y-S, Ghaemmaghami S, Lemoff A, Tu BP. 2025. The yeast Mkt1/Pbp1 complex promotes adaptive responses to respiratory growth. Journal of Cell Biology 224: e202411169.

Cachera P, Olsson H, Coumou H, Jensen ML, Sánchez BJ, Strucko T, van den Broek M, Daran J-M, Jensen MK, Sonnenschein N, et al. 2023. CRI-SPA: a high-throughput method for systematic genetic editing of yeast libraries. Nucleic Acids Research 51: e91–e91.

Camacho C, Coulouris G, Avagyan V, Ma N, Papadopoulos J, Bealer K, Madden TL. 2009. BLAST+: architecture and applications. BMC Bioinformatics 10: 421.

Chen S-AA, Kern AF, Ang RML, Xie Y, Fraser HB. 2023. Gene-by-environment interactions are pervasive among natural genetic variants. Cell Genomics 3: 100273.

Chen XJ. 2004. Sal1p, a Calcium-Dependent Carrier Protein That Suppresses an Essential Cellular Function Associated With the Aac2 Isoform of ADP/ATP Translocase in Saccharomyces cerevisiae. Genetics 167: 607–617.

Cubillos FA, Parts L, Salinas F, Bergström A, Scovacricchi E, Zia A, Illingworth CJR, Mustonen V, Ibstedt S, Warringer J, et al. 2013. High-Resolution Mapping of Complex Traits with a Four-Parent Advanced Intercross Yeast Population. Genetics 195: 1141–1155.

Danecek P, Bonfield JK, Liddle J, Marshall J, Ohan V, Pollard MO, Whitwham A, Keane T, McCarthy SA, Davies RM, et al. 2021. Twelve years of SAMtools and BCFtools. GigaScience 10: giab008.

Demogines A, Smith E, Kruglyak L, Alani E. 2008. Identification and Dissection of a Complex DNA Repair Sensitivity Phenotype in Baker’s Yeast. PLoS Genetics 4: e10000123.

Deutschbauer AM, Davis RW. 2005. Quantitative trait loci mapped to single-nucleotide resolution in yeast. Nature Genetics.

DiCarlo JE, Norville JE, Mali P, Rios X, Aach J, Church GM. 2013. Genome engineering in Saccharomyces cerevisiae using CRISPR-Cas systems. Nucleic Acids Research 41: 4336–4343.

Dimitrov LN, Brem RB, Kruglyak L, Gottschling DE. 2009. Polymorphisms in Multiple Genes Contribute to the Spontaneous Mitochondrial Genome Instability of Saccharomyces cerevisiae S288C Strains. Genetics 183: 365–383.

Ehrenreich IM, Torabi N, Jia Y, Kent J, Martis S, Shapiro JA, Gresham D, Caudy AA, Kruglyak L. 2010. Dissection of genetically complex traits with extremely large pools of yeast segregants. Nature 464: 1039–1042.

Engel SR, Aleksander S, Nash RS, Wong ED, Weng S, Miyasato SR, Sherlock G, Cherry JM. 2025. *Saccharomyces* Genome Database: advances in genome annotation, expanded biochemical pathways, and other key enhancements ed. S. Lacefield. GENETICS 229: iyae185.

Fay JC. 2013. The molecular basis of phenotypic variation in yeast. Current opinion in genetics & development 23: 672–677.

Fritsch ES, Chabbert CD, Klaus B, Steinmetz LM. 2014. A Genome-Wide Map of Mitochondrial DNA Recombination in Yeast. Genetics 198: 755–771.

Gaisne M, Bécam AM, Verdiere J, Herbert CJ. 1999. A “natural” mutation in Saccharomyces cerevisiae strains derived from S288c affects the complex regulatory gene HAP1 (CYP1). Current Genetics 36: 195–200.

Gerke J, Lorenz K, Cohen B. 2009. Genetic Interactions Between Transcription Factors Cause Natural Variation in Yeast. Science 323: 498–501.

Giaever G, Chu AM, Ni L, Connelly C, Riles L, Véronneau S, Dow S, Lucau-Danila A, Anderson K, André B, et al. 2002. Functional profiling of the Saccharomyces cerevisiae genome. Nature 418: 387–391.

Gietz RD, Schiestl RH. 2007. High-efficiency yeast transformation using the LiAc/SS carrier DNA/PEG method. Nature Protocols 2: 31–34.

Gorter de Vries AR, Couwenberg LGF, van den Broek M, de la Torre Cortés P, Ter Horst J, Pronk JT, Daran J-MG. 2019. Allele-specific genome editing using CRISPR-Cas9 is associated with loss of heterozygosity in diploid yeast. Nucleic Acids Res 47: 1362–1372.

Horton RM, Hunt HD, Ho SN, Pullen JK, Pease LR. 1989. Engineering hybrid genes without the use of restriction enzymes: gene splicing by overlap extension. Gene 77: 61–68.

Kataoka T, Powers S, McGill C, Fasano O, Strathern J, Broach J, Wigler M. 1984. Genetic analysis of yeast RAS1 and RAS2 genes. Cell 37: 437–445.

Kurtz S, Phillippy A, Delcher AL, Smoot M, Shumway M, Antonescu C, Salzberg SL. 2004. Versatile and open software for comparing large genomes. Genome Biol 5: R12.

Lappalainen T, Li YI, Ramachandran S, Gusev A. 2024. Genetic and molecular architecture of complex traits. Cell 187: 1059–1075.

Lee S-I, Dudley AM, Drubin D, Silver PA, Krogan NJ, Pe’er D, Koller D. 2009. Learning a Prior on Regulatory Potential from eQTL Data. PLoS Genetics 5: e1000358.

Lehner KR, Stone MM, Farber RA, Petes TD. 2007. Ninety-Six Haploid Yeast Strains With Individual Disruptions of Open Reading Frames Between *YOR097C* and *YOR192C*, Constructed for the *Saccharomyces* Genome Deletion Project, Have an Additional Mutation in the Mismatch Repair Gene *MSH3*. Genetics 177: 1951–1953.

Li H. 2013. Aligning sequence reads, clone sequences and assembly contigs with BWA-MEM. https://arxiv.org/abs/1303.3997 (Accessed August 28, 2025).

Liti G, Louis EJ. 2012. Advances in Quantitative Trait Analysis in Yeast ed. J.C. Fay. PLoS Genet 8: e1002912.

Lutz S, Brion C, Kliebhan M, Albert FW. 2019. DNA variants affecting the expression of numerous genes in trans have diverse mechanisms of action and evolutionary histories ed. J.C. Fay. PLoS Genet 15: e1008375.

Lutz S, Van Dyke K, Feraru MA, Albert FW. 2022. Multiple epistatic DNA variants in a single gene affect gene expression in *trans* ed. P. Wittkopp. Genetics 220: iyab208.

Lynch M, Walsh B. 1998. Genetics and analysis of quantitative traits. Sinauer Associates, Inc.

Mackay TFC, Anholt RRH. 2024. Pleiotropy, epistasis and the genetic architecture of quantitative traits. Nat Rev Genet 25: 639–657.

Malcova I, Farkasovsky M, Senohrabkova L, Vasicova P, Hasek J. 2016. New integrative modules for multicolor-protein labeling and live-cell imaging in Saccharomyces cerevisiae. FEMS Yeast Res 16.

Matheson K, Parsons L, Gammie A. 2017. Whole-Genome Sequence and Variant Analysis of W303, a Widely-Used Strain of *Saccharomyces cerevisiae*. G3 Genes|Genomes|Genetics 7: 2219–2226.

McLaren W, Gil L, Hunt SE, Riat HS, Ritchie GRS, Thormann A, Flicek P, Cunningham F. 2016. The Ensembl Variant Effect Predictor. Genome Biology 17.

Mulugu S, Bai W, Fridy PC, Bastidas RJ, Otto JC, Dollins DE, Haystead TA, Ribeiro AA, York JD. 2007. A Conserved Family of Enzymes That Phosphorylate Inositol Hexakisphosphate. Science 316: 106–109.

Nguyen Ba AN, Lawrence KR, Rego-Costa A, Gopalakrishnan S, Temko D, Michor F, Desai MM. 2022. Barcoded bulk QTL mapping reveals highly polygenic and epistatic architecture of complex traits in yeast. eLife 11: e73983.

Peter J, Chiara MD, Friedrich A, Yue J-X, Pflieger D, Bergström A, Sigwalt A, Barre B, Freel K, Llored A, et al. 2018. Genome evolution across 1,011 Saccharomyces cerevisiae isolates. Nature 556: 339–344.

Peter J, Friedrich A, Liti G, Schacherer J. 2022. Extensive simulations assess the performance of genome-wide association mapping in various *Saccharomyces cerevisiae* subpopulations. Phil Trans R Soc B 377: 20200514.

R Core Team. 2025. R: A Language and Environment for Statistical Computing. R Foundation for Statistical Computing, Vienna, Austria https://www.R-project.org.

Reid RJD, González-Barrera S, Sunjevaric I, Alvaro D, Ciccone S, Wagner M, Rothstein R. 2011. Selective ploidy ablation, a high-throughput plasmid transfer protocol, identifies new genes affecting topoisomerase I–induced DNA damage. Genome Res 21: 477–486.

Reid RJD, Sunjevaric I, Voth WP, Ciccone S, Du W, Olsen AE, Stillman DJ, Rothstein R. 2008. Chromosome-Scale Genetic Mapping Using a Set of 16 Conditionally Stable *Saccharomyces cerevisiae* Chromosomes. Genetics 180: 1799–1808.

Roy KR, Smith JD, Vonesch SC, Lin G, Tu CS, Lederer AR, Chu A, Suresh S, Nguyen M, Horecka J, et al. 2018. Multiplexed precision genome editing with trackable genomic barcodes in yeast. Nature Biotechnology 36: 512–520.

Sadhu MJ, Bloom JS, Day L, Kruglyak L. 2016. CRISPR-directed mitotic recombination enables genetic mapping without crosses. Science 352: 1113–1116.

Sadhu MJ, Bloom JS, Day L, Siegel JJ, Kosuri S, Kruglyak L. 2018. Highly parallel genome variant engineering with CRISPR–Cas9. Nat Genet 50: 510–514.

Saha D, Gregor JB, Hoda S, Eastman KE, Gutierrez-Schultz VA, Navarrete M, Wisecaver JH, Briggs SD. 2024. Candida glabrata maintains two HAP1 ohnologs, HAP1A and HAP1B, for distinct roles in ergosterol gene regulation to mediate sterol homeostasis under azole and hypoxic conditions. mSphere 9: e0052424.

Schell R, Hale JJ, Mullis MN, Matsui T, Foree R, Ehrenreich IM. 2022. Genetic basis of a spontaneous mutation’s expressivity ed. C.L. Peichel. Genetics 220: iyac013.

Sharon E, Chen S-AA, Khosla NM, Smith JD, Pritchard JK, Fraser HB. 2018. Functional Genetic Variants Revealed by Massively Parallel Precise Genome Editing. Cell 175: 544–557.e16.

She R, Jarosz DF. 2018. Mapping Causal Variants with Single-Nucleotide Resolution Reveals Biochemical Drivers of Phenotypic Change. Cell 172: 478–490.e15.

Sinha H, Nicholson BP, Steinmetz LM, McCusker JH. 2006. Complex Genetic Interactions in a Quantitative Trait Locus ed. J. Haber. PLoS Genet 2: e13.

Song G, Dickins BJA, Demeter J, Engel S, Dunn B, Cherry JM. 2015. AGAPE (Automated Genome Analysis PipelinE) for Pan-Genome Analysis of Saccharomyces cerevisiae ed. J. Schacherer. PLoS ONE 10: e0120671.

Sprouffske K, Wagner A. 2016. Growthcurver: an R package for obtaining interpretable metrics from microbial growth curves. BMC Bioinformatics 17: 172.

Steinmetz LM, Sinha H, Richards DR, Spiegelman JI, Oefner PJ, McCusker JH, Davis RW. 2002. Dissecting the architecture of a quantitative trait locus in yeast. Nature 416: 326–330.

Tange O. 2022. GNU Parallel 2022-08-22 (Rushdie’). https://zenodo.org/record/7015730 (Accessed August 28, 2025).

Teng X, Dayhoff-Brannigan M, Cheng W-C, Gilbert CE, Sing CN, Diny NL, Wheelan SJ, Dunham MJ, Boeke JD, Pineda FJ, et al. 2013. Genome-wide Consequences of Deleting Any Single Gene. Molecular Cell 52: 485–494.

Traba J, Froschauer EM, Wiesenberger G, Satrústegui J, Del Arco A. 2008. Yeast mitochondria import ATP through the calcium-dependent ATP-Mg/Pi carrier Sal1p, and are ATP consumers during aerobic growth in glucose. Molecular Microbiology 69: 570–585.

Tschopp JF, Emr SD, Field C, Schekman R. 1986. GAL2 codes for a membrane-bound subunit of the galactose permease in Saccharomyces cerevisiae. J Bacteriol 166: 313–318.

Van Leeuwen J, Pons C, Mellor JC, Yamaguchi TN, Friesen H, Koschwanez J, Ušaj MM, Pechlaner M, Takar M, Ušaj M, et al. 2016. Exploring genetic suppression interactions on a global scale. Science 354: aag0839.

Visscher PM, Brown MA, McCarthy MI, Yang J. 2012. Five Years of GWAS Discovery. The American Journal of Human Genetics 90: 7–24.

Wagih O, Usaj M, Baryshnikova A, VanderSluis B, Kuzmin E, Costanzo M, Myers CL, Andrews BJ, Boone CM, Parts L. 2013. SGAtools: one-stop analysis and visualization of array-based genetic interaction screens. Nucleic Acids Research 41: W591–W596.

Weiss CV, Roop JI, Hackley RK, Chuong JN, Grigoriev IV, Arkin AP, Skerker JM, Brem RB. 2018. Genetic dissection of interspecific differences in yeast thermotolerance. Nat Genet 50: 1501–1504.

Wilkening S, Lin G, Fritsch ES, Tekkedil MM, Anders S, Kuehn R, Nguyen M, Aiyar RS, Proctor M, Sakhanenko NA, et al. 2014. An Evaluation of High-Throughput Approaches to QTL Mapping in *Saccharomyces cerevisiae*. Genetics 196: 853–865.

Winzeler EA, Shoemaker DD, Astromoff A, Liang H, Anderson K, Andre B, Bangham R, Benito R, Boeke JD, Bussey H, et al. 1999. Functional characterization of the S. cerevisiae genome by gene deletion and parallel analysis. Science 285: 901–906.

Yin Y, Dominska M, Yim E, Petes TD. 2017. High-resolution mapping of heteroduplex DNA formed during UV-induced and spontaneous mitotic recombination events in yeast. Elife 6: e28069.

Young AI, Benonisdottir S, Przeworski M, Kong A. 2019. Deconstructing the sources of genotype-phenotype associations in humans. Science 365: 1396–1400.

Zhu J, Zhang B, Smith EN, Drees B, Brem RB, Kruglyak L, Bumgarner RE, Schadt EE. 2008. Integrating large-scale functional genomic data to dissect the complexity of yeast regulatory networks. Nature Genetics 40: 854–861.

Zuo E, Huo X, Yao X, Hu X, Sun Y, Yin J, He B, Wang X, Shi L, Ping J, et al. 2017. CRISPR/Cas9-mediated targeted chromosome elimination. Genome Biol 18: 224.

