## Supplementary Figures for "High-resolution, genotype-free mapping of genetic variation with CRI-SPA-Map"

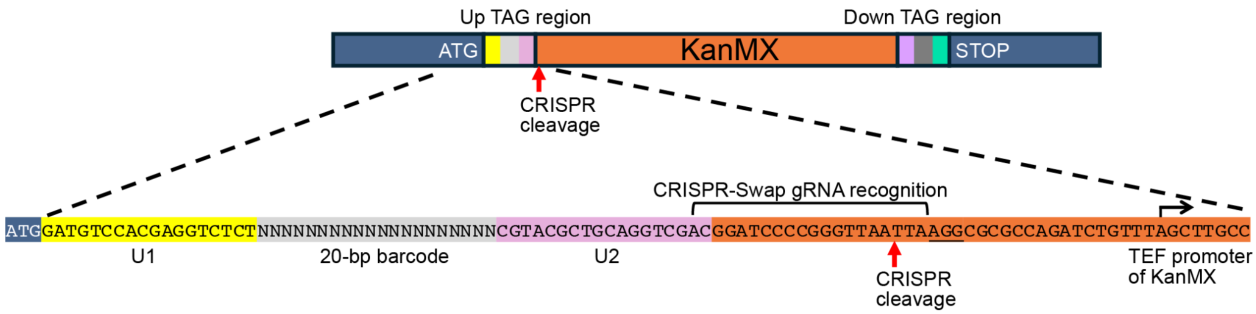


**SFigure1.** The recognition site of the CRISPR-Swap guide RNA in the linker region of the KanMX cassette. The inserted KanMX cassette in each YKO strain contains 20-bp barcodes at each end that are flanked by constant regions (U1 and U2) present in all YKO strains. On the 5′ and 3’ sides of the cleavage site, there are 71 bp and 1510 bp of KanMX cassette sequence, respectively.

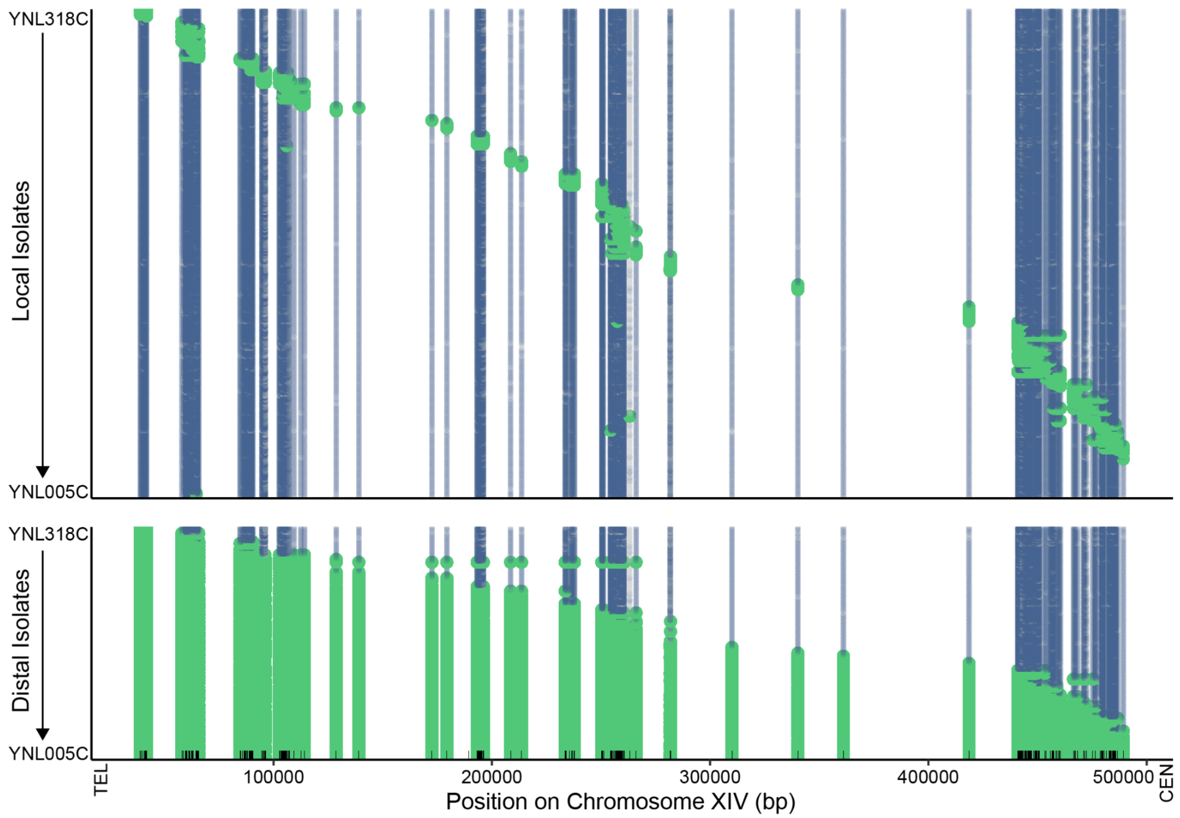


**SFigure 2.** Overview of all repair tracks in the CRI-SPA-Map isolates ordered by targeted ORF. Note the DNA fingerprint variant at 189,585 bp, which segregates in the YKO collection is removed.

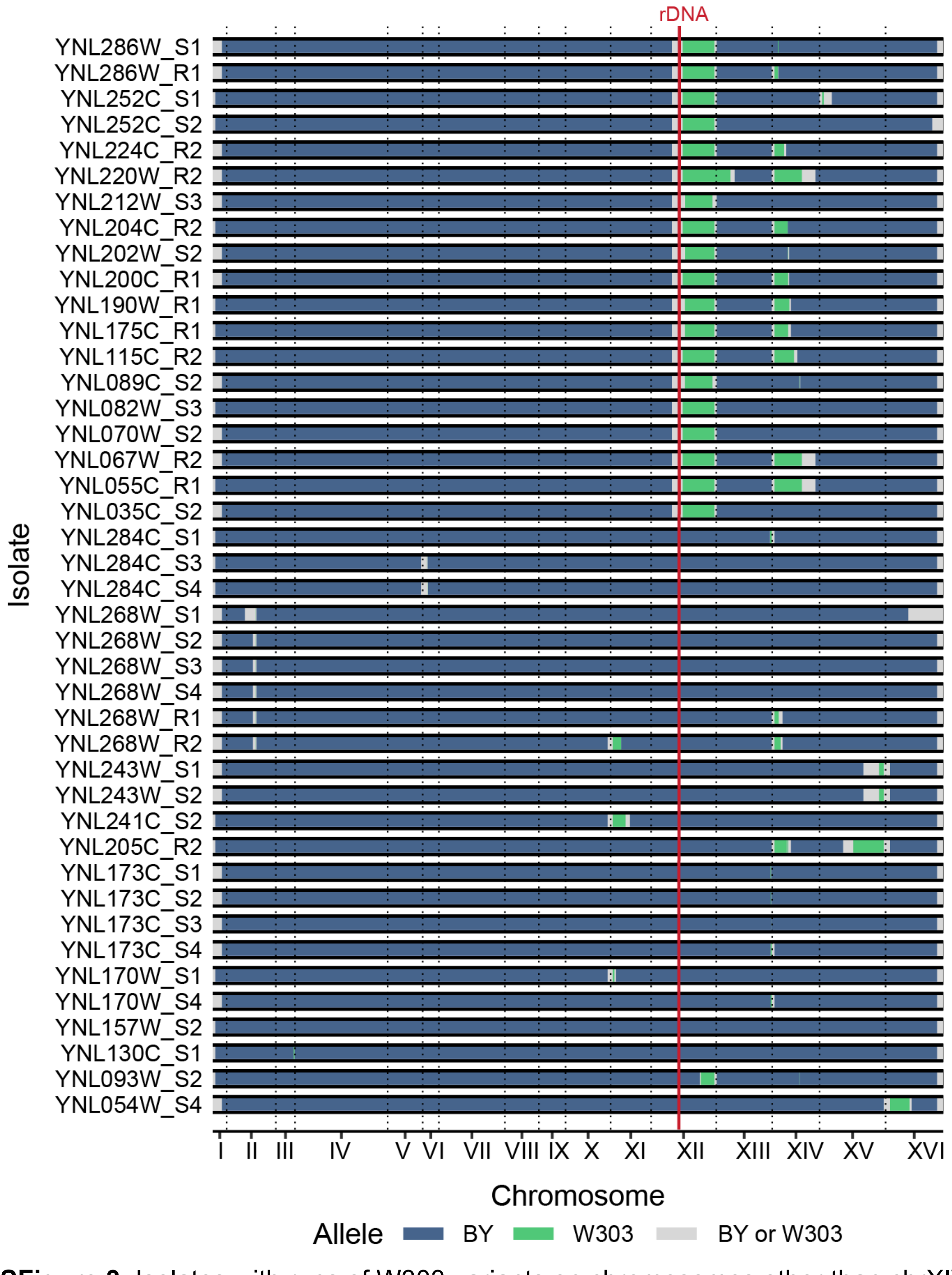


**SFigure 3.** Isolates with runs of W303 variants on chromosomes other than chrXIV-L. The region from the rDNA to the telomere is the most frequently observed location of W303 variant integration other than the targeted chrXIV-L. The isolates mostly come from different YKO strains. The exceptions include all six isolates of the YNL268W YKO strain that have W303 variants on chrII-L due to the presence of the *LYS2* gene in these isolates. All 4 isolates from YNL173C YKO strain (only $Hyg^S$ isolates were recovered) have W303 variants on chrXIII-R suggesting this is the site of the YKO ORF that was inadvertently targeted.

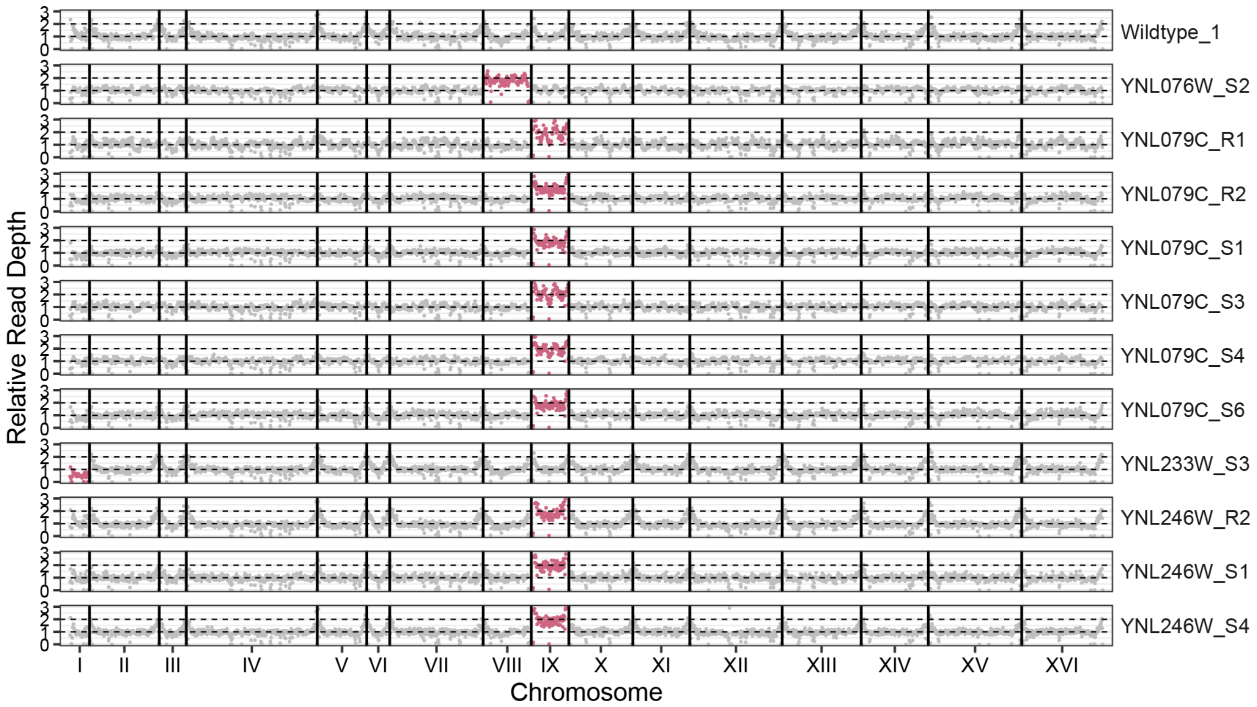


**SFigure 4.** Isolates with aneuploidies. Plotted is the ratio of the read depth in 5,000-bp bins compared to the average read depth across the nuclear genome for each isolate. Chromosomes with detected aneuploidies are colored in pink. Shown for comparison is the wildtype isolate 1, which has no detected aneuploidy.

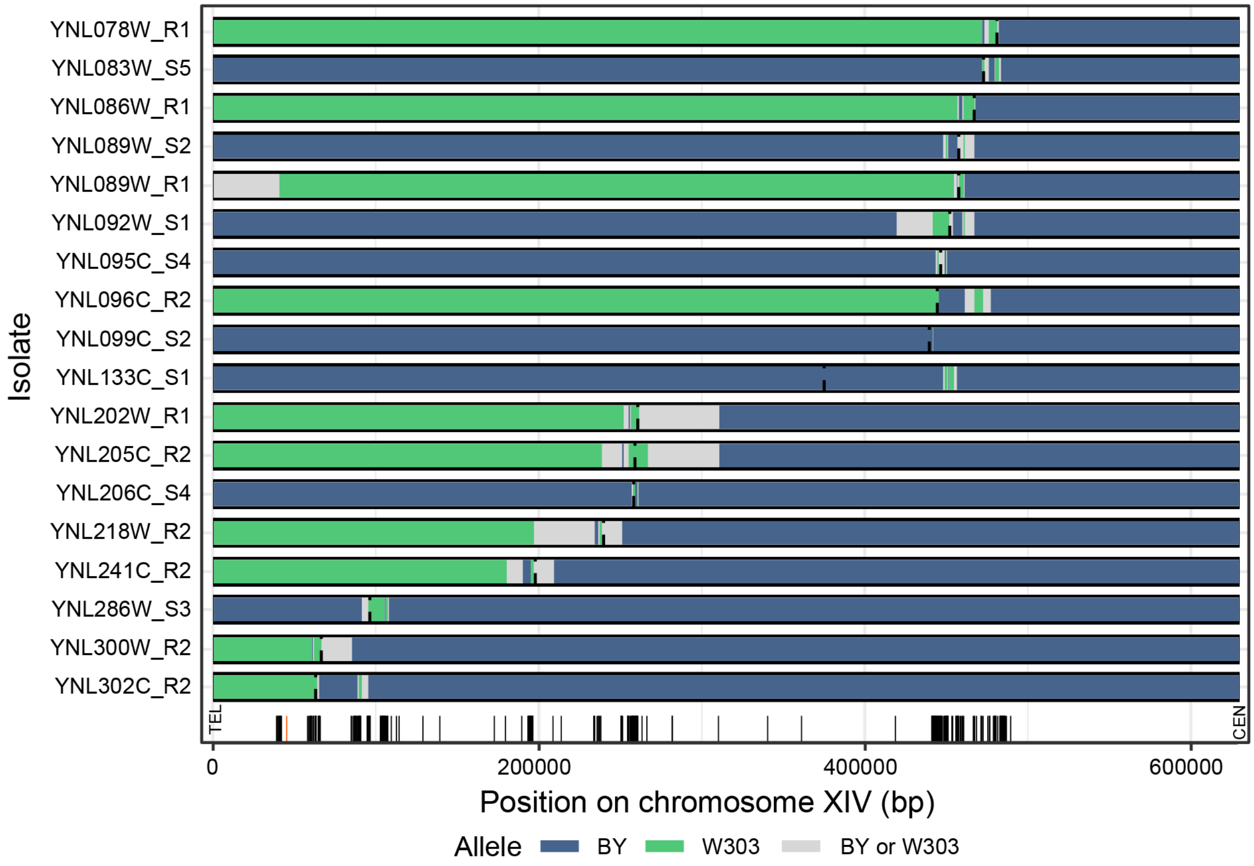


**SFigure 5.** Isolates with mosaic tracts of BY and W303 variants. BY alleles are in blue and W303 alleles in green. The sequence between the last tract of variants to the next tract of variants of the opposite allele could originate from either the BY or W303 strain and is therefore designated as gray. The position of the targeted ORF is shown with a dotted line. The position of the variants are marked by vertical lines along with the position of the telomere (TEL) and centromere (CEN) at the bottom of the plot.

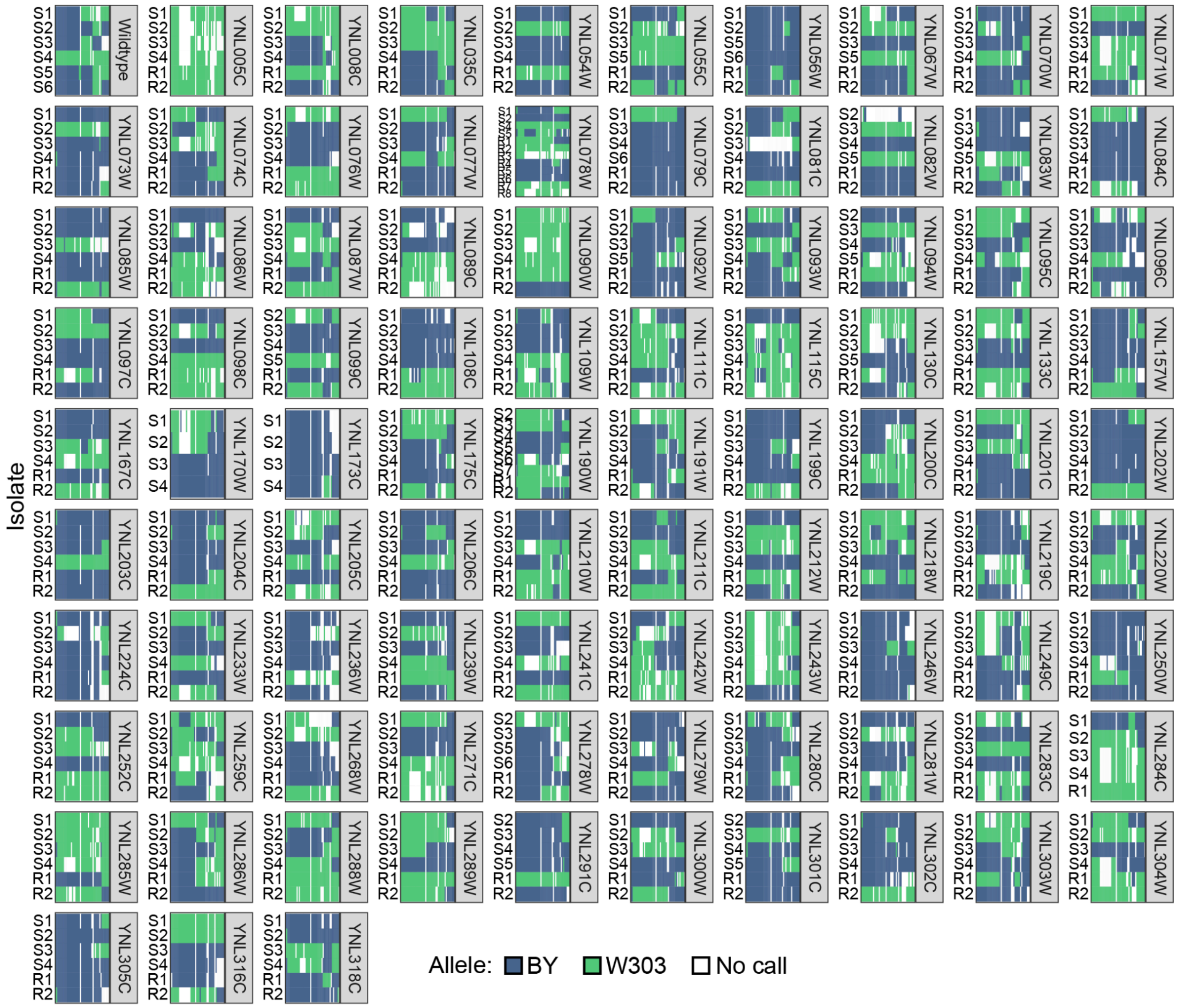


**SFigure 6.** Mitochondrial variants in the CRI-SPA-Map isolates. Variant calls were determined at 48 sites. Isolates from each YKO strain and the wildtype strain had mitochondria with differing genotypes, with the exception of YNL005C. All isolates of YNL005C had only W303 variant calls. YNL005C/*MRP7* encodes a subunit of the mitochondrial ribosome and deletion of this gene causes loss of mitochondria. Therefore, the potential absence of mitochondria in the YNL005C/*MRP7* YKO strain may have led to CRI-SPA-Map isolates with only W303 mitochondria.

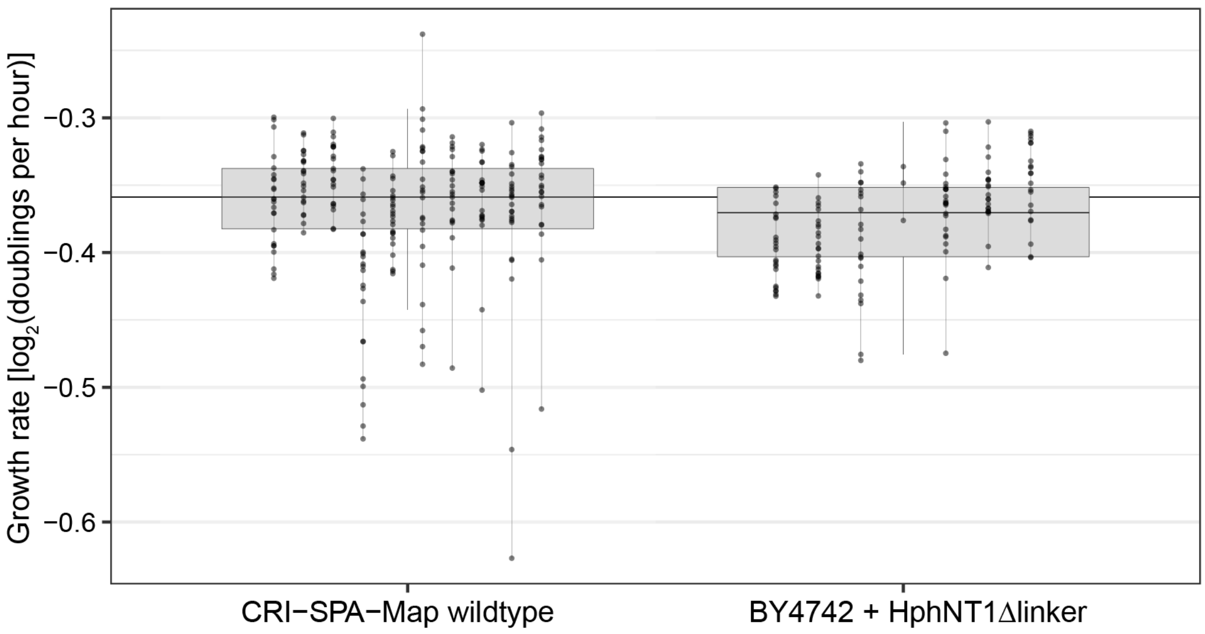


**SFigure 7**. The effect on growth in YPD liquid of the HphMX cassette integrated near the telomere on the left arm of chromosome XIV. The cassette did not have a significant effect on growth rate (T-test p-value = 0.47). Growth measurements from the same isolate are connected by a vertical line. Box plots designate the central 50% of the data with the median marked as a horizontal line within each box.

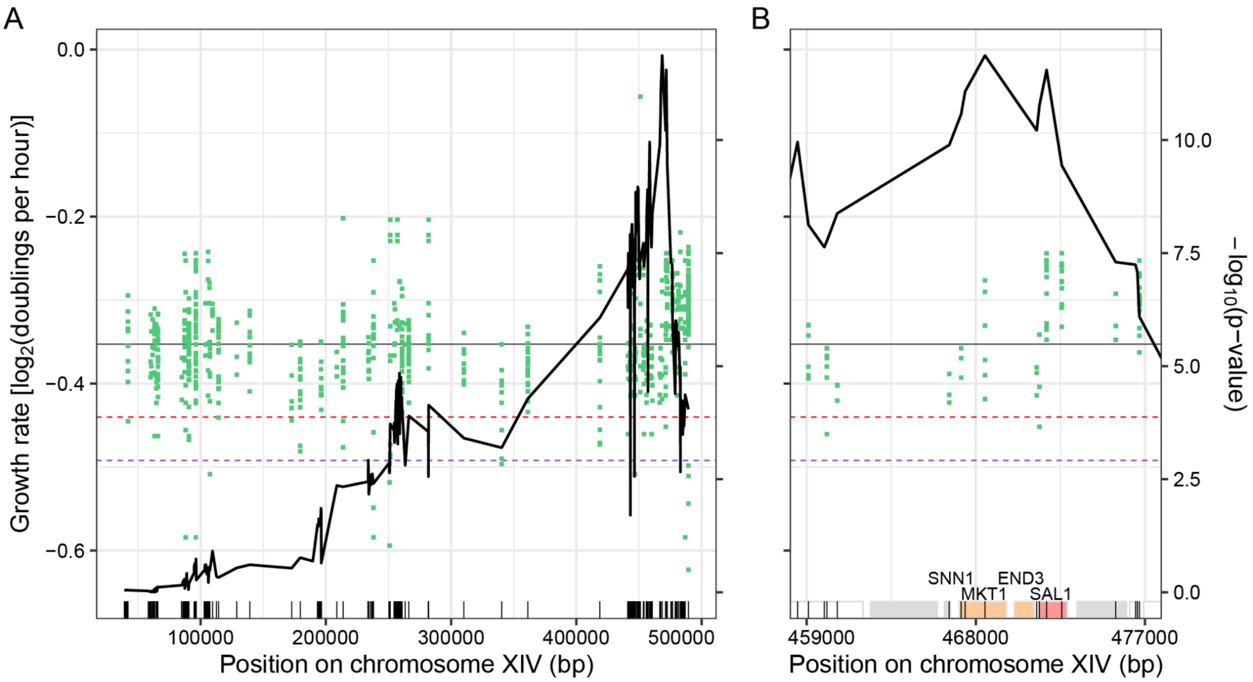


**SFigure 8.** Incorporation of genotype data in CRI-SPA-Map for distal isolates. (A) The variants on the left arm of chromosome XIV are depicted with vertical lines on the bottom of the plot. Green points depict the centromeric end of the W303 allele tracts in each of 163 distal isolates, as well as the growth rate of each isolate in liquid YPD. The median growth rate of all distal isolates is denoted by a horizontal gray line. Dashed lines denote significance thresholds as determined by Bonferroni correction (red) and 1,000 permutations (purple). (B) Zoom-in on the region containing the most significant variants. The ORFs in this region are depicted as boxes and those used in the ORF-based mapping are colored based on the significance of their effect on growth as in Figure 2, bottom panel.

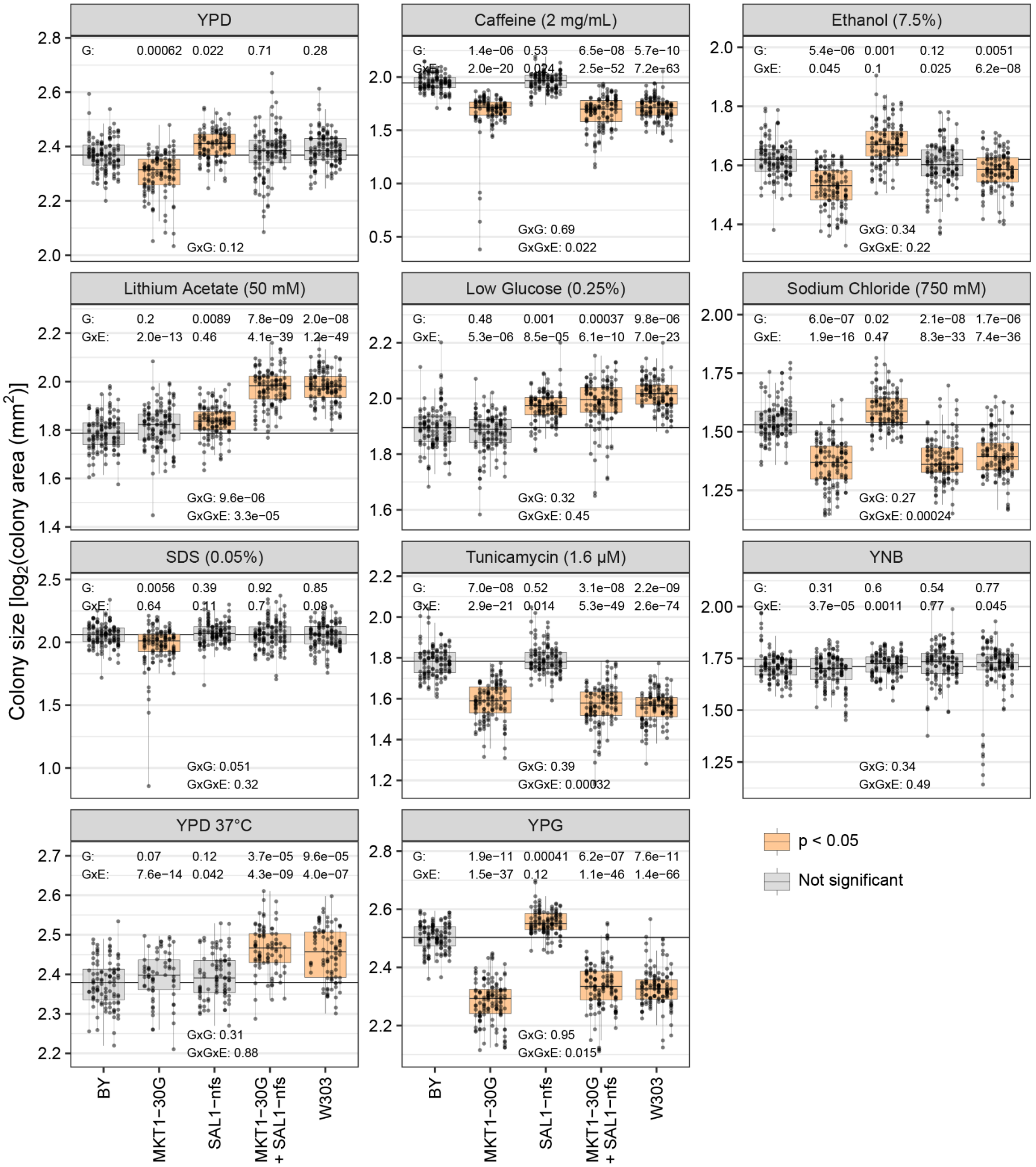


**SFigure 9.** Genetic and environmental interactions of *MKT1-30G* and *SAL1-nfs* alleles. Growth measurements from the same strain are connected by a vertical line. Box plots designate the central 50% of the data with the median marked as a horizontal line within each box. The median of the BY wildtype strain is depicted as a horizontal line through the plot. Box plots are shaded orange if $p < 0.05$. The p values for the various tests of genotype (G) and environmental (E) interactions are shown.
