## Supplementary File 1 for "High-resolution, genotype-free mapping of genetic variation with CRI-SPA-Map"

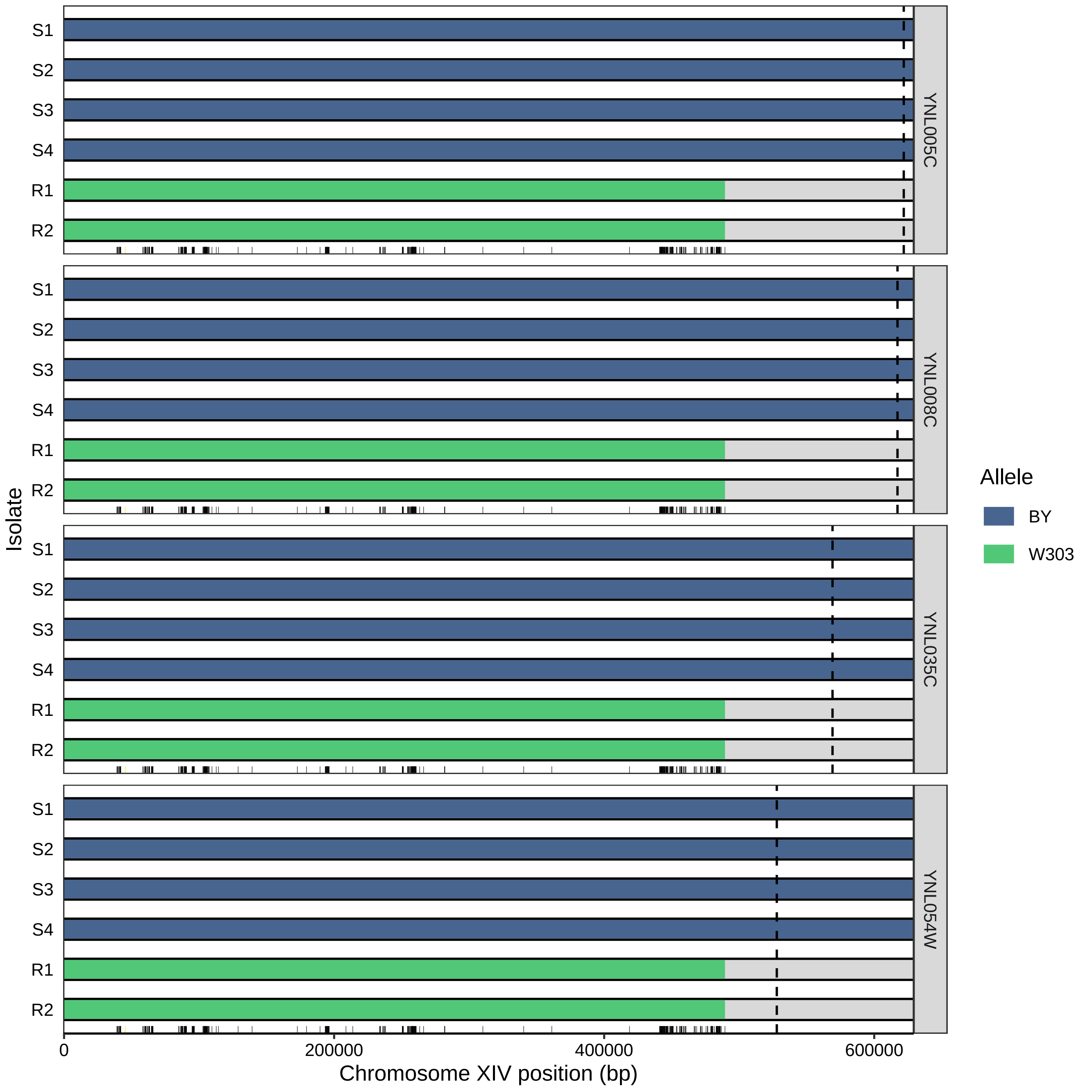
S1
S2
S3
S4
R1
R2
YNL005C
YNL008C
YNL035C
YNL054W
Isolate
Allele
BY
W303
0
200000
400000
600000
Chromosome XIV position (bp)

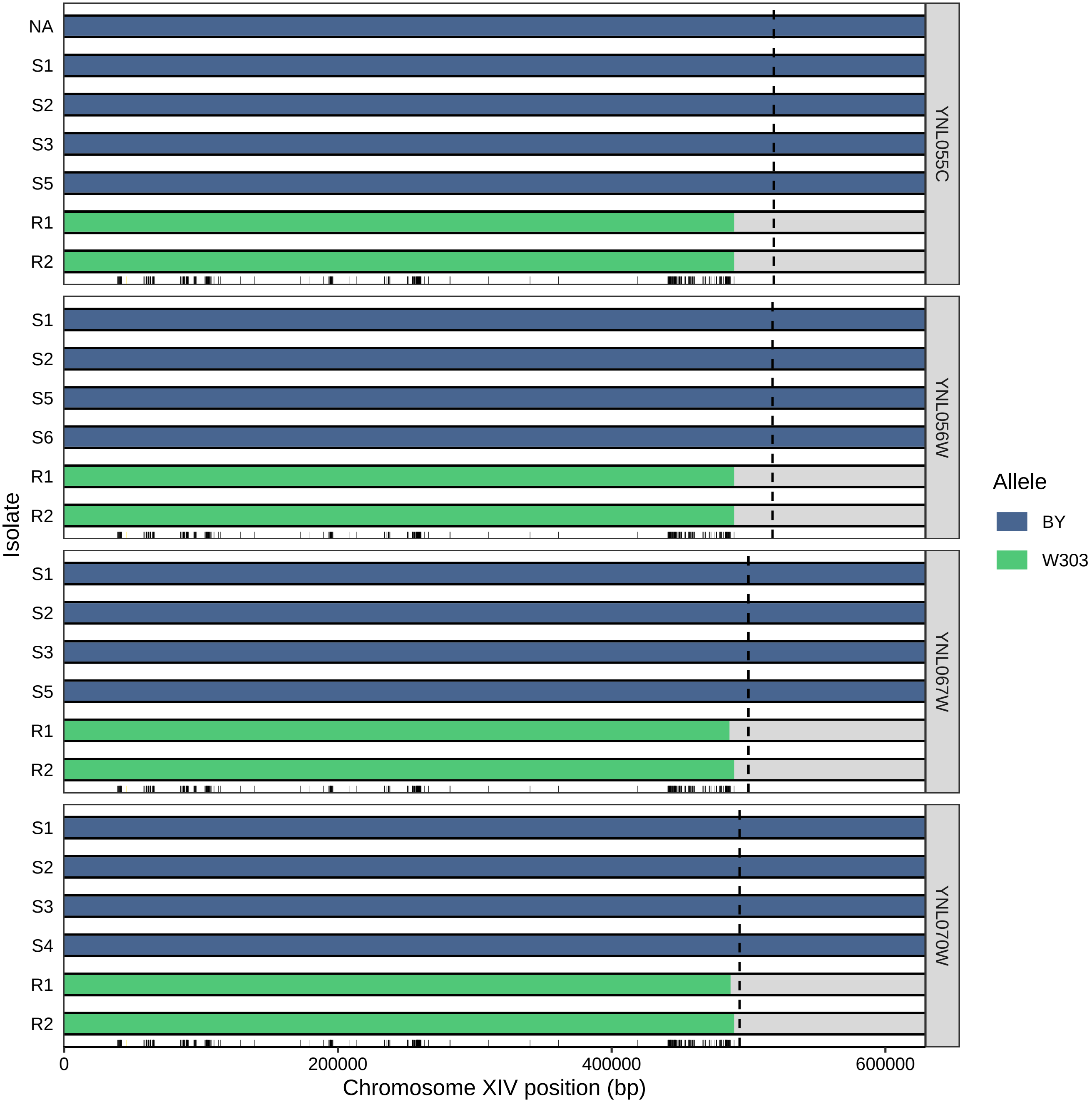

NA
S1
S2
S3
S5
R1
R2
YNL055C
S1
S2
S5
S6
R1
R2
YNL056W
S1
S2
S3
S5
R1
R2
YNL067W
S1
S2
S3
S4
R1
R2
YNL070W
Isolate
0
200000
400000
600000
Chromosome XIV position (bp)
Allele
BY
W303

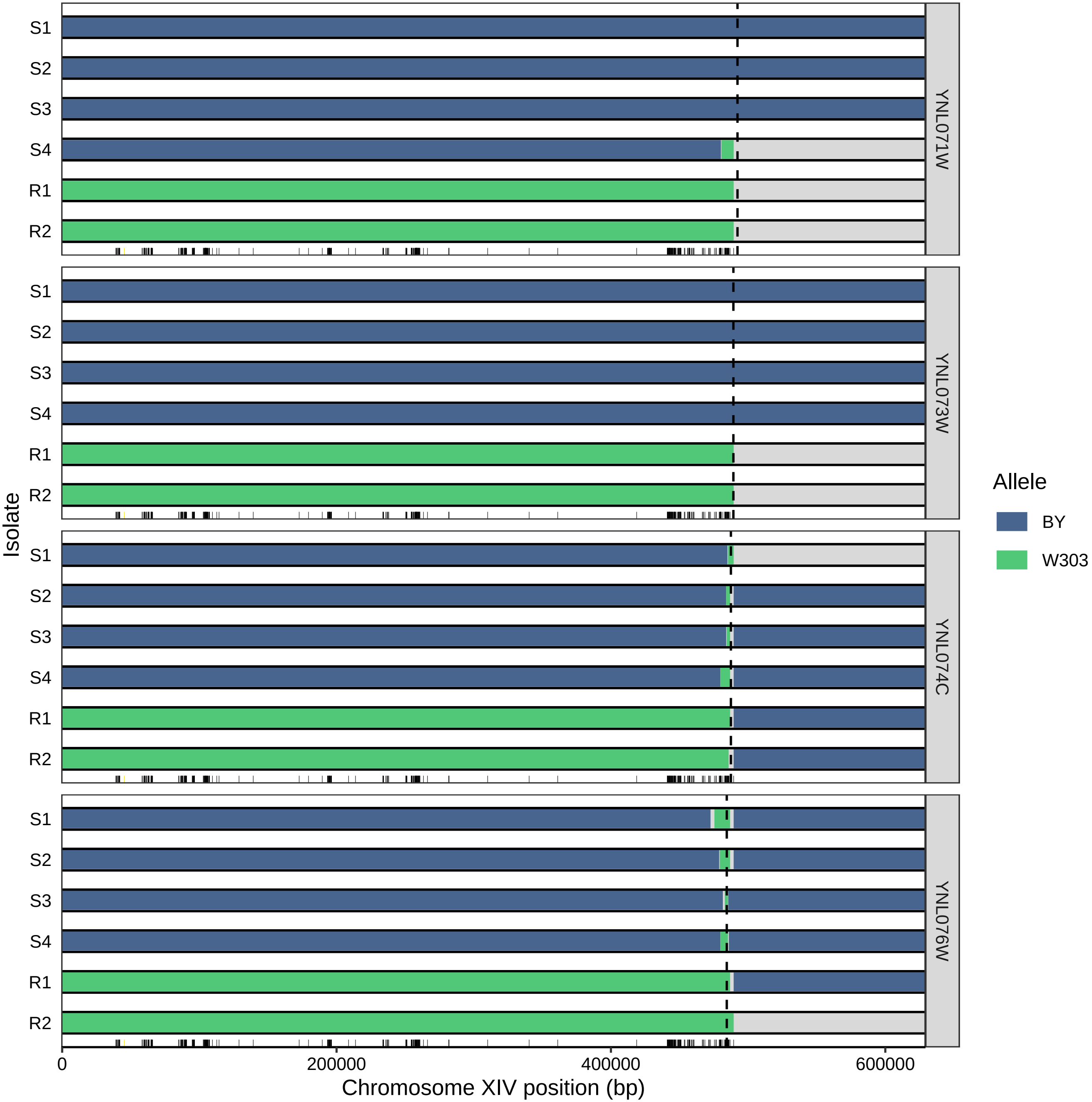

S1
S2
S3
S4
R1
R2
YNL071W
YNL073W
YNL074C
YNL076W
Isolate
Allele
BY
W303
0
200000
400000
600000
Chromosome XIV position (bp)

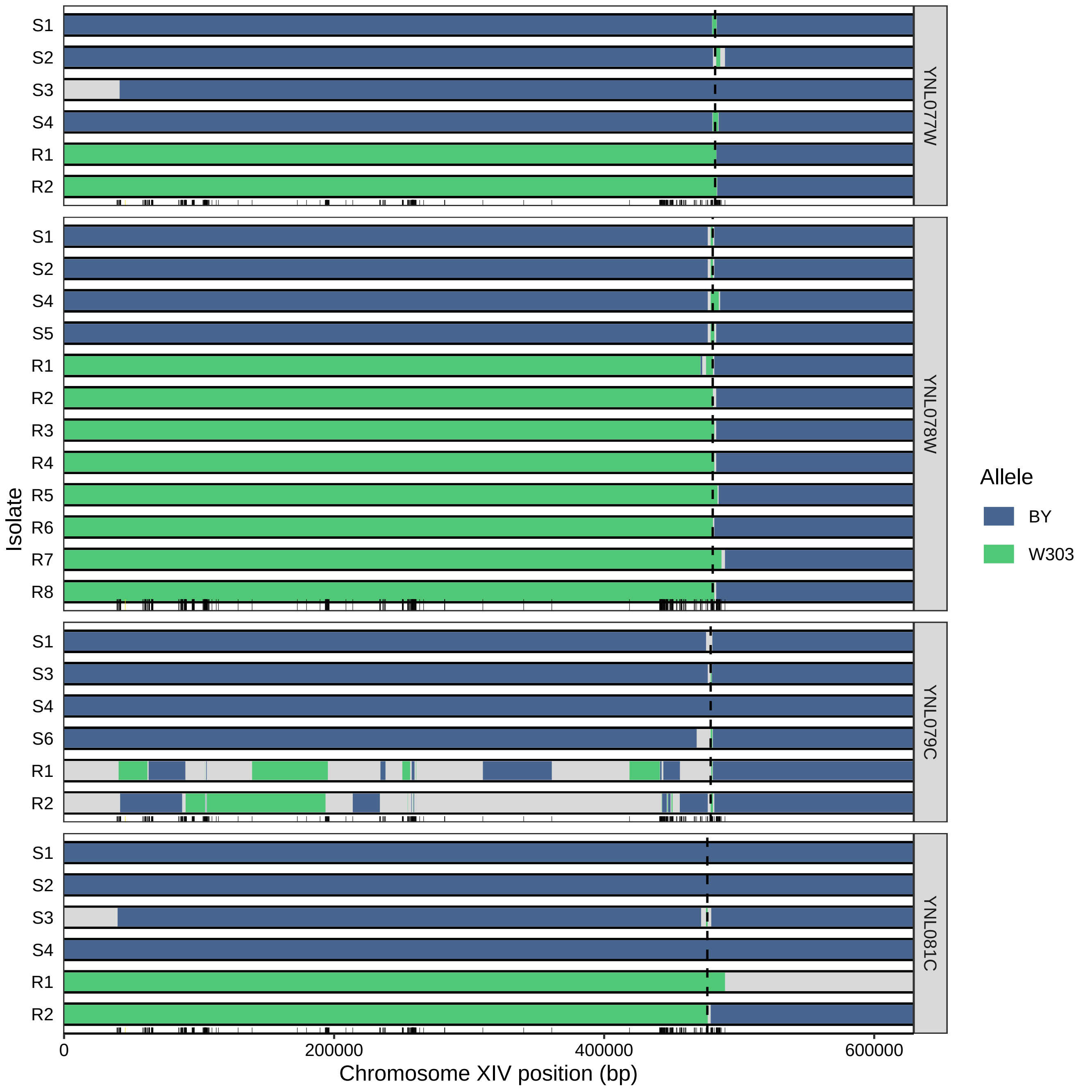

S1
S2
S3
S4
R1
R2
YNL077W
S1
S2
S4
S5
R1
R2
R3
R4
R5
R6
R7
R8
YNL078W
S1
S3
S4
S6
R1
R2
YNL079C
S1
S2
S3
S4
R1
R2
YNL081C
Isolate
0
200000
400000
600000
Chromosome XIV position (bp)
Allele
BY
W303

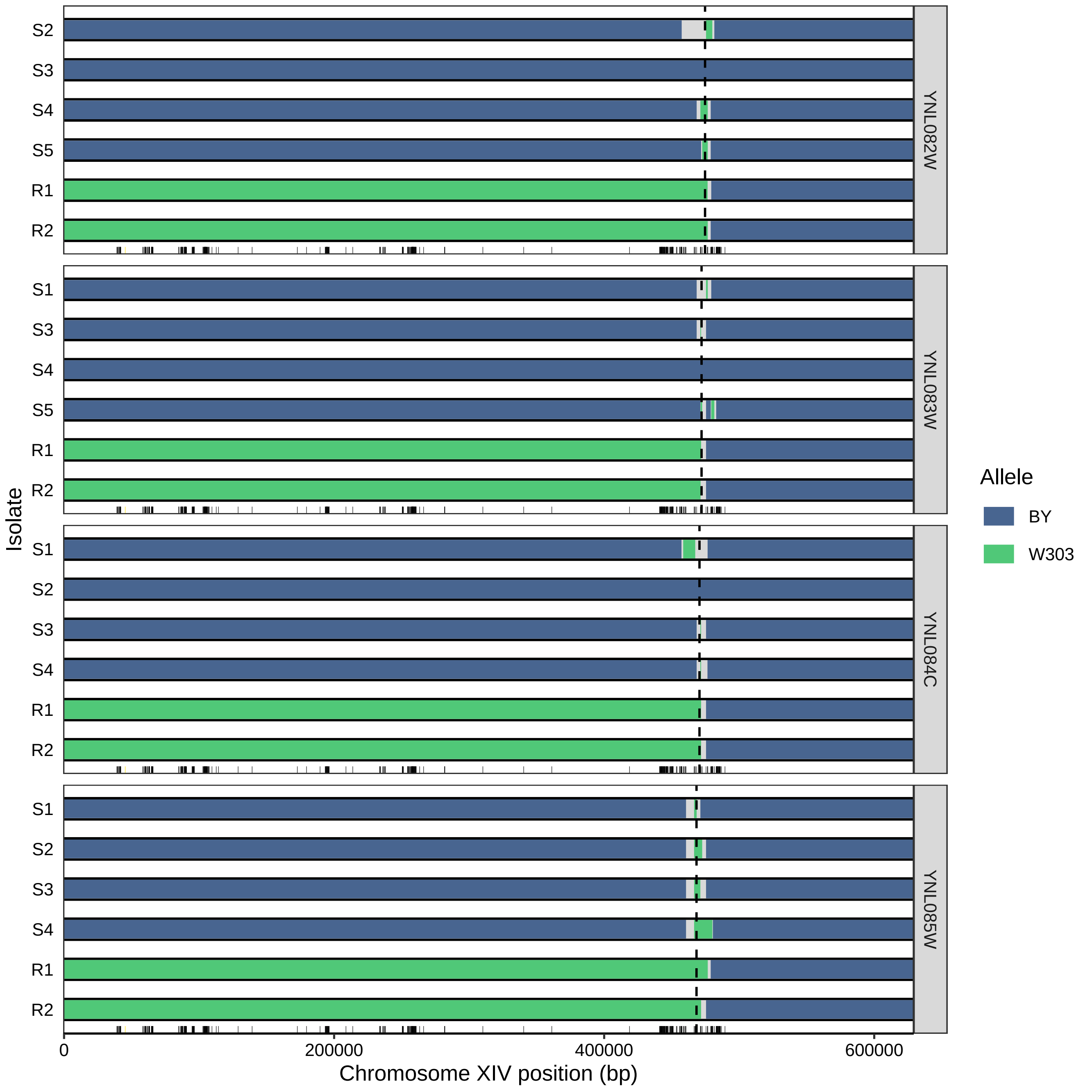

S2
S3
S4
S5
R1
R2
YNL082W
S1
S3
S4
S5
R1
R2
YNL083W
S1
S2
S3
S4
R1
R2
YNL084C
S1
S2
S3
S4
R1
R2
YNL085W
Isolate
0
200000
400000
600000
Chromosome XIV position (bp)
Allele
BY
W303

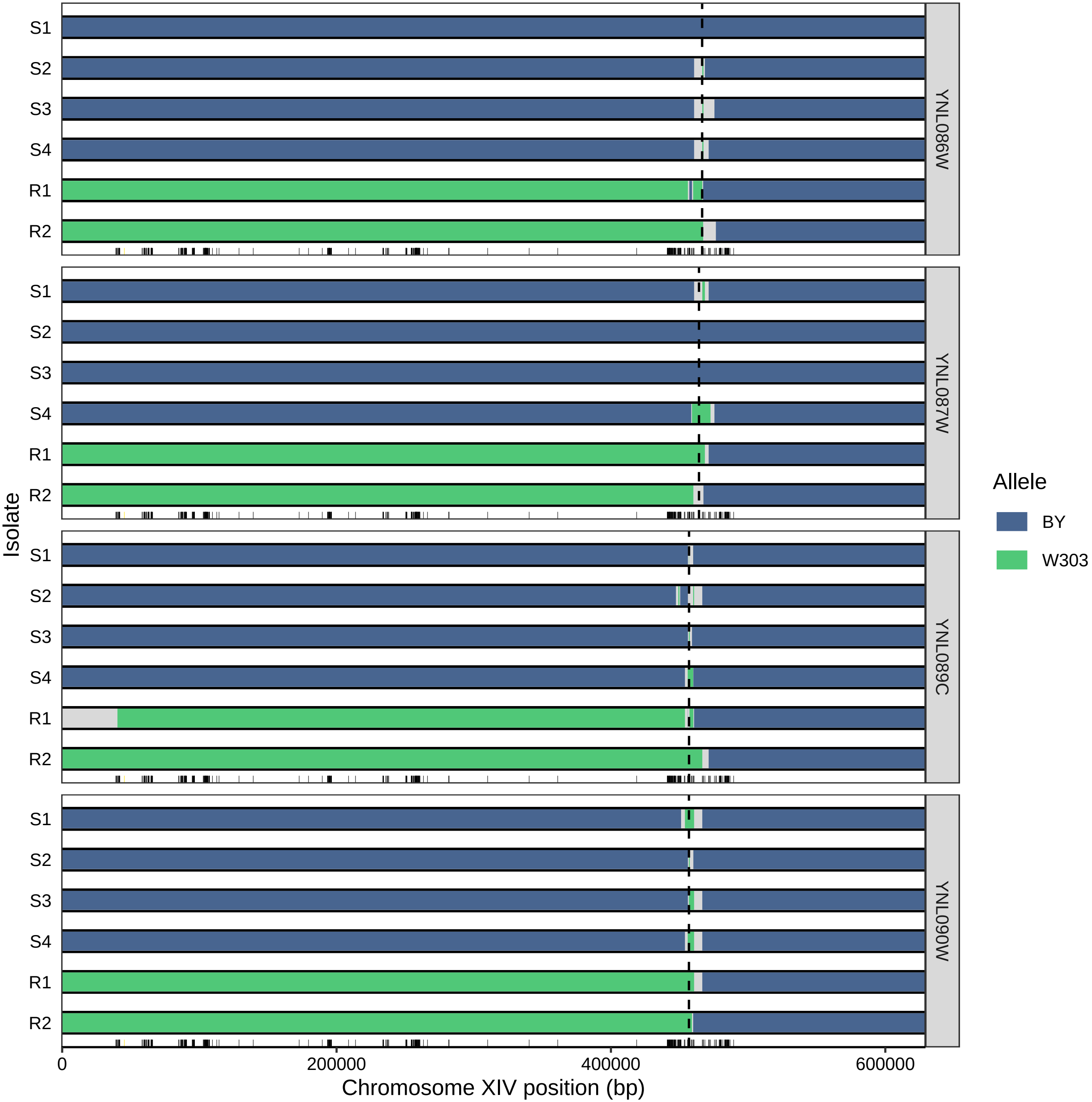
S1
S2
S3
S4
R1
R2
YNL086W
YNL087W
YNL089C
YNL090W
Isolate
0
200000
400000
600000
Chromosome XIV position (bp)
Allele
BY
W303

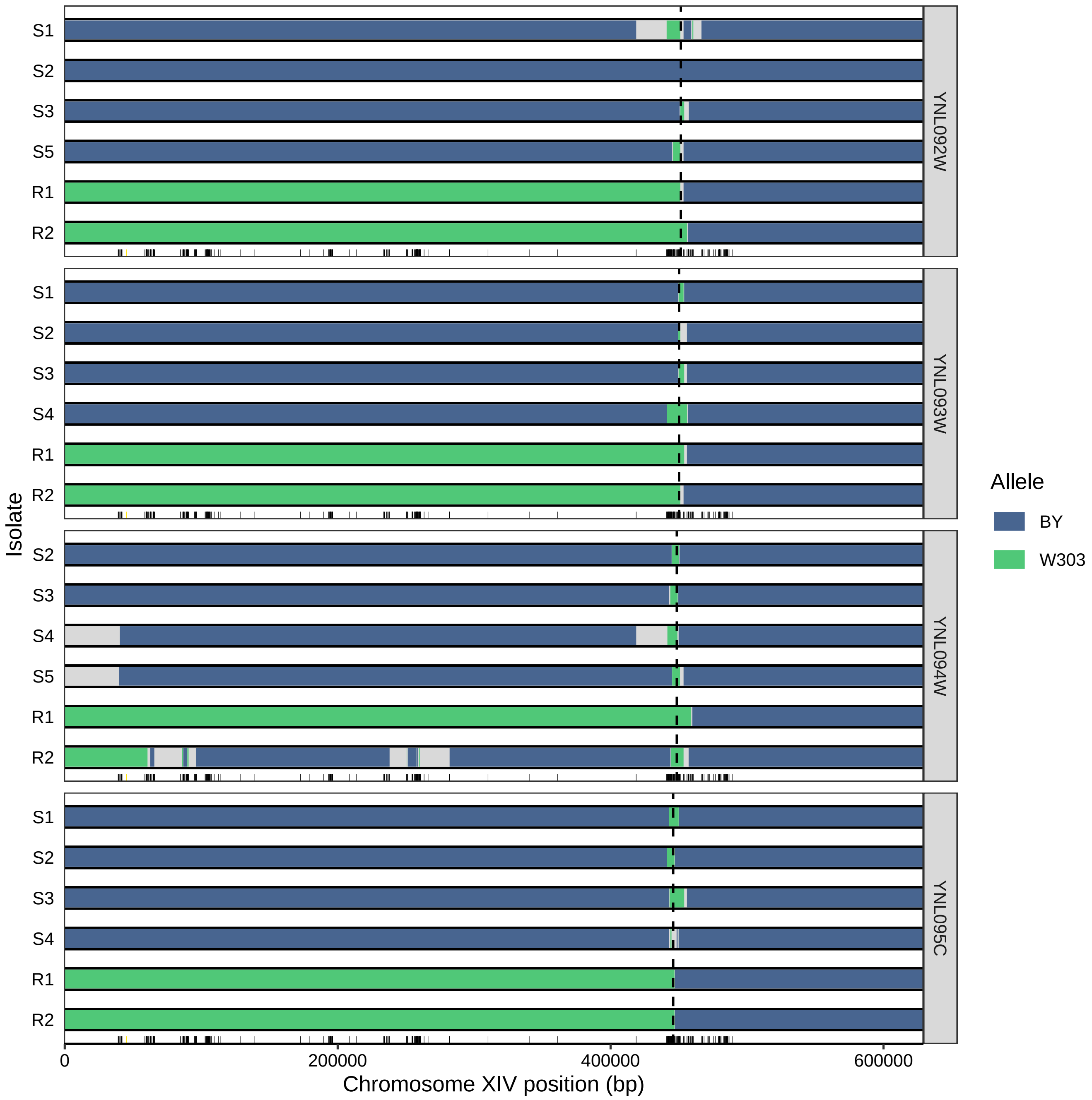

S1
S2
S3
S5
R1
R2
YNL092W
S1
S2
S3
S4
R1
R2
YNL093W
S2
S3
S4
S5
R1
R2
YNL094W
S1
S2
S3
S4
R1
R2
YNL095C
Isolate
0
200000
400000
600000
Chromosome XIV position (bp)
Allele
BY
W303

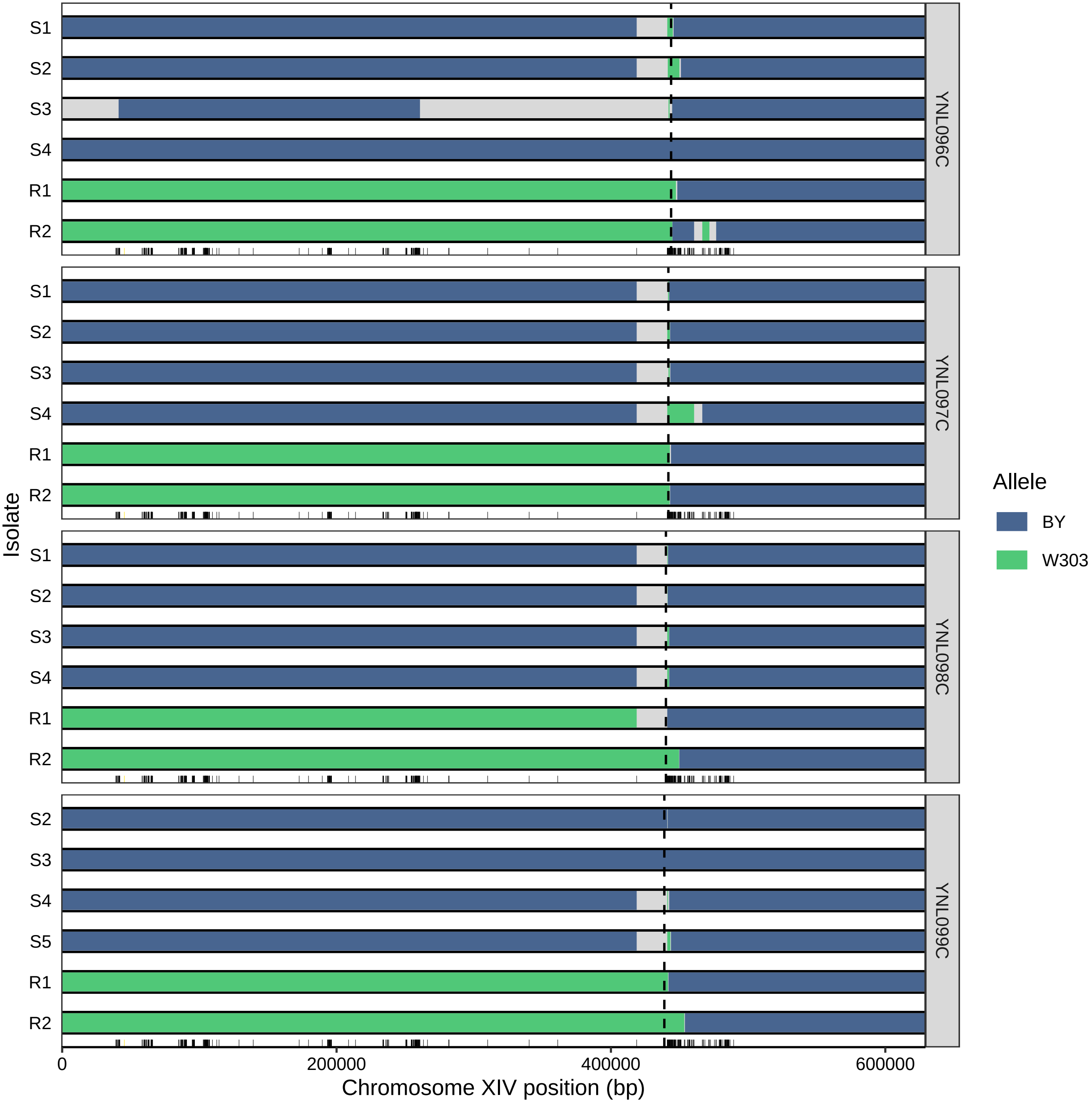
S1
S2
S3
S4
S5
R1
R2
YNL096C
YNL097C
YNL098C
YNL099C
Isolate
0
200000
400000
600000
Chromosome XIV position (bp)
Allele
BY
W303

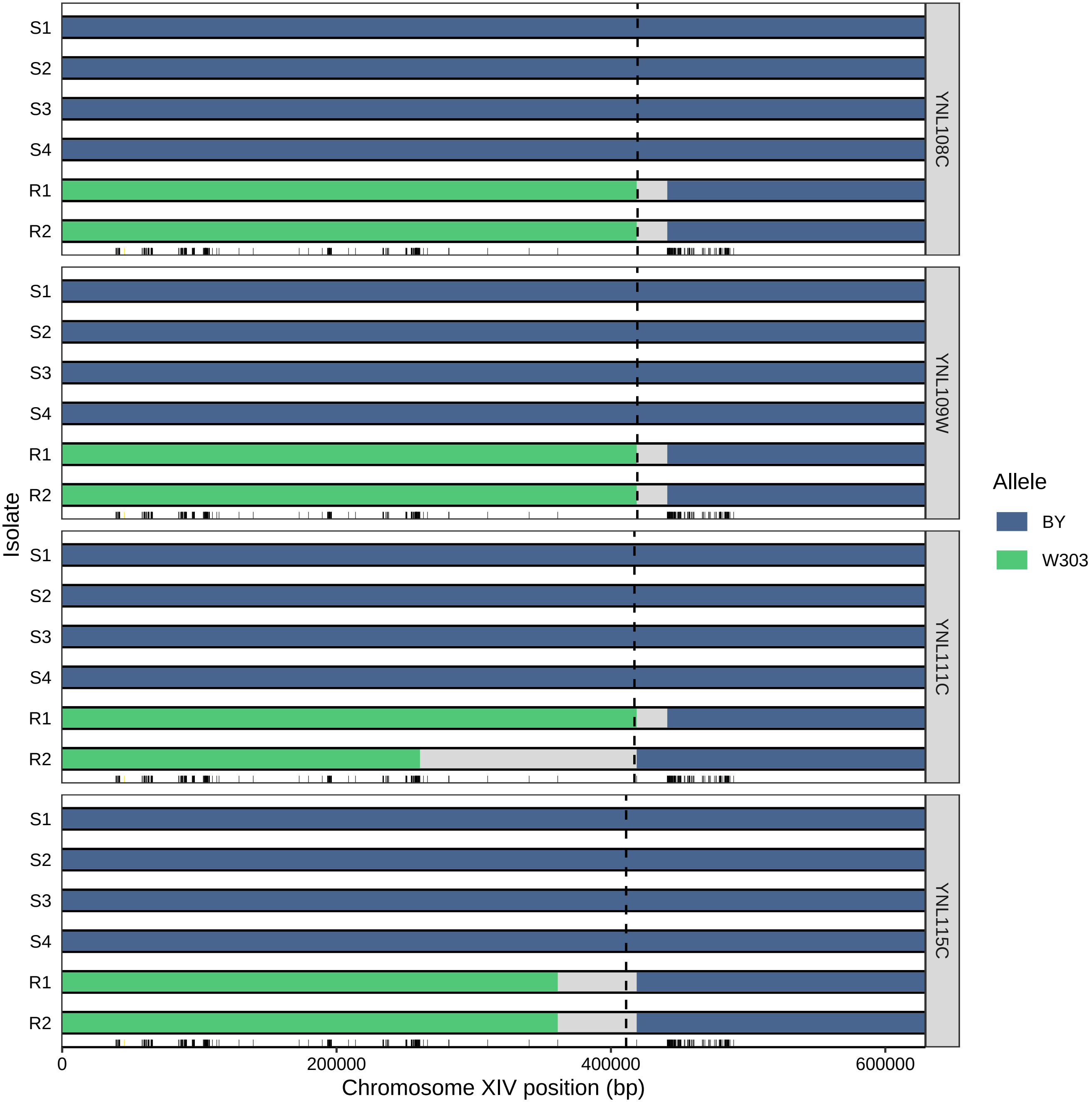

S1
S2
S3
S4
R1
R2
YNL108C
YNL109W
YNL111C
YNL115C
Isolate
0
200000
400000
600000
Chromosome XIV position (bp)
Allele
BY
W303

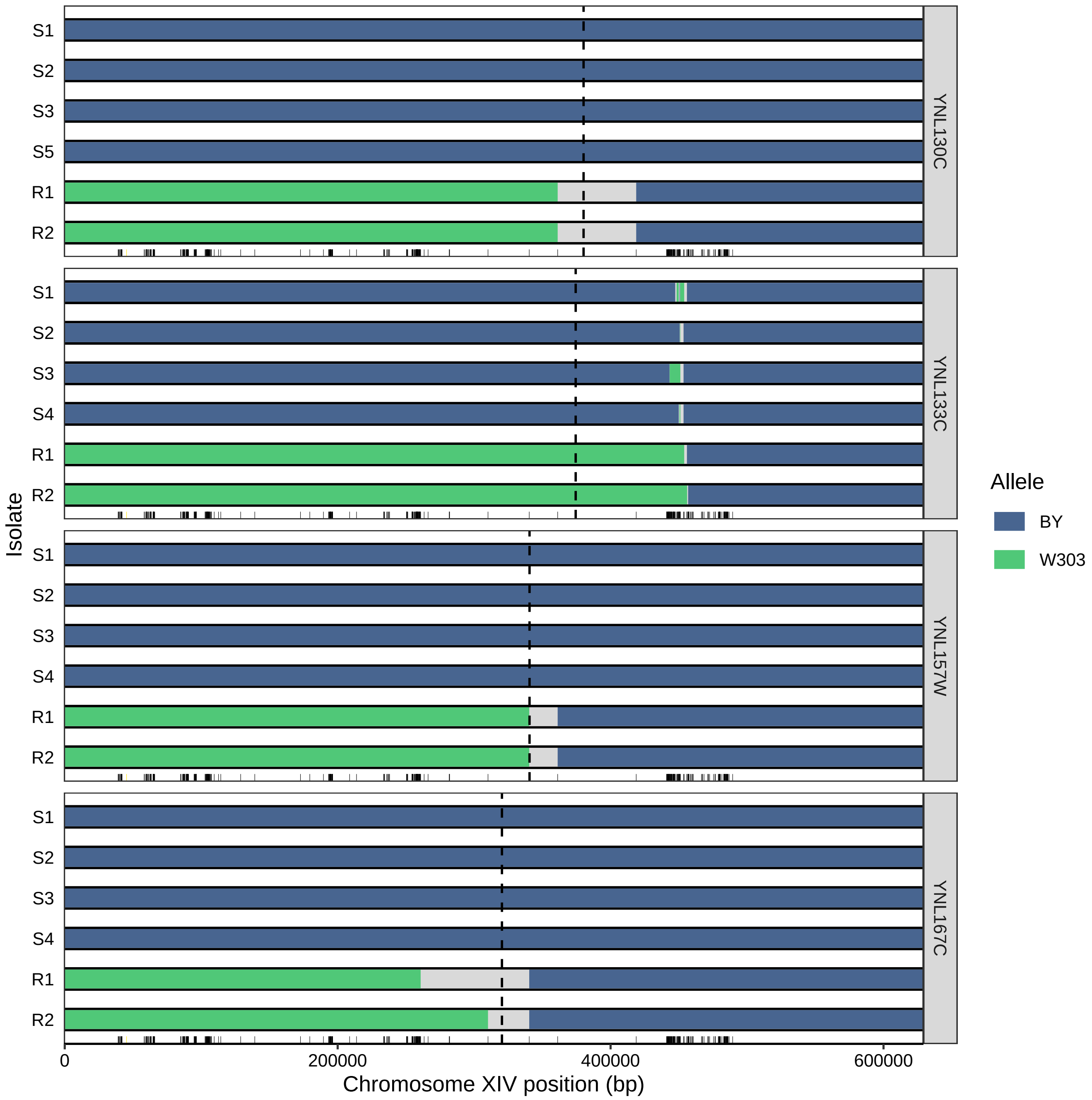

S1
S2
S3
S5
R1
R2
YNL130C
S1
S2
S3
S4
R1
R2
YNL133C
S1
S2
S3
S4
R1
R2
YNL157W
S1
S2
S3
S4
R1
R2
YNL167C
Isolate
0
200000
400000
600000
Chromosome XIV position (bp)
Allele
BY
W303

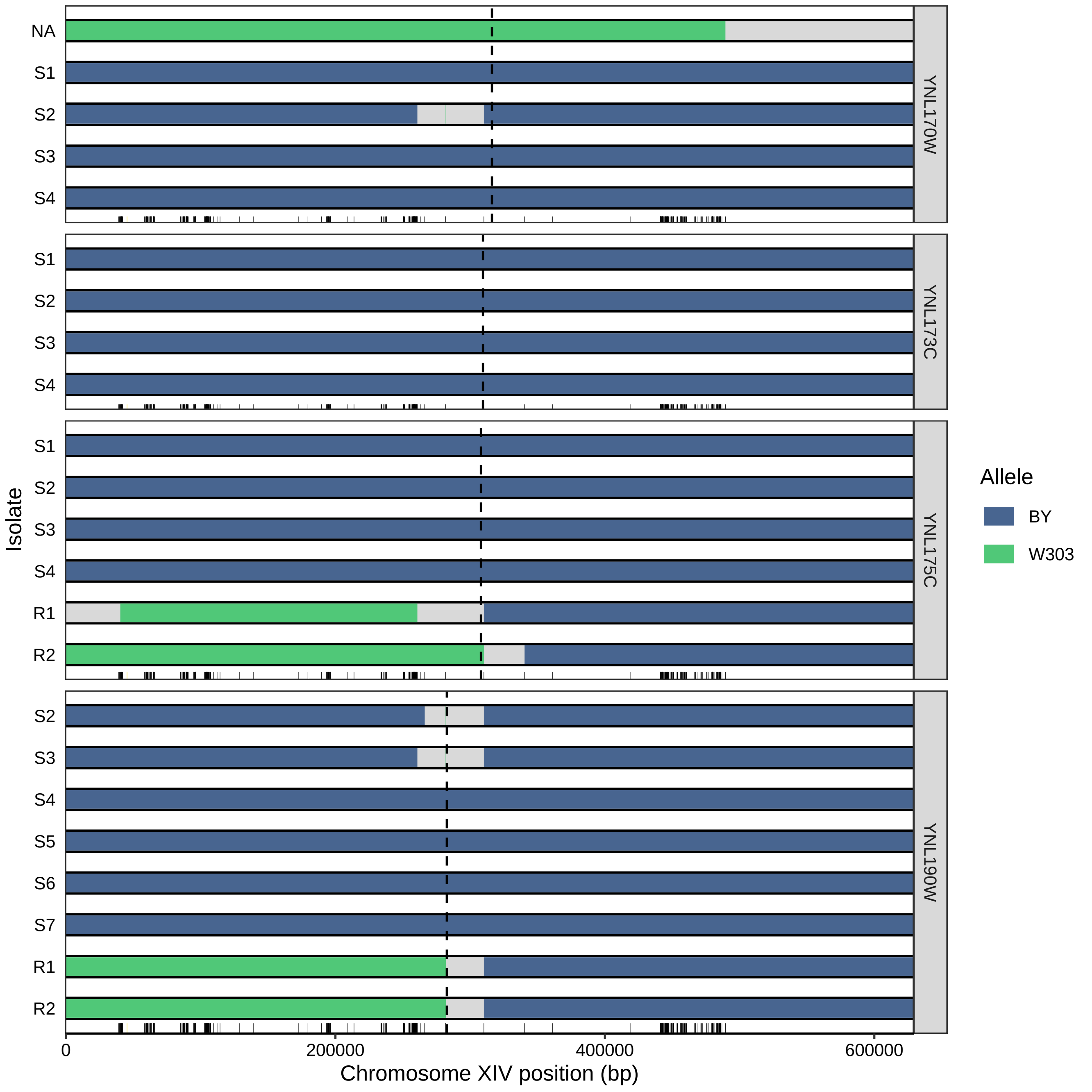

NA
S1
S2
S3
S4
YNL170W
S1
S2
S3
S4
YNL173C
S1
S2
S3
S4
R1
R2
YNL175C
S2
S3
S4
S5
S6
S7
R1
R2
YNL190W
Isolate
0
200000
400000
600000
Chromosome XIV position (bp)
Allele
BY
W303

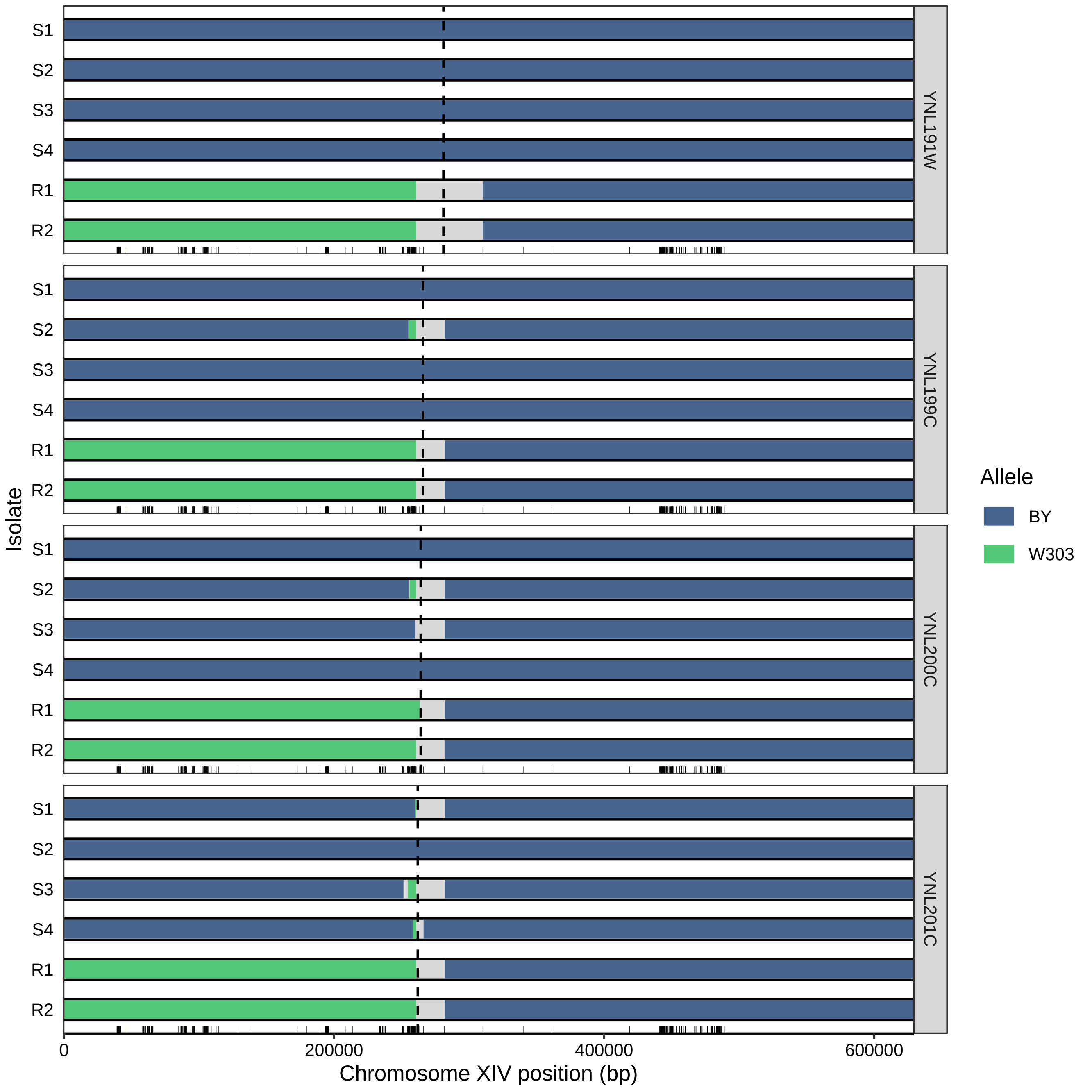

S1
S2
S3
S4
R1
R2
YNL191W
YNL199C
YNL200C
YNL201C
Isolate
Allele
BY
W303
0
200000
400000
600000
Chromosome XIV position (bp)

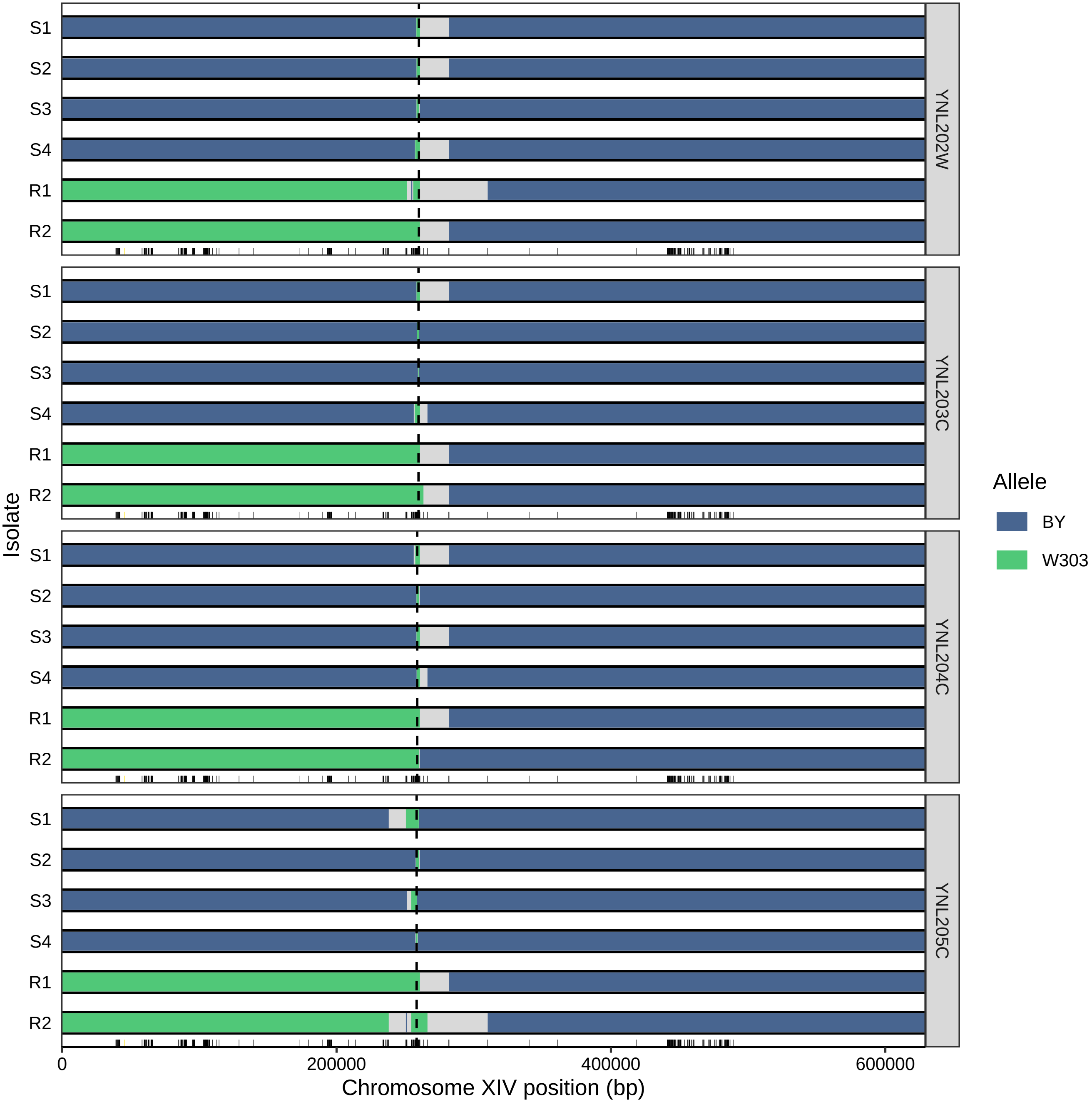

Isolate
S1
S2
S3
S4
R1
R2
YNL202W
YNL203C
YNL204C
YNL205C
0
200000
400000
600000
Chromosome XIV position (bp)
Allele
BY
W303

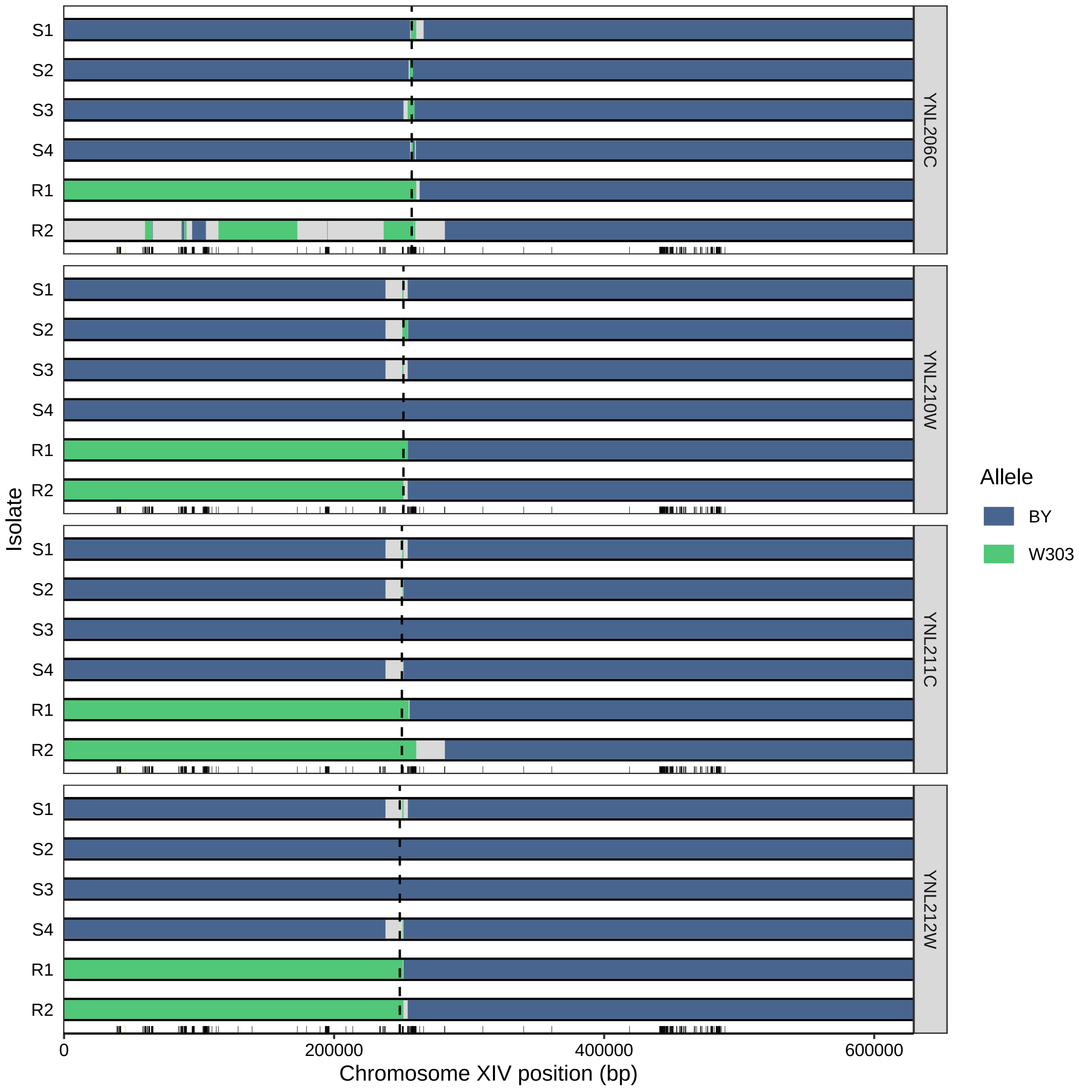

S1
S2
S3
S4
R1
R2
YNL206C
YNL210W
YNL211C
YNL212W
Isolate
0
200000
400000
600000
Chromosome XIV position (bp)
Allele
BY
W303

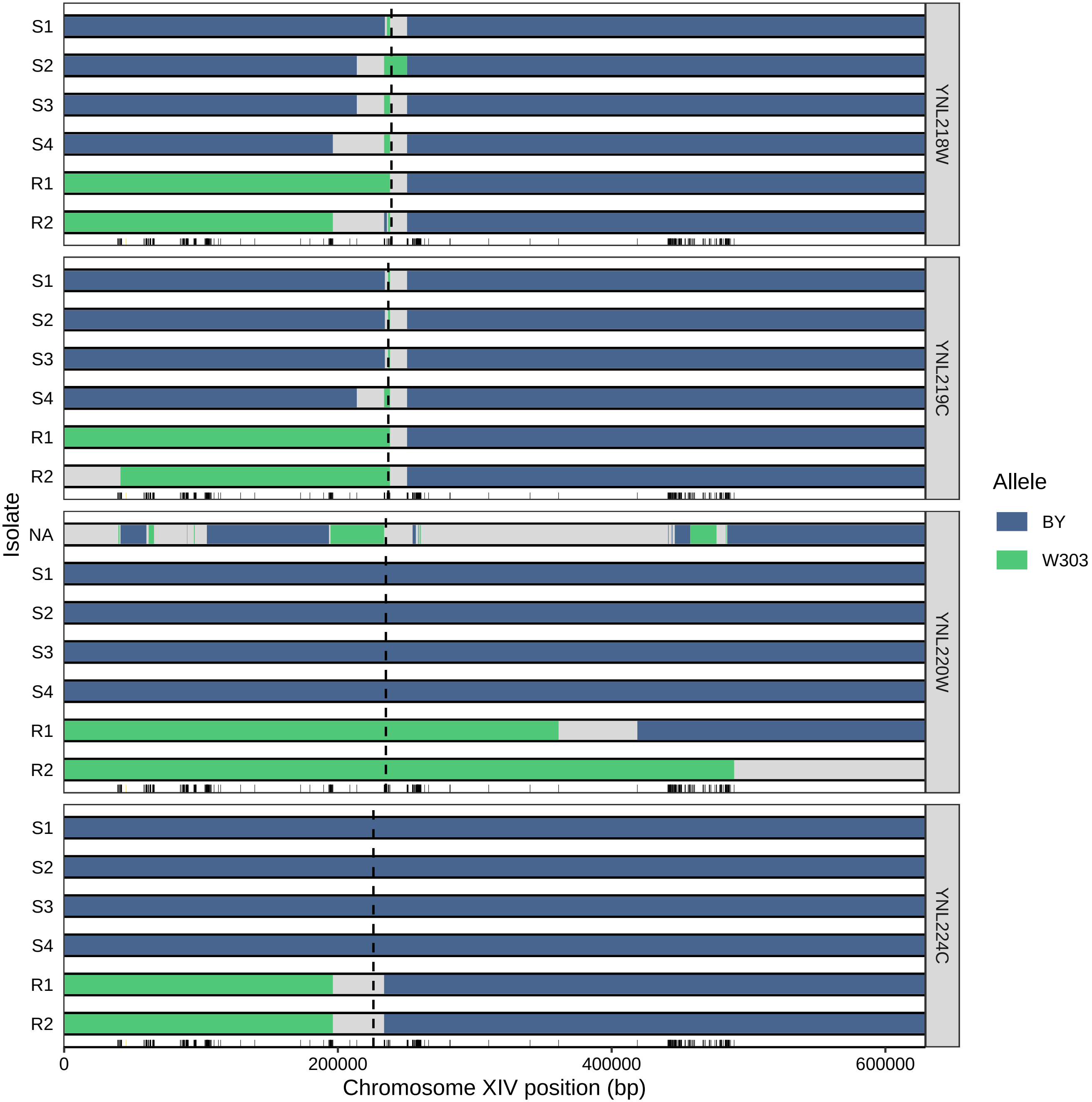

S1
S2
S3
S4
R1
R2
NA
YNL218W
YNL219C
YNL220W
YNL224C
Isolate
Allele
BY
W303
0
200000
400000
600000
Chromosome XIV position (bp)

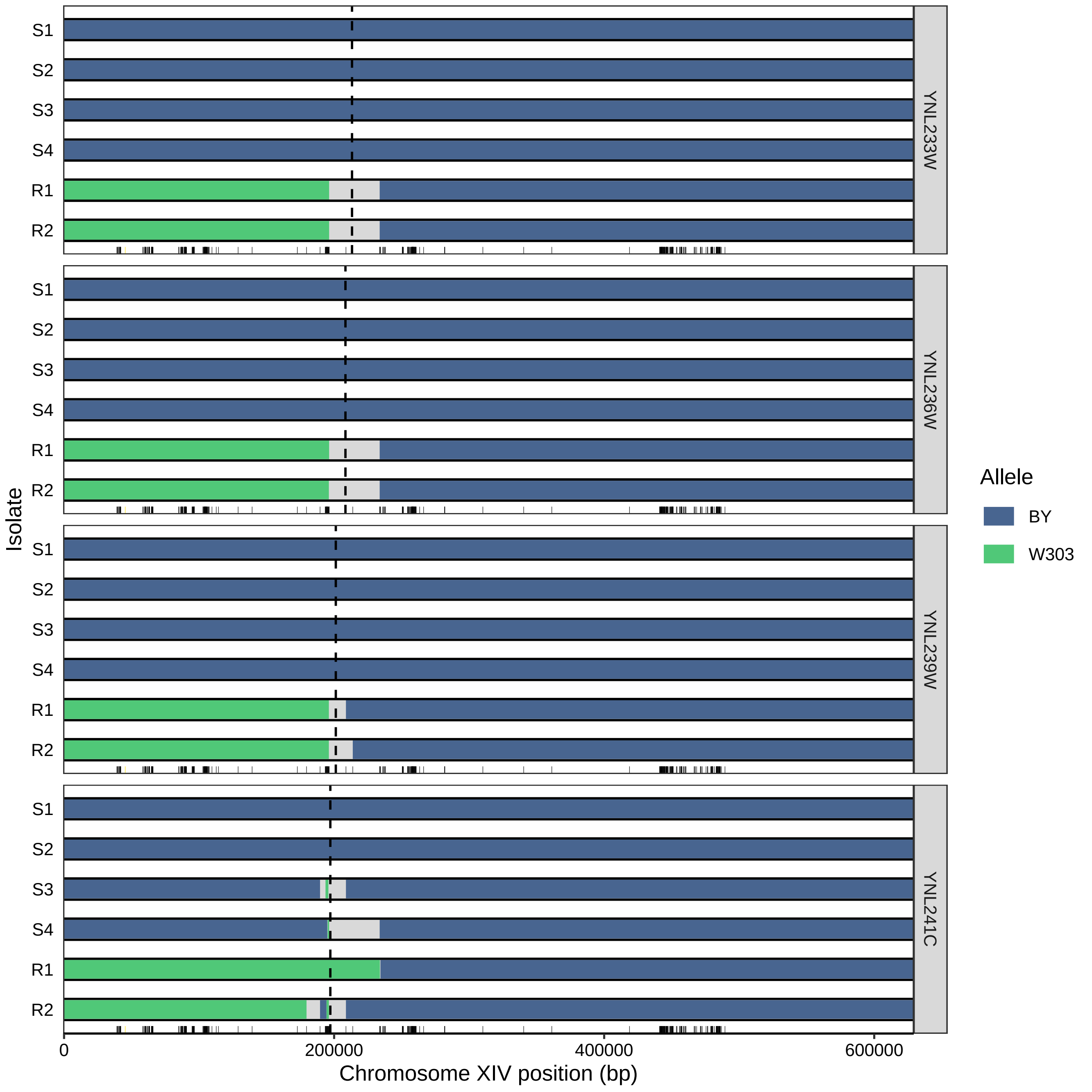

Isolate
S1
S2
S3
S4
R1
R2
YNL233W
YNL236W
YNL239W
YNL241C
0
200000
400000
600000
Chromosome XIV position (bp)
Allele
BY
W303

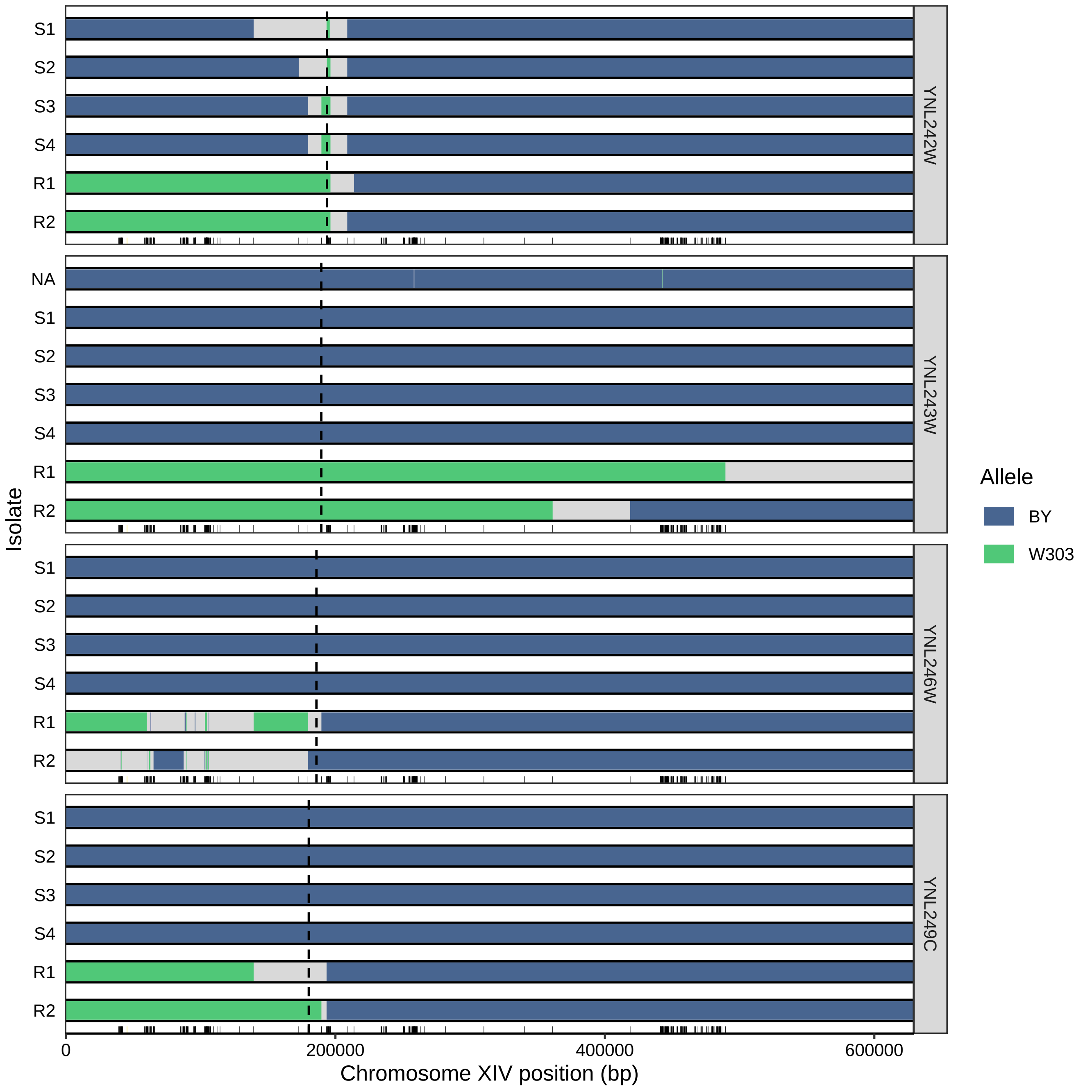

S1
S2
S3
S4
R1
R2
YNL242W
NA
S1
S2
S3
S4
R1
R2
YNL243W
S1
S2
S3
S4
R1
R2
YNL246W
S1
S2
S3
S4
R1
R2
YNL249C
Isolate
0
200000
400000
600000
Chromosome XIV position (bp)
Allele
BY
W303

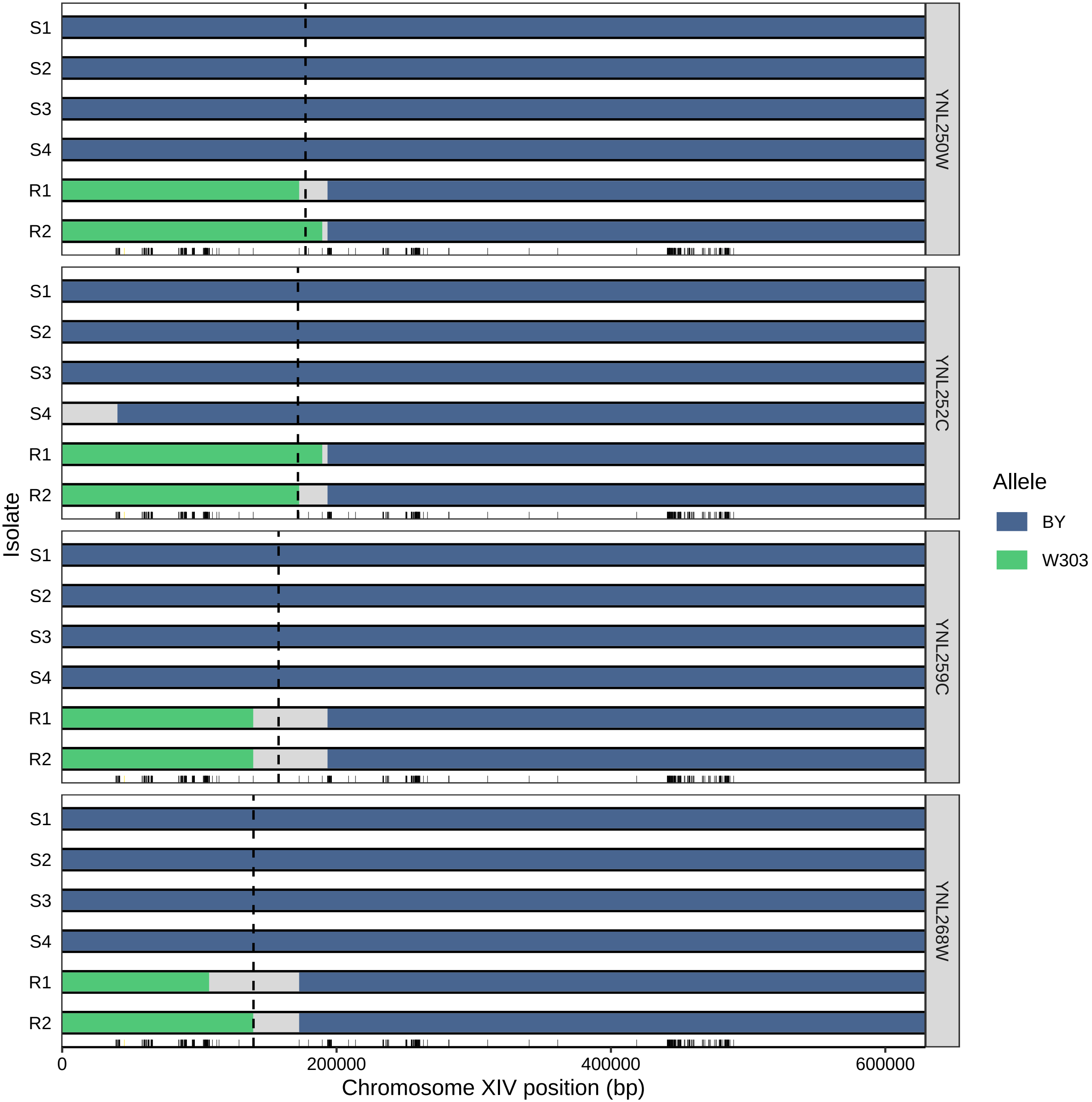

S1
S2
S3
S4
R1
R2
YNL250W
YNL252C
YNL259C
YNL268W
Isolate
0
200000
400000
600000
Chromosome XIV position (bp)
Allele
BY
W303

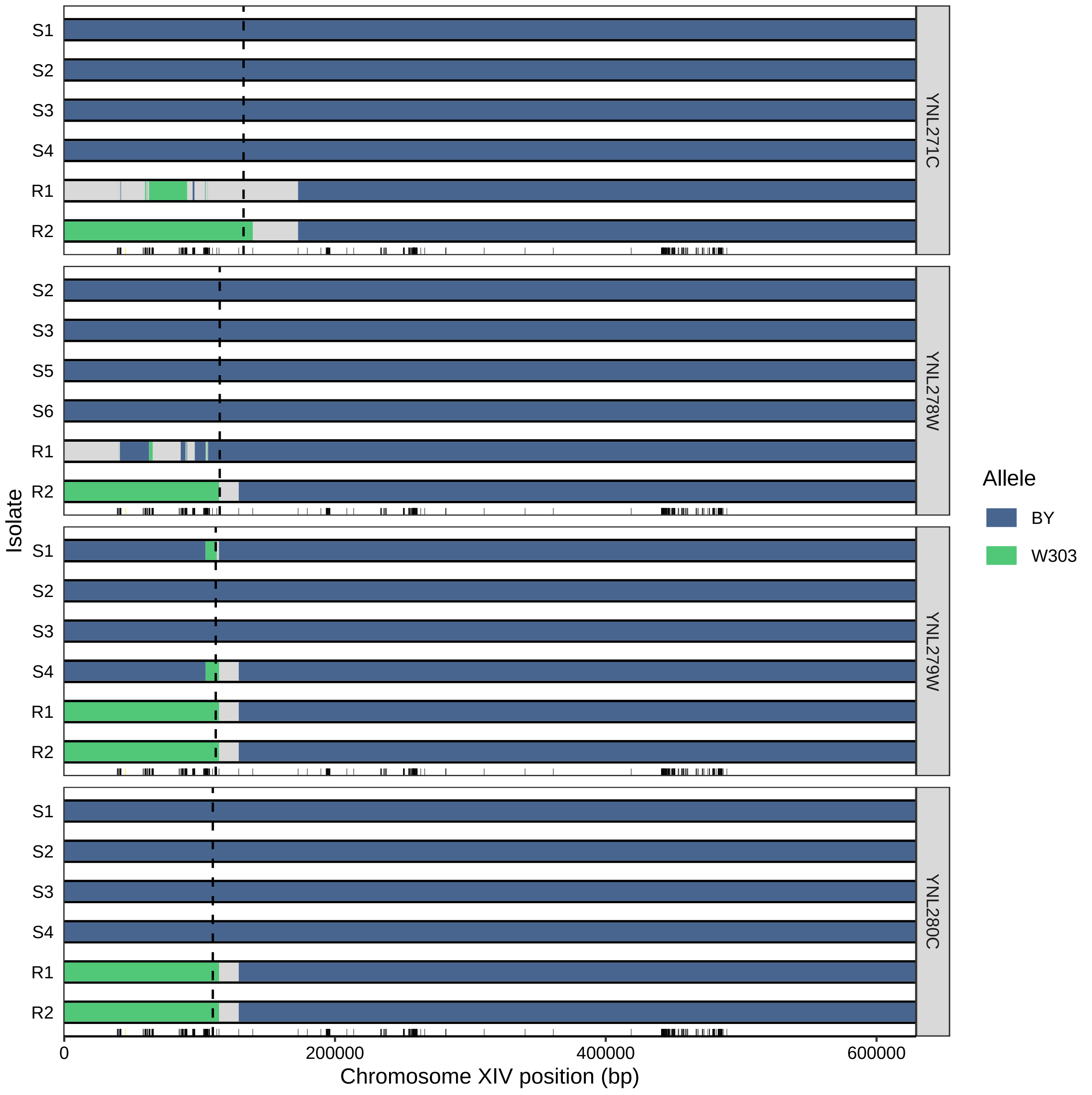
Isolate
S1
S2
S3
S4
R1
R2
S5
S6
YNL271C
YNL278W
YNL279W
YNL280C
0
200000
400000
600000
Chromosome XIV position (bp)
Allele
BY
W303

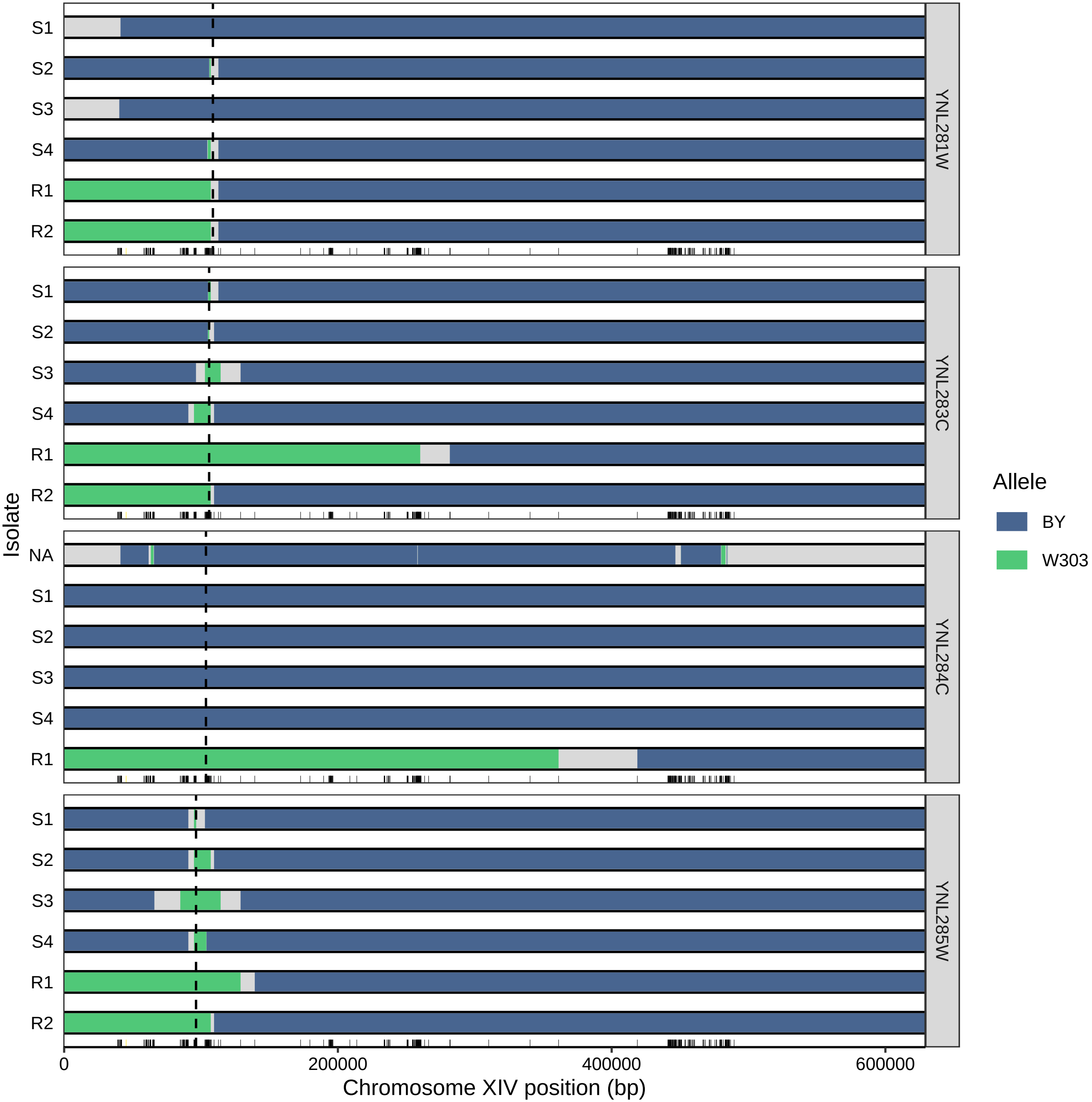

Isolate
S1
S2
S3
S4
R1
R2
NA
YNL281W
YNL283C
YNL284C
YNL285W
0
200000
400000
600000
Chromosome XIV position (bp)
Allele
BY
W303

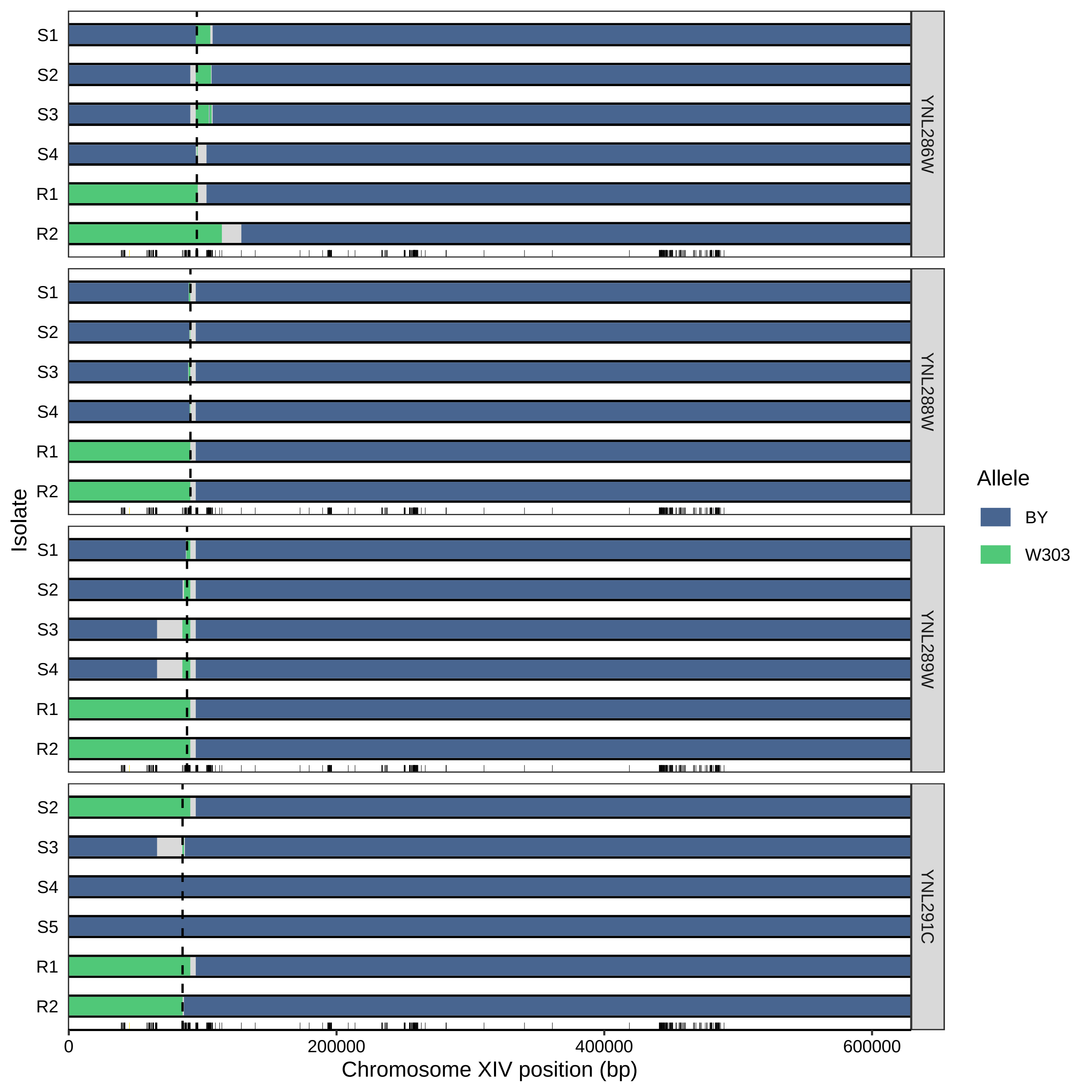

YNL286W
YNL288W
YNL289W
YNL291C
S1
S2
S3
S4
S5
R1
R2
Isolate
0
200000
400000
600000
Chromosome XIV position (bp)
Allele
BY
W303

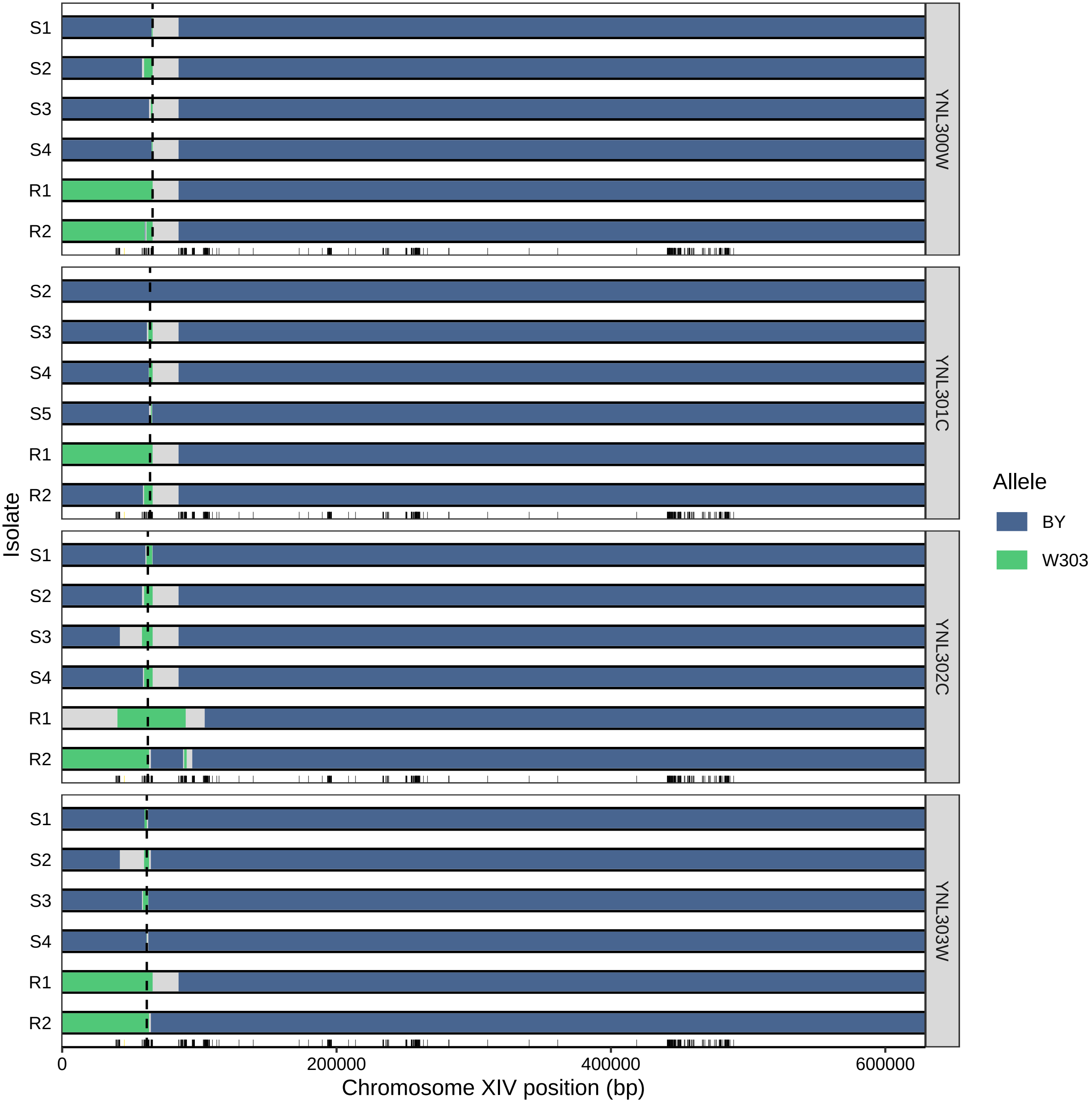

S1
S2
S3
S4
R1
R2
YNL300W
S2
S3
S4
S5
R1
R2
YNL301C
S1
S2
S3
S4
R1
R2
YNL302C
S1
S2
S3
S4
R1
R2
YNL303W
Isolate
0
200000
400000
600000
Chromosome XIV position (bp)
Allele
BY
W303

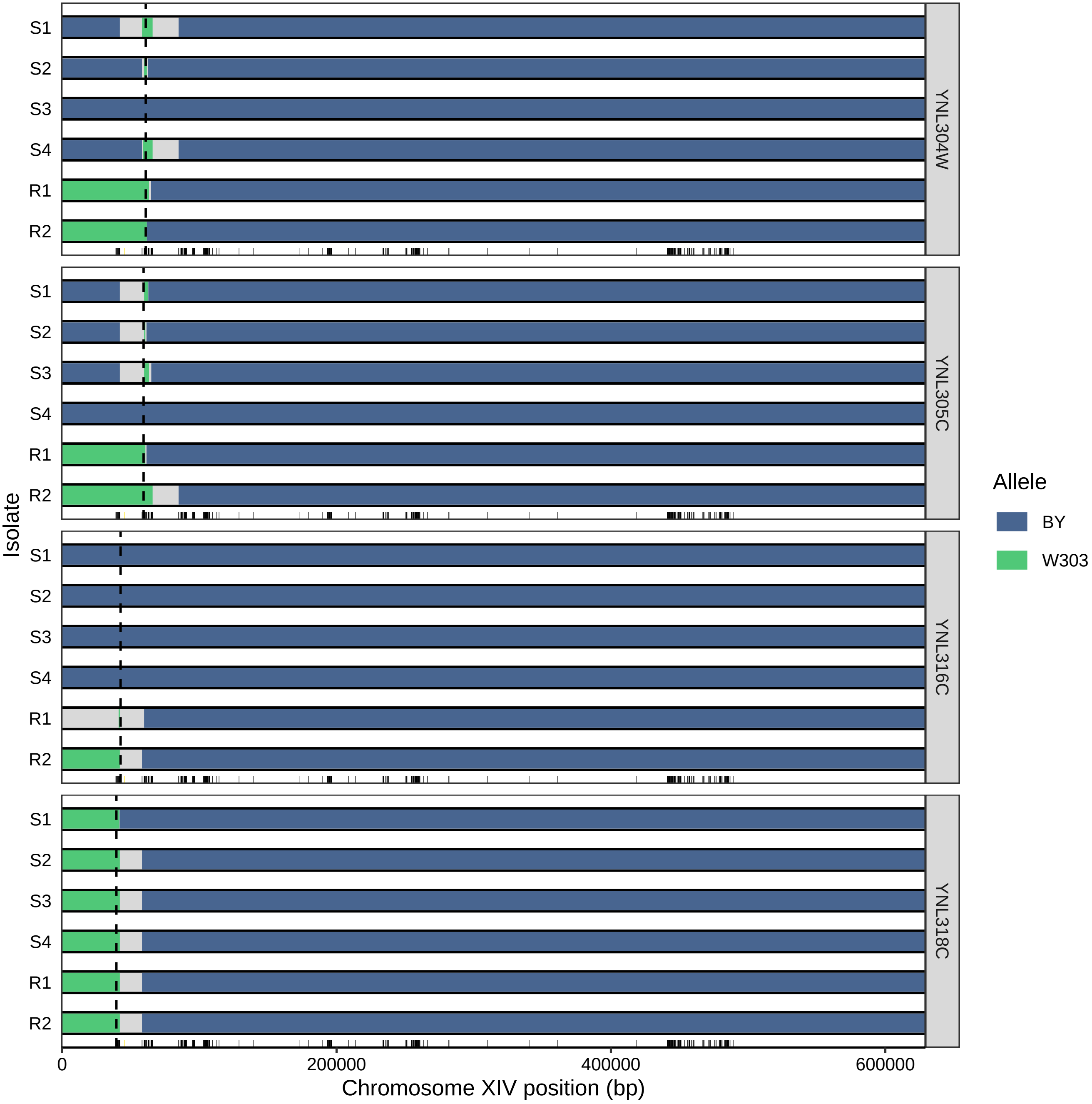

S1
S2
S3
S4
R1
R2
YNL304W
YNL305C
YNL316C
YNL318C
Isolate
Allele
BY
W303
0
200000
400000
600000
Chromosome XIV position (bp)
